# Defining the Active Conformation of Typical Protein Kinase Domains from Substrate-Bound PDB Structures Enables Active-State AlphaFold2 Models for All 437 Human Catalytic Protein Kinases

**DOI:** 10.64898/2026.02.19.706771

**Authors:** Joan Gizzio, Bulat Faezov, Qifang Xu, Roland L. Dunbrack

## Abstract

Humans have 437 catalytically competent protein kinase domains with the typical kinase fold, similar to the structure of Protein Kinase A (PKA). The active form of a kinase must satisfy requirements for binding ATP, magnesium, and substrate. From structural bioinformatics analysis of 248 crystal structures of 54 unique substrate-bound kinases, we derived structural criteria for the active form of typical protein kinases. We include well-known requirements on the DFG motif of the activation loop and the N-terminal domain salt bridge, but also on the positions of the N-terminal and C-terminal segments of the activation loop that must be placed appropriately to bind substrate. With these criteria, only 130 of the 437 human catalytic protein kinases (30%) are in the Protein Data Bank in their active form. Because the active forms of catalytic kinases are needed for understanding substrate specificity and the effects of mutations on catalytic activity in cancer and other diseases, we used AlphaFold2 to produce models of all 437 human protein kinases in the active form. This was accomplished with templates from the PDB that resemble substrate-bound structures, shallow multiple sequence alignments of orthologs and close paralogs of the query protein, and application of the active-kinase criteria to the output models. We selected models for each kinase based on intramolecular ipSAE scores of the activation loop residues of these models, demonstrating that the highest scoring models have the lowest or close to the lowest RMSD to 29 non-redundant substrate-bound structures in the PDB. A larger benchmark of 117 active kinase structures with solved activation loops in the PDB shows that 71% of the highest scoring AlphaFold2 models had backbone RMSD < 1.0 Å to the benchmark structures and 92% were within 2.0 Å. Models for all 437 catalytic kinases are available at https://dunbrack.fccc.edu/kincore/activemodels. We believe they may be useful for interpreting mutations leading to constitutive catalytic activity in cancer as well as for templates for modeling substrate and inhibitor binding for molecules which bind to the active state.

## INTRODUCTION

Protein kinases regulate most cellular processes in eukaryotes. In humans, their dysregulation is often involved in disease and they are therefore often targets in drug development, especially in cancer [1]. A large majority of human protein kinases take on a common fold first determined by Susan Taylor and colleagues in 1991 [2], consisting of an N-terminal domain of five beta strands and the C-helix and a largely helical C-terminal domain. The residues involved in catalytic activity are contained in the catalytic and activation loops that form a pocket for ATP binding and a binding site for substrates. Humans have 480 genes which contain at least one full-length typical protein kinase domain; 13 of these have two kinase domains, for a total of 493 typical kinase domains [3]. Of these 493 kinase domains, 437 domains from 429 genes are likely catalytic kinases, participating in phosphorylation of Ser, Thr, or Tyr residues on proteins, and 56 are likely pseudokinases. As of December 1, 2025, in the PDB there are structures for 305 human typical kinase domains, of which 277 are catalytic kinases and 28 are pseudokinases [4]. Active and inactive conformations of typical kinases have been classified in several ways [5–12].

The active form is generally very similar across kinases because of the requirements of binding ATP, magnesium ions, and substrate, which impart constraints on the conformation of the activation loop [13]. Early in the history of structure determination of kinases, a classification of structures into “DFGin” and “DFGout” was described [14]. In DFGin structures, the Asp side chain of the DFG motif is “in” the ATP binding site, and the Phe side chain of the DFG motif is in a pocket adjacent to the C-helix of the N-terminal domain. In DFGout structures, the Asp side chain is “out” of the active site, and the Phe side chain is removed from the C-helix pocket, allowing for the binding of Type 2 inhibitors such as imatinib that span both the ATP site and the C-helix pocket [15].

We previously used the presence of bound ATP, magnesium ion, and a phosphorylated activation loop to identify a set of 24 “catalytically primed” structures of 12 different kinases in the PDB [10]. We found that in addition to being “DFGin,” these structures possess specific backbone and side-chain dihedral angles for the DFG motif (“*BLAminus*”), comprising the backbone dihedral angles of the residue immediately preceding DFG as well as the D and F residues, and the side-chain χ_1_ dihedral angle of the DFG-Phe residue. They also possess a previously characterized salt bridge between a conserved glutamic acid residue in the C-helix and a conserved lysine residue in beta strand 3 of the N-terminal domain [16]. These structures are often referred to as “C-helix-in.” Using these basic criteria for active kinases, only 195 of 437 catalytic typical human kinase domains (45%) are currently represented in the PDB with active structures.

Previous efforts [5–12] have not considered whether an observed experimental structure can bind substrate, which may require additional structural features [13, 17]. In this paper, we present a structural bioinformatics analysis of 54 non-redundant substrate-kinase complexes from the PDB to define criteria for identifying catalytically active protein kinases structures, including the ability to bind substrate. In addition to the dihedral angle criteria on the X-DFG motif and a distance criterion on the N-terminal domain salt bridge, we impose distance and dihedral angle criteria on the N and C terminal halves of the activation loop, whose positions are necessary for the formation of a substrate binding site. With these criteria, the number of human kinases with active structures is limited to 130 out of 437 catalytic protein kinases or 30%. Only 116 human catalytic protein kinases (26%) possess active structures and complete coordinates for the activation loop in the PDB.

The program AlphaFold2 from DeepMind is a deep-learning program for highly accurate protein structure prediction that is trained on structures from the PDB [18]. It takes as input the query sequence, a multiple sequence alignment (MSA) of homologs of the query, and optionally template structures related to the query. DeepMind has provided AlphaFold2 models of nearly all human proteins found in UniProt, which are available on a website provided by the European Bioinformatics Institute [19]. These were generated from a single structure produced by one seed of model 1.1.1 described in Jumper et al. [18]. However, with the criteria established in this paper, only 240 of the 437 (55%) catalytic human protein kinases have a fully active model in the EBI data set.

Because of the importance of knowing the active-state structures of kinases for understanding such features as substrate recognition, the effect of activating mutations in cancer, and drug development, we present a pipeline for producing active models of typical protein kinases using the program AlphaFold2. Several groups have found that using MSAs of reduced depth and templates in specific conformational states coerces AF2 to produce conformationally variable models, including some models in the conformational state of the templates [20, 21]. We use similar techniques to compute predicted structures of active kinases, using templates identified as active by our new criteria for active kinase structures, MSAs derived from orthologs of each kinase (with less than 90% sequence identity to each other), and applying the same criteria to the output models. From the active output models, we choose the top model using a modified intramolecular ipSAE score [22] based on aligned residues in the kinase domain and scored residues in the activation loops.

We benchmark our protocol with 29 substrate-bound kinase structures in the PDB with complete activation loops and a set of 117 kinase structures from the PDB that satisfy our active criteria and have complete (or nearly complete) activation loop coordinates around the substrate binding site. We show that the ipSAE scores for the activation loop are inversely correlated with RMSD of the activation loop for well characterized kinases. With these methods, we have produced active models of all 437 catalytic human protein kinase domains and made these models available on our Kinase Conformational Resource website and database, KinCore (https://dunbrack.fccc.edu/kincore/activemodels).

## RESULTS

### Catalytic protein kinases We previously published an alignment of all 497 human kinase domains from 484 genes annotated by Uniprot [3]. This list excluded atypical kinases, such as ADCK, PI3/PI4, Alpha, FAST, and

RIO family kinases (https://www.uniprot.org/docs/pkinfam.txt). Since that time, three kinase genes have been identified as likely pseudogenes (SIK1B, PDPK2P, and PRKY) [23]. Omitting these pseudogenes and the truncated protein kinase domain in PLK5 (65 amino acids), we are left with 480 genes containing 493 typical protein kinase domains in the human proteome.

To make catalytically active models of all human kinases with typical protein kinase domains, we need to distinguish between catalytic protein kinase domains and non-catalytic protein kinase domains or pseudokinases. *Catalytic* protein kinase domains are those able to phosphorylate proteins on Ser, Thr, or Tyr residues. *Non-catalytic* protein kinase domains or pseudokinases are domains that possess the typical protein kinase fold but lack protein kinase activity, although they may have other catalytic activity (e.g., POMK). We identified catalytic protein kinases based on the presence of the Asp residue in the HRD motif, the Asp residue in the DFG motif, and the Lys residue of the N-terminal domain salt bridge. We reclassified WNK (“With No Lysine”) kinases as catalytic protein kinases. Several kinases were reclassified based on literature annotations (e.g., BUB1B [24] and RYK are pseudokinases [25]). The result of these efforts was a list of 437 active kinase domains in 429 genes. Eight of these 429 genes have two (likely) catalytic protein kinase domains: RPS6KA1, RPS6KA2, RPS6KA3, RPS6KA4, RPS6KA5, RPS6KA6, OBSCN, and SPEG. Our previous phylogenetic analysis classified these 437 active domains into families as follows: AGC (60 kinases), CAMK (83), CK1 (11), CMGC (65), NEK (11), OTHER (43), STE (45), TKL (37), and TYR (82). On our Kincore website (https://dunbrack.fccc.edu/kincore) and in this paper, we use the family name as a prefix in front of the HUGO gene name (e.g., TYR_BTK) [26]. The catalytic protein kinase domains and associated data are listed in **Supplementary Table 1**. The pseudokinase domains are listed in **Supplementary Table 2**.

### The characteristics of active protein kinase domains derived from substrate-bound structures

To identify structural features of the active form of catalytic protein kinases, we compiled a set of structures that constitute likely catalytically active structures of protein kinases. It consists of 248 structures of kinases in the August 2024 Protein Data Bank (PDB) with peptide or protein substrates bound at the active site (**Supplementary Table S3**); the set contains 54 unique kinase/substrate combinations (**Table 1**). The example shown in **Figure 1** is an active form of human AKT1 bound to a peptide substrate from GSK3B, PDB:4ekk [27]. Some of the “substrates” are in fact substrate-mimicking inhibitors, which bind similarly to substrates. Kinases in this table are represented more than once if: 1) they contain different bound substrates in the active site; 2) if the same substrate protein has multiple different constructs (e.g., peptide fragment vs full-length); or 3) if the same substrate protein is bound at different phosphorylation sites. There are twenty unique complexes with folded proteins domains, and seventeen of these are autophosphorylation complexes (marked “A” in column 5 of **Table 1**). Autophosphorylation complexes are homodimers of protein kinases inside crystals (or within cryo-EM structures) in which a known trans-autophosphorylation site of one monomer (Ser, Thr, or Tyr) is sitting in the active site of another monomer in the manner of a kinase substrate (i.e., similar to known peptide substrate complexes). We identified these as we did previously [28] by building out a 3×3×3 group of unit cells and searching for hydroxyl OH atoms of Ser, Thr, and Tyr of one monomer in hydrogen bonding distance of the aspartic acid side chain of the HRD motif of another monomer.

**Figure 1.**
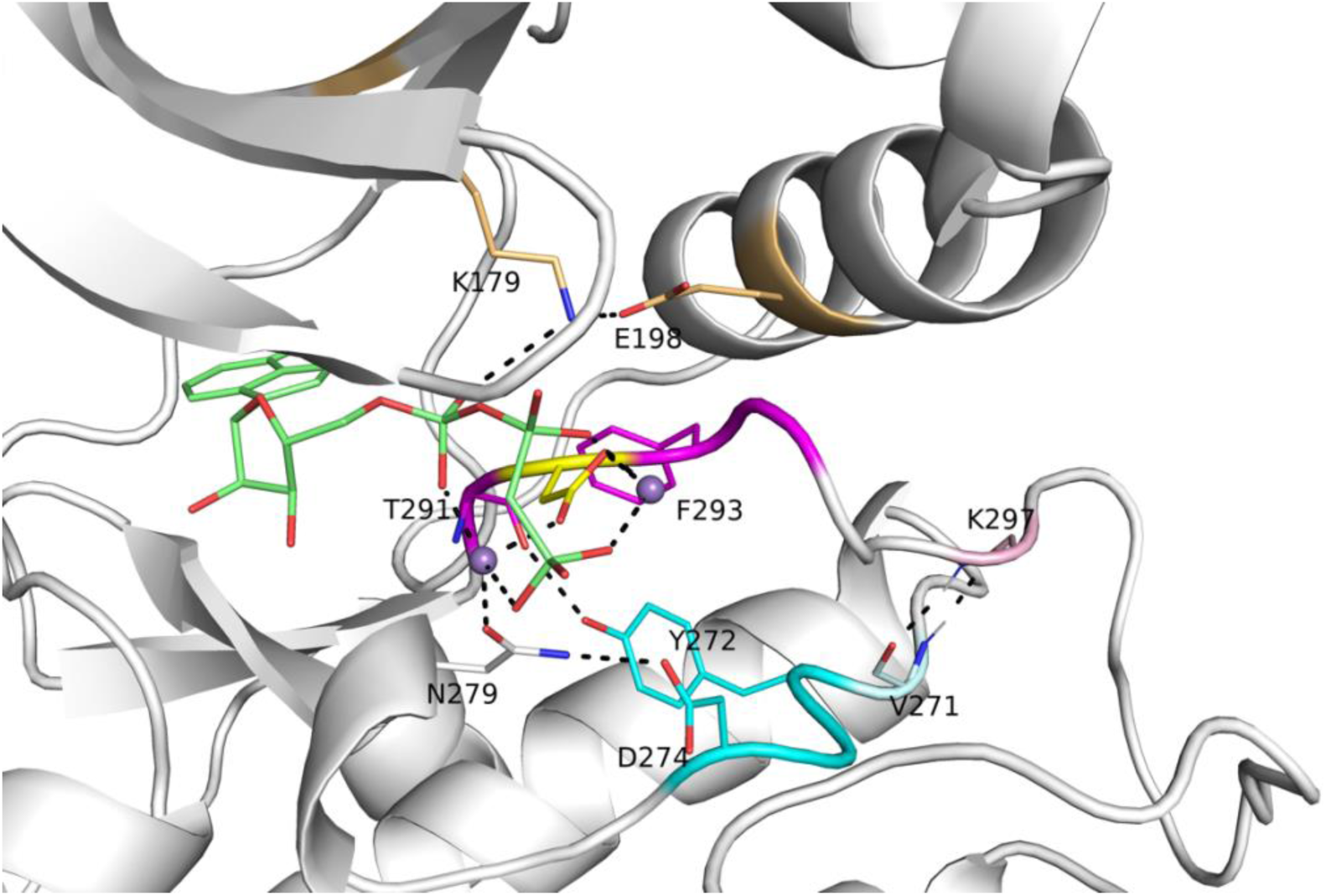
Active site of human AKT1 (PDB:4ekk, chain A). Residues making hydrogen bonding interactions with ATP (green sticks), Mg^2+^ (purple spheres), the catalytic aspartic acid residue of the HRD motif (in AKT1, this is YRD; residues 272-274, cyan), and the aspartic residue (yellow) of the XDFG motif (residues 291-294, magenta) are shown in dashed lines. These include the salt-bridge residues of the N-terminal domain (K179, E198, gold). Residue K297, which is the sixth residue of the activation loop (light pink), makes backbone-backbone hydrogen bonds with V271, which immediately precedes the YRD motif. In the stick representations, oxygen atoms are in red and nitrogen atoms are in blue. A substrate peptide is present in this structure but not shown in this figure.

**Table 1.**
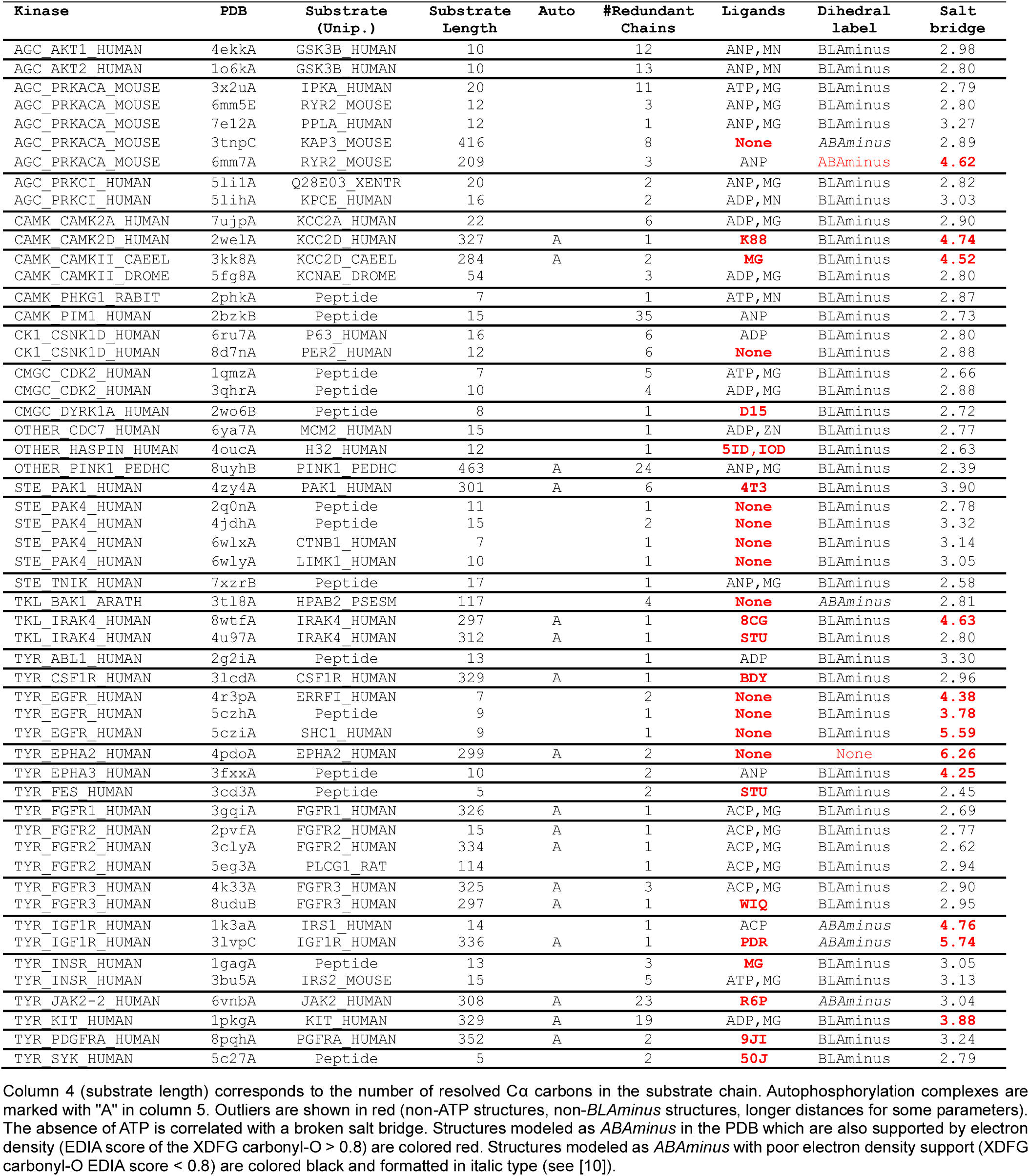
Kinase-substrate complexes in the Protein Data Bank (PDB)

We previously identified three criteria for active structures in the PDB for catalytic protein kinase domains[10]: the spatial label must be *DFGin*; the dihedral label must be *BLAminus*; this indicates that the X, D, and F residues of the XDFG motif are in the “B”, “L”, and “A” regions of the Ramachandran map respectively, and the χ_1_ rotamer of the Phe side chain is *g^-^* (~ −60°); there must be a salt bridge between the C-helix glutamic acid side chain and the beta strand 3 lysine side chain (the WNK kinases are an exception to this rule). In **Table 1**, 47 of the 54 kinase-substrate pairs are in the DFGin-BLAminus conformation. As we noted previously, *ABAminus* structures involve a peptide flip [29] from *BLAminus* structures, and as much as 50% of these may be missolved [10]. Of the six *ABAminus* structures, only one of them has good electron density for the X residue’s carbonyl oxygen (diagnostic of a mismodeled structure when the electron density is low). Twelve of 54 structures have a broken salt bridge but only two of these also have an ATP analog bound. All the structures are “Chelix-in”.

We examined the 54 substrate-bound structures listed in **Table 1** for additional characteristics of the activation loop structure that may be required for binding substrates by determining contacts of the substrate with residues in the activation loop. These residues are in the appropriate position for forming an unobstructed substrate binding groove. Besides the conformation of the DFG motif and presence of the N-terminal domain salt bridge listed in **Table 1**, two other features are evident in the substrate-bound structures. The first is that the first few residues of the activation loop, up to at least the sixth residue (colored teal in **Figure 2**), have similar conformations and positions across all the structures, regardless of family. The second is that the C-terminal segment of the activation loop, up to at least 10 residues from the end of the activation loop, also shares a common conformation and position across family members. This segment is sometimes referred to as the “P+1 loop,” since it binds the side chain of the substrate residue immediately after the phosphorylation site [17, 30]. In **Figure 2**, residues 8 and 9 from the end of the activation loop (APE8 and APE9) are shown in dark orange. In non-TYR kinases, the conformation of residues 8-11 from the end of the activation loop resembles the hull shape of an upside-down, round-bottom boat. In TYR kinases, APE8 and APE9 are also in a common position, although the structure fluctuates in APE10 and APE11 far more than in the non-TYR kinase members.

**Figure 2.**
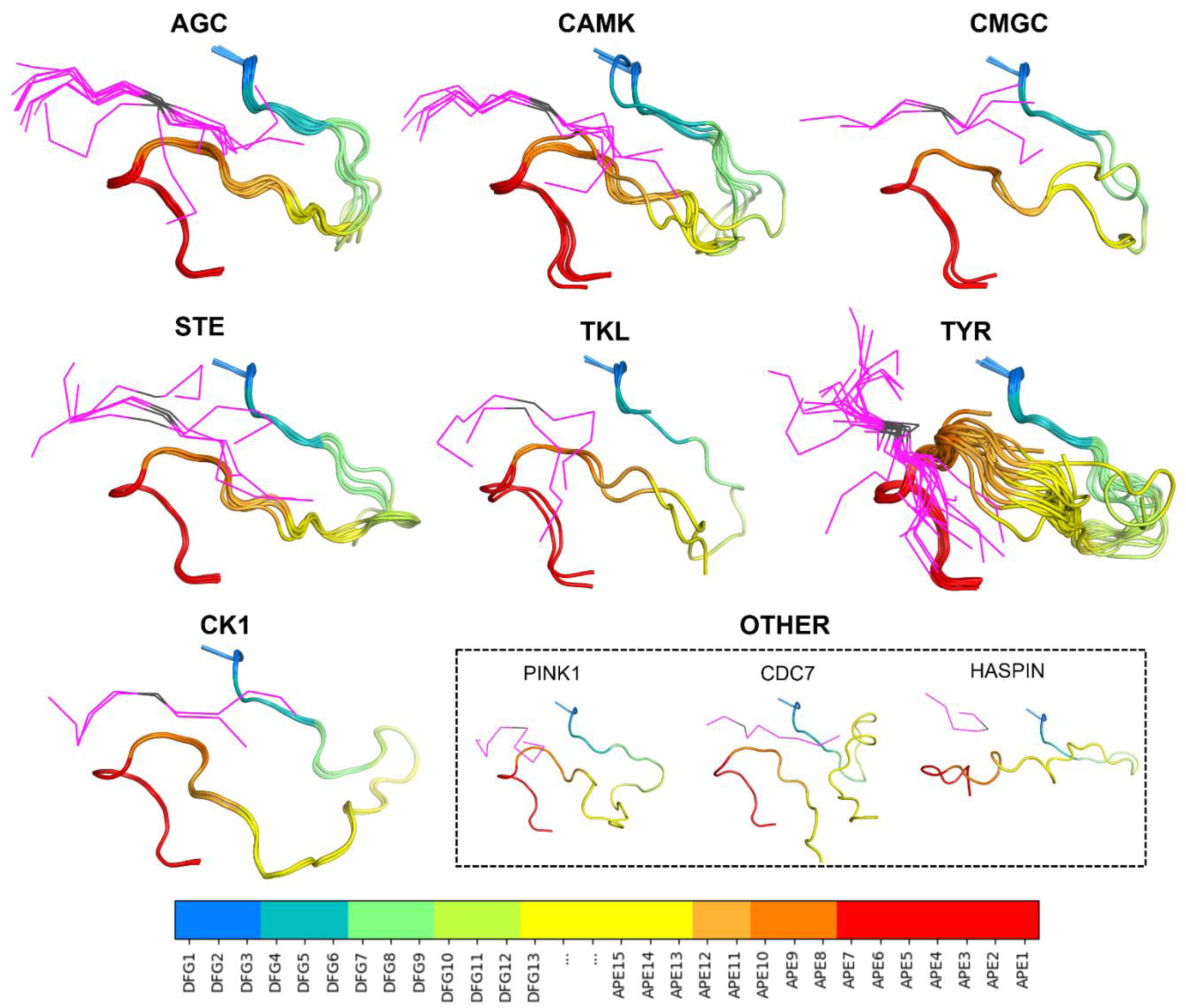
Substrate-bound structures from Table 1 organized by family. In each figure, the substrate peptides (or pieces of longer proteins) are in magenta, the phosphoacceptor is colored black and the activation loop is colored according to residue number (see bottom legend). In the legend, N-terminal activation loop residues are labeled DFG*n* where *n* is the *n*^th^ residue from the beginning of the loop, starting from DFG Asp (*n*=1) and ending at DFG13 (*n*=13). C-terminal activation loop residues are labeled APE*n* in reverse order where *n* is the *n*^th^ residue from the end of the loop, starting from APE15 (*n*=15) and ending at APE1 Glu (*n*=1).

Below, we validate our previously defined active-state criteria: 1) DFGin; 2) BLAminus; 3) presence of salt bridge. Guided by the structures of substrate-bound kinases shown in **Figure 2**, we extend the criteria to include: 4) the activation loop must be “extended” as determined by the presence of a backbone-backbone hydrogen bond between the sixth residue of the activation loop (X in DFGxxX) and the residue before the HRD motif (X in XHRD); 5) the C-terminal segment of the activation loop must be positioned for binding a substrate, as determined by the conformation of residues six through twelve from the end of the activation loop (X in XXXXXXXxxAPE); 6) the backbone dihedral angles of the HRD motif must be in the α and L regions of the Ramachandran map for the His and Arg residues respectively. We also consider the presence of the regulatory spine defined by Kornev and Taylor [30]. We review each of these in turn.

### DFGin conformation

The position of the DFG-Phe residue determines, in part, the position of the catalytic DFG-Asp residue. As we did previously [10], we define DFGin by the distance between the DFG Phe Cς atom and the Cα atoms of two residues in the N-terminal domain: the Lys residue in the β3 strand of the N-terminal domain salt bridge and the “Glu4” residue in the C-helix (**Figure 1**), which is the residue four residues following the Glu residue of the salt bridge. Based on these distances, structures are labeled as follows: *DFGin*, where the DFG-Phe residue is near the C-helix Glu4 residue but far from the Lys residue; *DFGout*, where the Phe residue is far from the C-helix Glu4 residue and close to the Lys residue; and *DFGinter*, where the Phe residue is not far from either the Glu4 or Lys residues. These distances are plotted for substrate-bound and a *control group* consisting of substrate-free structures of the same kinases in **Figure 3A**. All 248 substrate-bound structures identified in the PDB are DFGin (required for the *BLAminus* and *ABAminus* conformations of the XDF motif).

**Figure 3.**
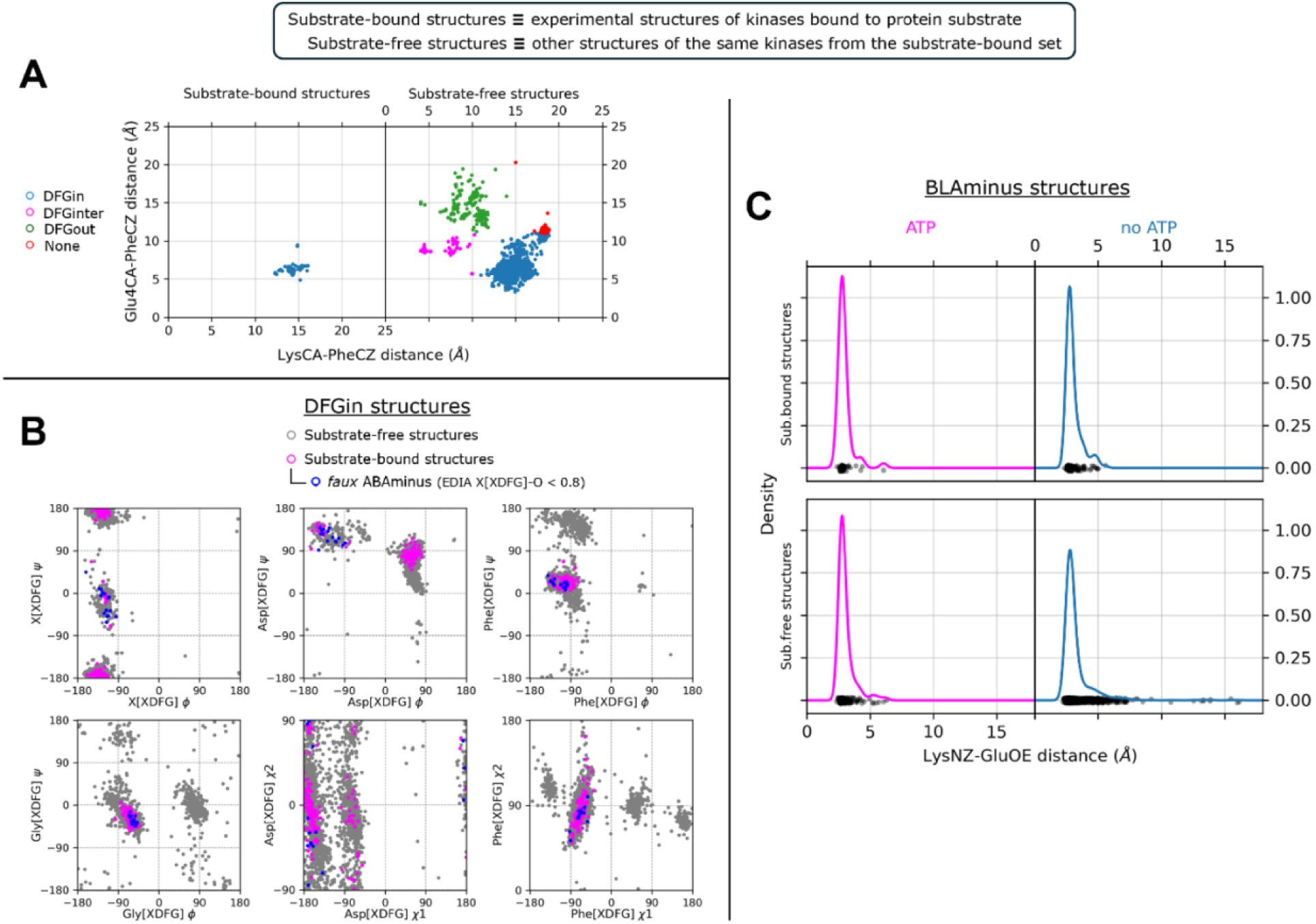
Standard measurements of kinase active state and the effects of ATP. **A.** Defining DFGin and DFGout structures. Distances of the Cς atom of DFG-Phe from the Cα atoms of the salt bridge Lys residue and Glu4 (4 residues after the salt bridge Glu) for two different datasets: 248 substrate-bound structures (*left* panel), and substrate-free structures of the same kinases (*right* panel), as a control. **B.** Ramachandran plots and side-chain dihedral angle plots for the XDFG motif residues of substrate-bound structures in the DFGin state (*magenta*) and substrate-free DFGin structures of the same kinases (*gray*). Substrate-bound structures modeled as *ABAminus* with poor electron density support (EDIA score of X[XDFG] carbonyl oxygen < 0.8) are colored blue. **C.** Distribution of N-terminal domain Lys-Nς (β3-strand) and Glu-Oε (αC-helix) distances for substrate-bound structures (*top* panels) and substrate-free structures of the same kinases (*bottom* panels) in the *BLAminus* state, comparing distances in structures bound to ATP analogs (*left* panels, magenta) vs those without ATP analog (*right* panels, blue). The presence of ATP analogs biases the formation of the Lys‒Glu salt bridge. The presence of protein substrate biases the formation of this salt bridge even in the absence of ATP analogs. In the absence of substrate and an ATP analog, *BLAminus* alone cannot guarantee salt bridge formation.

### BLAminus conformation and Salt bridge formation

The conformation of the XDFG motif and the formation of the salt bridge in the N-terminal domain work together to form an active site capable of binding ATP and magnesium ions for the phosphorylation reaction. These interactions are shown in **Figure 1**, where the Asp of the DFG motif interacts with the active site magnesium ions which chelate ATP. The carbonyl oxygen of the residue before the DFG motif (X of XDFG, T291) forms a hydrogen bond with the Tyr residue of the YRD motif in AKT1 (usually HRD, but YRD in AKT1). This hydrogen bond helps position the catalytic aspartic acid residue of the Y/HRD motif, which interacts with the Ser, Thr or Tyr hydroxyl atoms of substrate residues to be phosphorylated. The *BLAminus* conformation is required for these interactions [10]. As noted above, *ABAminus* structures involve a “peptide flip” of the X-D residues [29], such that the carbonyl of the X residue points upwards and does not interact with Y/H of the Y/HRD motif (due to a 180° change in ψ at the X residue of XDFG and a 180° change in ϕ of the Asp). Many of these structures are missolved and should be *BLAminus*, as demonstrated by poor electron density for the X residue carbonyl oxygen [10, 31]. Of the 248 substrate-bound structures, 34 of them are *ABAminus*: 18 of these 34 have poor electron density of the X residue’s carbonyl oxygen, according to the EDIA (Electron Density score for Individual Atoms) program [32]. The backbone and side-chain dihedral angles of the XDFG motif for all substrate-bound structures identified in the PDB (including those listed in **Table 1**) are plotted in **Figure 3B**.

The Lys of the N-terminal domain salt bridge interacts directly with the alpha-beta phosphate linkage of ATP. The Glu of the salt bridge helps position the Lys in this interaction [16]. The *g*-minus rotamer of the Phe side chain is required for this interaction, since the *g*+ rotamer of the inactive *DFGin-BLAplus* and *DFGin-BLBplus* conformations points upwards (instead of downwards as in the *BLAminus* and *BLBminus* conformations) and pushes the C-helix outwards, breaking the salt bridge [10]. In **Figure 3C** the distribution of distances of the salt bridge atom pairs (Nς in the β3 Lys residue with Oε_1_ or Oε_2_ in the C-helix Glu residue, whichever is shorter) is compared for *BLAminus* structures with/without ATP and with/without protein substrate. When *BLAminus* structures are bound with ATP, the salt bridge is strongly favored with a mean distance of about 3.0 Å regardless of whether protein substrate is also bound (left panels of **Figure 3C**). However, in the absence of ATP and protein substrate there is a significant number of *BLAminus* structures with broken salt bridges. From this we emphasize that the *BLAminus* state alone is an insufficient proxy for catalytic potential, and the salt bridge must be measured as well to determine whether a structure would form catalytically competent interactions with ATP.

### ActLoopNT *and* ActLoopCT conformations required for substrate binding

To investigate potential requirements for substrate binding, we determined which residues in the activation loop form direct contacts with substrate residues (any atom-atom contact within 5 Å between substrate residues and the DFG…APE sequence). The results for unique complexes from **Table 1** are shown in **Figure 4** and listed in **Supplementary Table 4**. In **Figure 4**, we plotted the frequency of contact of substrate residues P-5 to P+5 (where P0 is the phosphorylation site itself) with activation loop residues counting from the beginning of the activation loop (DFG1=Asp, DFG2=Phe, DFG3=Gly, etc.) and from the end of the activation loop (APE1=Glu, APE2=Pro, APE3=Ala, etc.). The results are shown in **Figure 4A** and **Figure 4B** for non-TYR and TYR family kinases respectively.

**Figure 4.**
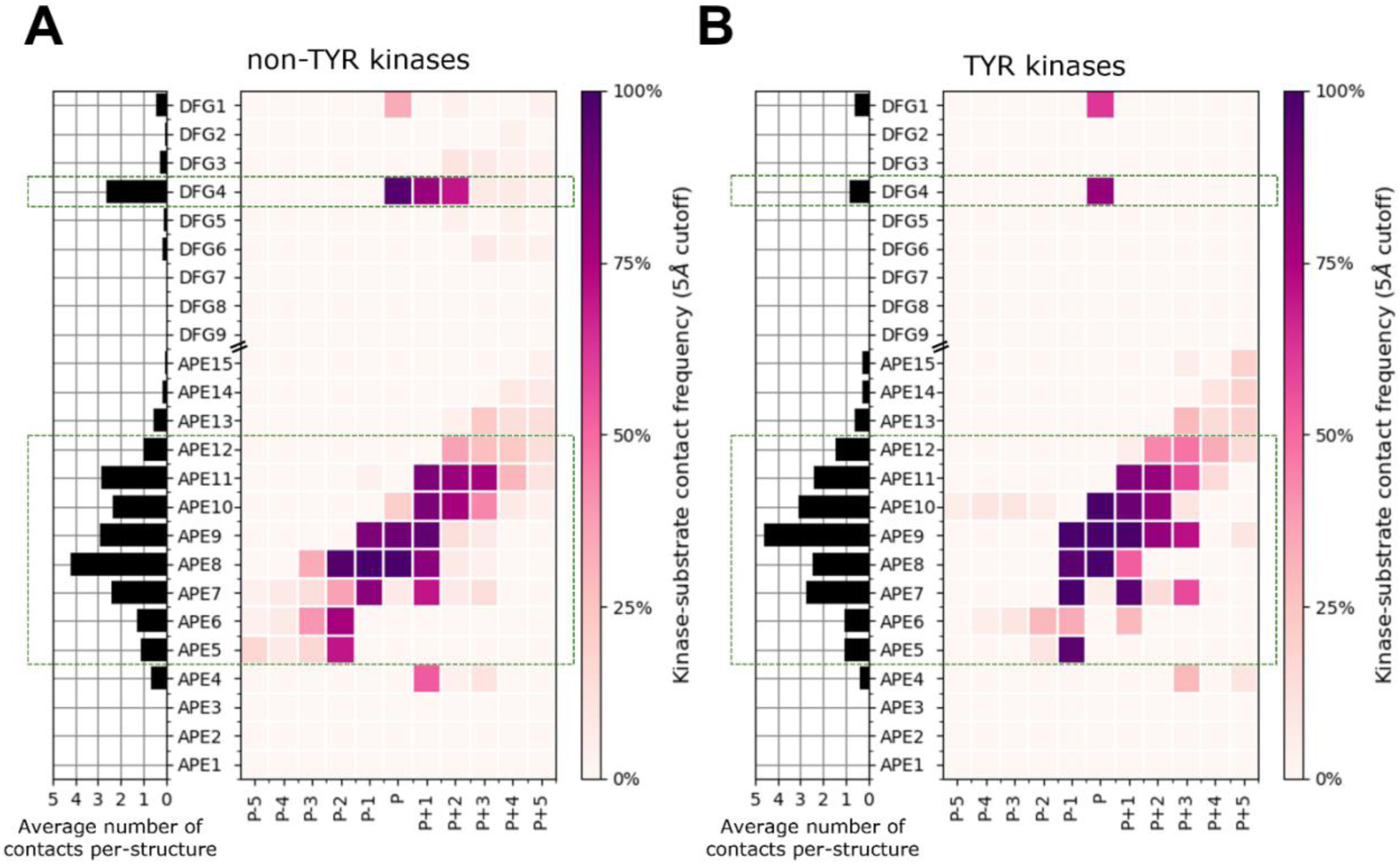
Substrate orientation and activation loop contacts for all 248 substrate-bound kinases in the PDB. N-terminal activation loop residues are labeled DFG*n* where *n* is the n^th^ residue from the beginning of the loop, starting from DFG Asp (*n*=1) and ending at DFG9 (*n*=9). C-terminal activation loop residues are labeled APE*n* in reverse order where *n* is the *n*^th^ residue from the end of the loop, starting from APE15 (*n*=15) and ending at APE1 Glu (*n*=1). Substrate residues are labeled P-5 through P+5 where P is the substrate phosphoacceptor. Activation loop residues that form contacts with approximately one or more substrate residues per structure (on average) are highlighted with dashed boxes. Left-side panel in **A** and **B**: Average number of substrate contacts formed by each activation loop residue per-structure, which is ≥1 for APE12-APE5. **A.** Substrate contacts with the activation loop in non-TYR-family kinases. Left panel: Average number of substrate contacts formed by each activation loop residue per-structure, which is ≥1 for DFG4 and APE12-APE5. **B.** Same as (A) but for TYR-family kinases, which bind substrates in an overall different orientation than non-TYR-family kinases.

TYR-family kinases bind substrates in an overall different orientation from non-TYR kinases. Relative to non-TYR kinases, TYR kinases lack contacts between P+1/P+2 and DFG4 and between P-2 and APE8 and gain contacts between P-1 and APE5, between P and APE10, and between P+2 and P+3 with APE9. For both families, contacts with the substrate P-sites and flanking residues occur mainly with the N- and C-terminal segments of the activation loop. We focus our analysis on these two segments to arrive at a set of conformational criteria for active kinases beyond what is currently described in the literature. Structural depictions of these criteria and examples of “pseudo-active” structures that violate them (structures which we now designate as “inactive”) are depicted in **Figure 5** and referenced in the following sections.

**Figure 5.**
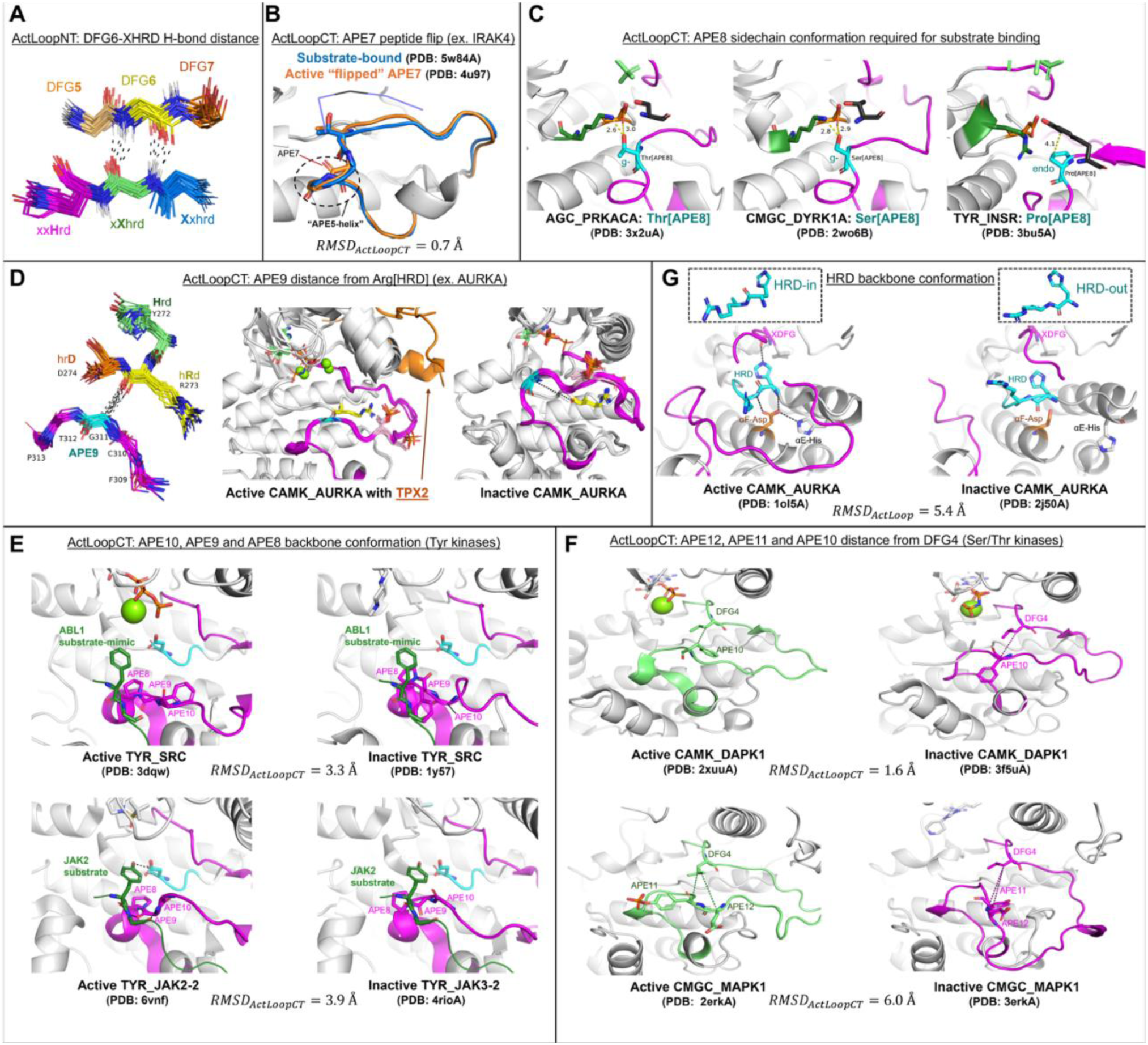
Conformational states involving the activation loop that correlate with the binding of protein substrates. **A.** Beta bridge hydrogen bonds between DFG6 and XHRD residues in kinase-substrate complex structures listed in Table 1. The carbon atoms are colored as follows: DFG5 (gold), DFG6 (yellow), DFG7 (orange), Xxhrd (blue), xXhrd (green), xxHrd (magenta, including the side chain, which is sometimes Tyr). Oxygen atoms are in red, hydrogen atoms are in white (modeled with PyMol), and nitrogen atoms are in blue. Hydrogen bonds in a few selected structures are marked with dashes. **B.** Example of an APE7-APE6 peptide flip in IRAK4 where both the flipped and un-flipped structures are considered active by our criteria. The differences in global ActLoopCT conformation between the two structures is very small (RMSD ≈ 0.7 Å). The active substrate-bound structure of IRAK4 and its substrate are colored blue, and the active peptide-flipped structure is colored orange. **C.** Non-TYR-family kinases typically have Thr at APE8 (*left*) or Ser (*center*) and the *g^−^* rotamer of these sidechains allows APE8 to form simultaneous hydrogen bonds with the conserved Asp and Lys of the HRD loop (HR**D**x**K**). Tyr kinases have Pro at APE8 (*right*) where the gamma carbon forms a short contact with Tyr substrates when it is in the Cγ-*endo* rotamer state (χ_1_>0°). Transition to Cγ-*exo* (χ_1_<0°) would likely inhibit catalysis as described in the main text. **D.** (*left* panel) Contact between Cα of the APE9 residue (XxxxxxAPE) and backbone carbonyl oxygen of HRD2 (conserved Arg of H**R**D motif). His/Tyr (green), Arg (yellow), and Asp (orange) of the HRD loop are shown in sticks, including side chains, numbered according to AKT1 residues 272-274. Residues APE11 (AKT1 F309, magenta), APE10 (C310, magenta), APE9 (G311, cyan), APE8 (T312, magenta), APE7 (P313, magenta) are shown in sticks without side chains. These structures correspond to PDB entries from the AGC, CAMK, CMGC, STE, and TKL kinases listed in Table 1 of the main text. (*middle* and *right* panels) Two conformations of human AURKA. (*middle* panel) Active structures with bound TPX2 (orange): PDB: 1ol5, 3e5a, 3ha6, 5lxm, 6vph. (*right* panel) Inactive structures without TPX2. PDB: 4dee, 5dt3, 5oro, 5os0, 6i2u, 6r49, 6r4d and others. Gly291 Cα (cyan sticks) is in contact with HRD-Arg255 backbone carbonyl O (yellow sticks) in the active structures (average distance 3.6 Å), while there is no contact in the inactive structures (average distance > 10 Å). The activation loop C-terminal region in the active structures resembles the structures of substrate-bound complexes, while the inactive structures would block substrate binding. **E.** Comparison of active (*left* panels) TYR-family kinases and examples from the same family which are inactive (*right* panels) due to substrate-incompatible ActLoopCT dihedrals, superimposed with a bound-substrate from the most closely related family member. In the upper panels the substrate-mimic (P0 Tyr → Phe mutant) from TYR_ABL1 (PDB: 2g2i) is superimposed with active and inactive (ActLoopCT-out) structures of TYR_SRC. In the bottom panels the substrate from active TYR_JAK2 (green) is shown for reference. **F.** Comparisons of active (*left*: green) and inactive ActLoopCT-out structures (*right*: magenta) of CAMK_DAPK1 (*top row*) and CMGC_MAPK1 (*bottom rows*) (also called ERK2). The inactive DAPK1 structure (3f5uA) has an expanded APE10-DFG4 distance relative to the active conformation (2xuuA). The inactive MAPK1 structures have a contracted APE11-DFG4 distance (4xj0A) and expanded APE12-DFG4 distance (3erkA) relative to the active, phosphorylated conformation (2erkA). **G.** Structural visualization of CAMK_AURKA in the active (*left*) vs inactive/HRD-out (*right*) state. The HRD-out conformation is a peptide flip relative to HRD-in that breaks a hydrogen bond between the HRD-Arg backbone nitrogen and the conserved Asp in the αF-helix (orange). These hydrogen bonds stabilize a salt bridge between the αF-Asp and conserved His in the αE-helix. The inactive DFGinter structure of CAMK_AURKA, 2j50A (right panel) demonstrates how the HRD peptide flip can propagate structural changes to the XDFG motif and αE-His, resulting in a more globally inactive conformation.

#### ActLoopNT

We find that most substrates have a contact with one or more of the DFG-adjacent residues and, notably, residue 4 of the activation loop (DFG4). For all substrate-bound structures, the DFG4 residue interacts with the phosphoacceptor residue (“P-site”). For non-TYR kinases, due to the different orientation of the substrate in these kinases, the DFG4 residue is also in contact with the P+1 residue, and in a smaller number of cases, forms contacts with P+2. Based on these observations, it is not surprising to find that, in the activation loops of active kinase structures, DFG4 (DFG**X**xx) is flanked by two structural motifs that constrain its positioning: besides DFG-in/*BLAminus* (**XDF**Gxxx), there are two anti-parallel beta strands formed by backbone-backbone hydrogen bonds between residue 6 of the activation loop (DFGxx***X***) and the residue immediately preceding the HRD motif (***X***hrd) (**Figure 5A**). These short beta strands (3 residues each, centered on DFG6 and Xhrd) have been labeled beta strand 6 (comprising the XXH residues of XXHrd) and beta strand 9 (comprising residues 5-7 of the activation loop, dfgxXXX) in protein kinase structures [30]. This hydrogen bond is present in all of the substrate-bound structures in **Table 1** (“DFG6” in the table) and is likely an evolved feature of the active conformation that positions DFG4 to interact with substrate P-site and P+1 residues.

This DFG6/Xhrd hydrogen bonds also cooperate with the conformation of the XDF motif: 99% of *BLAminus* structures (660 of 666) with bound ATP or ATP analogs contain this hydrogen bond with the minimum N-O or O-N backbone-backbone distance less than 3.6 Å (**Figure 6**, upper and middle panels). The only exceptions are the partially disordered activation loop structures of the bacterial kinase PknB (4eqm chains D and E) and TYR_HCK (1ad5 chains A and B), and an activation-loop swapped structure of STE_MAP4K1 (6cqd chains A and B) in which the N-terminal portion of the activation loop forms an α-helix. Without ATP and protein substrate, 2% of *BLAminus* structures have a broken DFG6/XHRD hydrogen bond (**Figure 6**, middle right panel). Although this fraction is small (comprising ~150 PDB chains), it indicates that the DFG6 criterion is useful for distinguishing active conformations among *BLAminus* structures. When the XDF state is not *BLAminus* the fraction of structures with a broken DFG6/XHRD hydrogen bond increases from 2% to 75% (**Figure 6**, bottom right panel).

**Figure 6.**
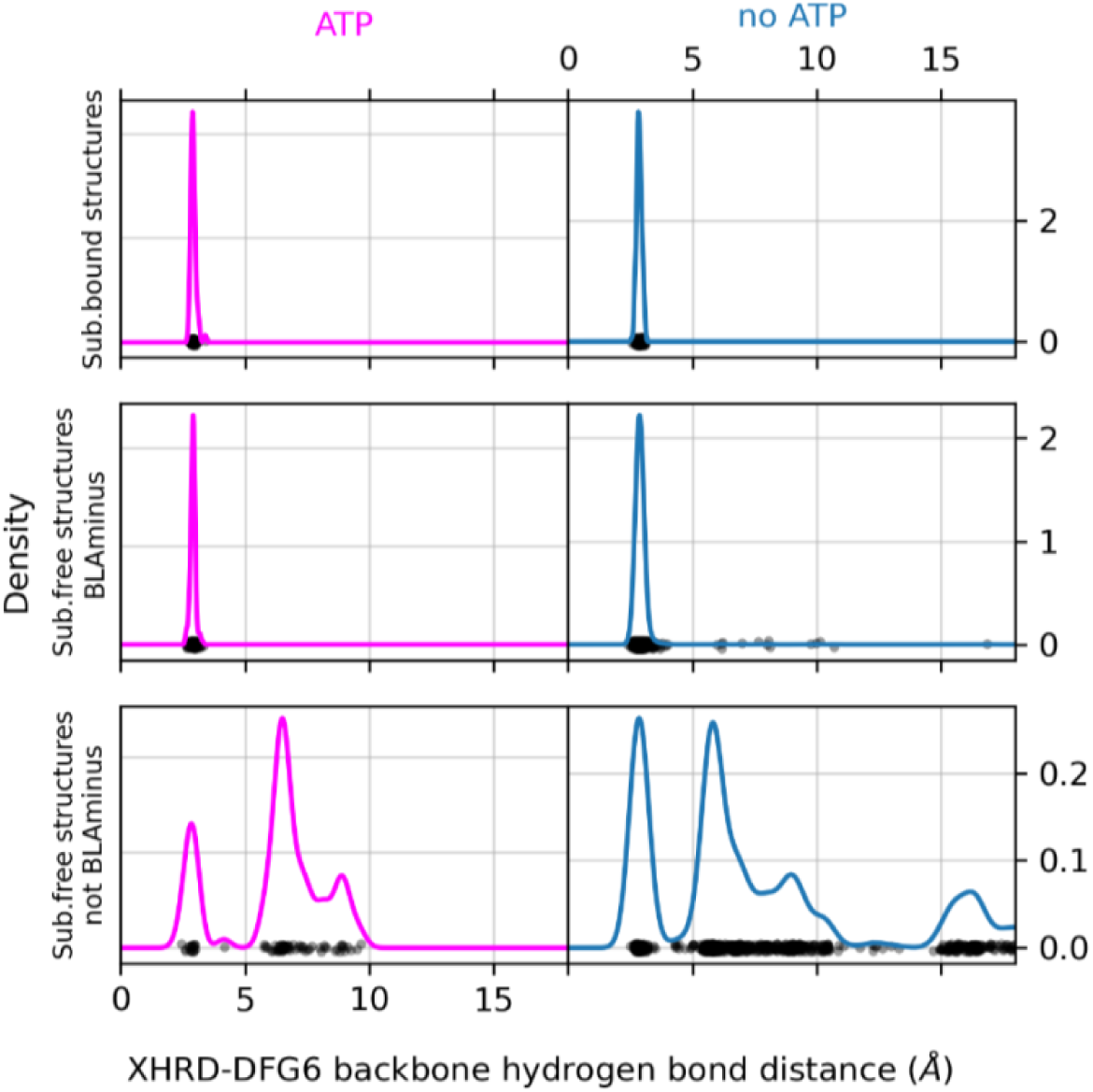
Interactions of residue 6 of the activation loop and the residue before the HRD motif (“XHRD”) Distribution of the XHRD-DFG6 backbone-backbone hydrogen bond distance in substrate-bound structures (*top* panels), substrate-free structures which are also *BLAminus* (*middle* panels), and substrate-free structures which are *not BLAminus* (*bottom* panels), comparing structures bound to ATP analog (*left* panels, magenta) vs those which are not bound to ATP analog (*right* panels, blue). The distance plotted is the minimum of the N-O or O-N distances between these two residues. The DFG6 residue is identified with the last X in the DFGxxX sequence, where x/X is any amino acid. The XHRD-DFG6 hydrogen bond is formed (ActLoopNT-in) in all substrate-bound structures regardless of whether an ATP analog is present (*top* panels). In the control structures without protein substrate and without ATP analog (*middle* and *bottom* panels) the distance distributions adopt a wider range. Structures that are not *BLAminus* are mostly ActLoopNT-out.

#### ActLoopCT

The conformation of the C-terminal end of the activation loop is critical for binding substrates. Most substrate-bound structures in **Table 1** contain contacts between the substrate and residues 4-12 from the end of the activation loop (detailed in **Supplementary Table 4**), which usually ends in the sequence motif “APE” (APE in 306 kinase domains, SPE in 47, PPE in 29, and other sequences in the remaining 55 kinases). The positioning of the APE motif is conserved in most catalytic kinases, owing to its close packing with the αF-helix.

Differences in the bound orientation of substrates for the TYR family compared to non-TYR families require the C-terminal region of the activation loop to adopt a different, more distended conformation (**Figure 2**). This results in distinct sets of contacts with the substrate (**Figure 4**), suggesting that TYR family kinases require different conformational criteria of the activation loop C-terminal region to accurately define the active conformation. In TYR kinases, the substrate binds directly to these residues in the form of a short beta strand (magenta lines in **Figure 2**). In other kinase families, the substrate binds to a groove between the N and C-terminal segments of the activation loop.

##### APE5 helix

Moving in the N-terminal direction from the APE motif, APE5-APE7 belong to a short helix-like structure that interacts with substrate residues P-1 through P-5 (**Figure 4**). The APE5 helix can be captured by imposing conformational criteria on the backbone dihedrals ϕ and ψ of APE7 and APE6, requiring both residues to fall in the broad alpha helical “A” region of the Ramachandran map: ϕ ∈ (−180°,0°), ψ ∈ (−100°,50°). In addition to interacting directly with substrate residues P-3 through P-1 (**Figure 4**), the APE5 helix also constrains the conformational variability of the rest of the C-terminal segment (APE8 – APE12) allowing these residues to form an extended binding surface for the substrate as described below.

In a few structures, there is a peptide flip, such that residues APE7 and APE6 fall in the B/beta and L/left-handed regions respectively, preserving the trajectory of the backbone structure (**Figure 7A-B**). For example, residues APE7 and APE6 of TKL_IRAK4 (5w84A) appear in the B-L Ramachandran regions (**Figure 7A**, left panel), and structural superposition with the substrate-bound structure of IRAK4 (4u97A) reveals that the main difference between the two structures is the orientation of the APE7 carbonyl oxygen, which is flipped “up” in the former (B-L) and pointing “down” in the latter (A-A) (**Figure 5B**). The rest of the ActLoopCT conformation is largely unaffected by the flip (ActLoopCT RMSD < 1Å). The structural change involves a flip from a Type I beta turn (A-A) to a Type II beta turn (B-L) [33]. In the active-kinase criteria, we allow for this peptide flip (see below), such that for APE7: ϕ ∈ (−180°,0°), ψ ∈ (50°,180°) and for APE6: ϕ ∈ (0°,180°), ψ ∈ (−50°,100°). Most of the kinases that contain the flip have a glycine residue at position 6 (AGC_GRK1-7, TKL_LRRK1-2, OTHER_ERN1-2), which is unsurprising given the L conformation.

**Figure 7.**
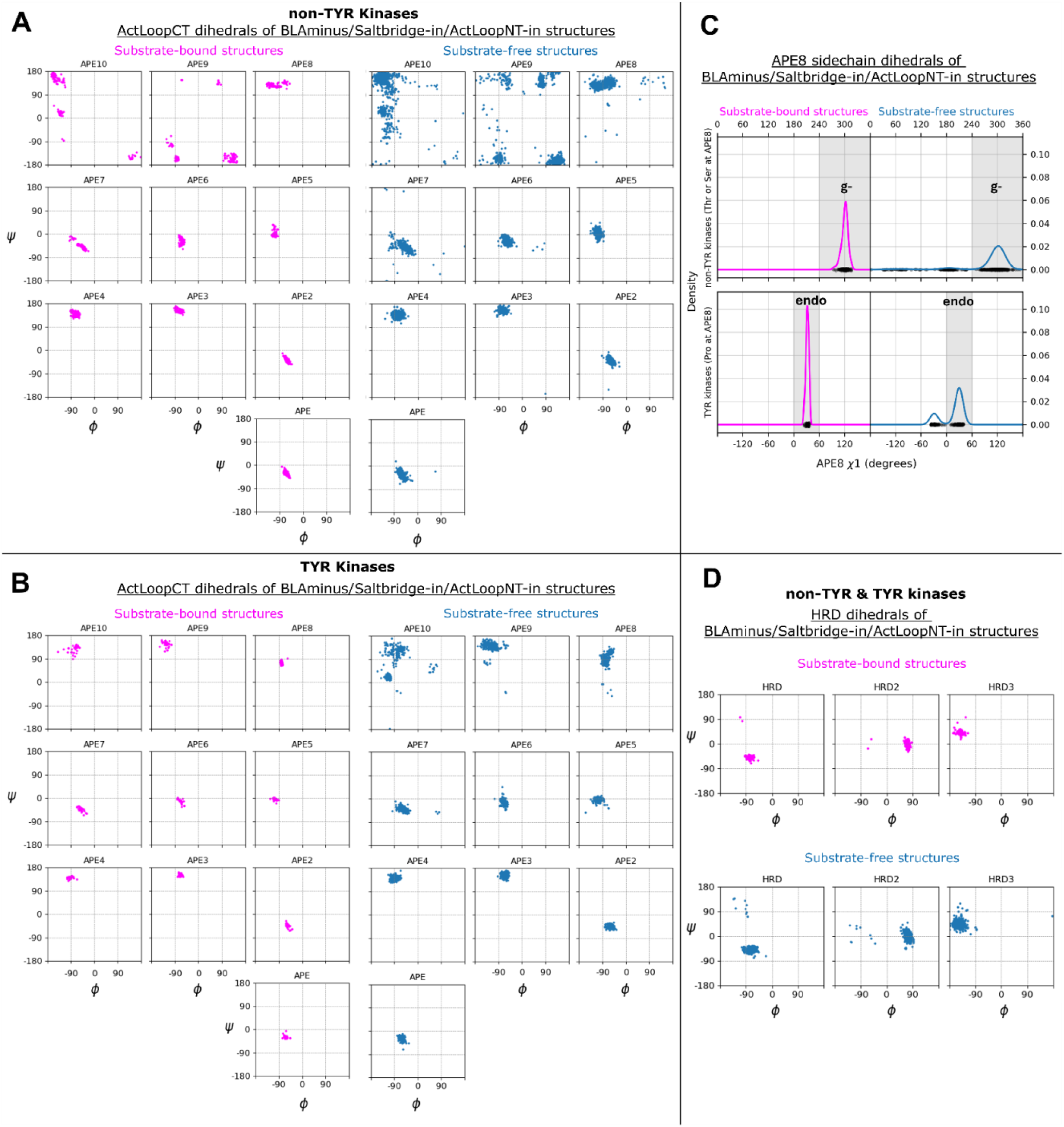
Dihedral criteria for Activation Loop C-terminus and catalytic HRD loop residues. Comparison of 248 substrate-bound structures vs substrate-free structures of the same kinases (as controls) which satisfy previously defined active-state criteria (*BLAminus*, Saltbr-in, ActLoopNT-in) to identify the need for active-state criteria based on dihedral angles. **A.** ActLoopCT backbone dihedrals for substrate-bound non-TYR kinase structures (magenta, *left*) compared with the substrate-free set comprised of the same kinases (blue, *right*). B. ActLoopCT backbone dihedrals for substrate-bound TYR kinase structures (magenta, *left*) compared with the substrate-free control set comprised of the same kinases (blue, *right*). **C.** APE8 sidechain χ_1_ dihedrals for non-TYR kinases (Ser[APE8] and Thr[APE8]) and TYR kinases (Pro[APE8]) compared between substrate-bound (*left,* magenta) and substrate-free (*right*, blue) sets. Ser[APE8] and Thr[APE8] side chains in the non-TYR kinase substrate-bound set are always in the g-conformation. Pro[APE8] sidechains in TYR kinase substrate-bound set (*bottom left* panel) are always Cγ *endo* (χ_1_>0°). APE8 side chains in the substrate-free sets can deviate significantly from these conformations. **D.** HRD loop backbone dihedrals for the substrate-bound (*top*) and substrate-free (*bottom*) sets plotted for both non-TYR and TYR kinases. Peptide flips at the HRD-His and HRD-Arg positions (conserved His and Arg, respectfully) can be observed in the substrate-free set which has functional consequences for kinase activity.

##### APE8

For both TYR and non-TYR kinases, the APE8 residue is critical for binding substrates, interacting with substrate residues P-2 to P+1 with high frequency in non-TYR kinases (**Figure 4A**) and with P-1 to P+1 in TYR Kinases (**Figure 4B**) as well as catalytic residues in the HRD loop. APE8 is a conserved Pro in all 83 TYR kinases, and Ser or Thr in 335 non-TYR kinases (95%). Atypical residue types such as Asn can also be observed at this position (e.g., OTHER_WEE1 and OTHER_WEE2). We also note that the APE motif and its helical structure are missing altogether from three unusual human kinases (OTHER_HASPIN (4oucA from **Table 1**), OTHER_PKDCC, OTHER_TP53RK) which do not have an APE motif [3]. None of the APE criteria (above and below) are applied to these three kinases.

For non-TYR kinases, all substrate-bound structures have a backbone conformation in the beta region of the Ramachandran map (**Figure 7A**), while non-substrate bound structures in the *BLAminus* conformation are sometimes in the upper right of the Ramachandran map (top right of **Figure 7A**). Flipping ϕ of APE8 (assuming APE7 and APE6 are properly placed) would break typical backbone-backbone hydrogen bonds of APE9 with P+1 of the substrate. We therefore place a requirement on the ϕ,ψ values of APE8 of non-TYR kinases: ϕ ∈ (−180°,0°), ψ ∈ (50°,180°).

When APE8 is Ser or Thr in non-TYR kinases, it is always found in a gauche-minus rotamer (−60°). The g^−^ rotamer orients the gamma hydroxyl to accept a hydrogen bond from the side-chain Nς of a conserved Lys in the HRD loop (HRDx**K**xxx); simultaneously it donates a hydrogen bond to the side-chain carboxylate of the conserved Asp in the HR**D** motif (**Figure 7C**, top panel and middle panel) (**Figure 5C**). These hydrogen bonds are present in all but three of the substrate-bound kinases from **Table 1** that contain Thr/Ser at APE8 (not counting 5lihA which is missing atoms from Lys[HRDx**K**]). This configuration promotes catalysis by orienting the other carboxylate oxygen atom “upward” toward the substrate hydroxyl. Consequently, we place constraints on the χ_1_ rotamer of the APE8 residue: when this residue is a Ser or Thr in non-TYR kinases, we require the g-rotamer: χ_1_ ∈ (−120°, 0°).

In TYR family kinases, APE8 is instead Pro. In substrate-bound structures, the backbone conformation is always close to (−75°,+85°) (**Figure 7A**), and the side-chain conformation is always in a Cγ-*endo* rotamer (χ_1_ ~ +30°) (**Figure 7C**, bottom panel). This is consistent with data from the PDB, where nearly all Pro residues with ψ between 45° and 135° are in the <u>Cγ-*endo* rotamer</u> and their ϕ dihedral is < −70° [34]. This proline is in the middle of an inverse gamma turn with a hydrogen bond between the carbonyl oxygen of APE9 and the backbone NH of APE7 (NB: APE7 is a proline in 45% of non-TYR human kinases [3]). The Cγ-*endo* rotamer creates a small divot in the proline ring, creating space for the Tyr substrate which forms a short contact with the Pro beta carbon (**Figure 7C**) (**Figure 5C**); the Cγ-*exo* rotamer would lift this gamma carbon by ~1.4 Å, pushing the substrate away from its hydrogen bond acceptor and donor (Asp and Arg of HR**D**xxx**R**). The rotamer of proline is strongly correlated with the backbone dihedrals ϕ and ψ with a small energy barrier of less than 3 kcal/mol between the Cγ-endo and Cγ-exo conformations [35, 36]. Consequently, we do not put a requirement on the side-chain conformation of this residue for TYR kinases, but we do put the same restriction on the backbone dihedrals as for non-TYR kinases: ϕ ∈ (−180°,0°), ψ ∈ (50°,180°). The Pro residue at position APE8 enforces a beta conformation on the APE9 residue (see below), which is therefore a pre-Pro residue, whose Ramachandran map is distinct from non-pre-Pro residues [37, 38], with significantly reduced populations in the alpha region.

##### APE9

From examination of the substrate-bound structures, we identified a contact that is consistent with substrate binding, and which is absent in structures that likely block substrate binding: a contact (or near contact) between the APE9 Cα atom and the backbone carbonyl oxygen of the Arg residue in the HRD motif (HRD-Arg). **Figure 5D** shows this contact for 23 non-TYR kinase structures from **Table 1**. The Cα-O distance is ≤ 4.2 Å in all of these structures.

Aurora A kinase (CAMK_AURKA) is a good example of the utility of the APE9 distance criterion. In the ATP-bound, *BLAminus* state, there are two dominant conformations of the entire activation loop of AURKA, one where the APE9/HRD-Arg contact is formed and the other where the contact is broken. **Figure 5D** shows five structures that contain these contacts. This comprises five structures of AURKA with TPX2 (PDB: 1ol5, 3e5a, 3ha6, 5lxm, 6vpg). Two other structures bound with MYCN (PDB: 5g1x, 7ztl) are very similar (not shown). Both proteins are known to activate AURKA by binding to the N-terminal domain and the tip of the activation loop [39, 40]. Most *BLAminus* structures of AURKA, however, resemble the structures shown in the right panel of **Figure 5D** where the APE9-Arg contact is broken (e.g., PDB: 4dee, 5dt3, 5oro, 5os0, 6i2u, 6r49, 6r4d). In these structures, the C-terminal end of the activation loop (APE6-APE10) deviates significantly from the TPX2- and MYCN-bound structures and from the structures of substrate bound kinases in the related AGC and CAMK families. The Cα-O distances in these AURKA structures are all more than 10 Å.

We examined the distributions of this distance (**Figure 8A**) for both substrate-bound and substrate-free structures which satisfy all the active criteria described above (*BLAminus*, ActLoopNT-in, SaltBr-in, intact APE5 helix). For the substrate-bound non-TYR kinase structures, the APE9(Cα)-hRd(O) distances range from 3.4 to 4.2 Å. This suggests that the Cα-O interaction is a CH-O hydrogen bond, which has been observed in proteins [41]. For 76% of human non-TYR catalytic kinases [42], the APE9 residue is a glycine, which forms Cα-O hydrogen bonds more readily than other amino acids likely for steric reasons. This contact is broken in many substrate-free structures, with the APE9(Cα)-hRd(O) distance exceeding 6 Å even those which satisfy other active-state criteria described above (**Figure 8A** bottom panels).

**Figure 8.**
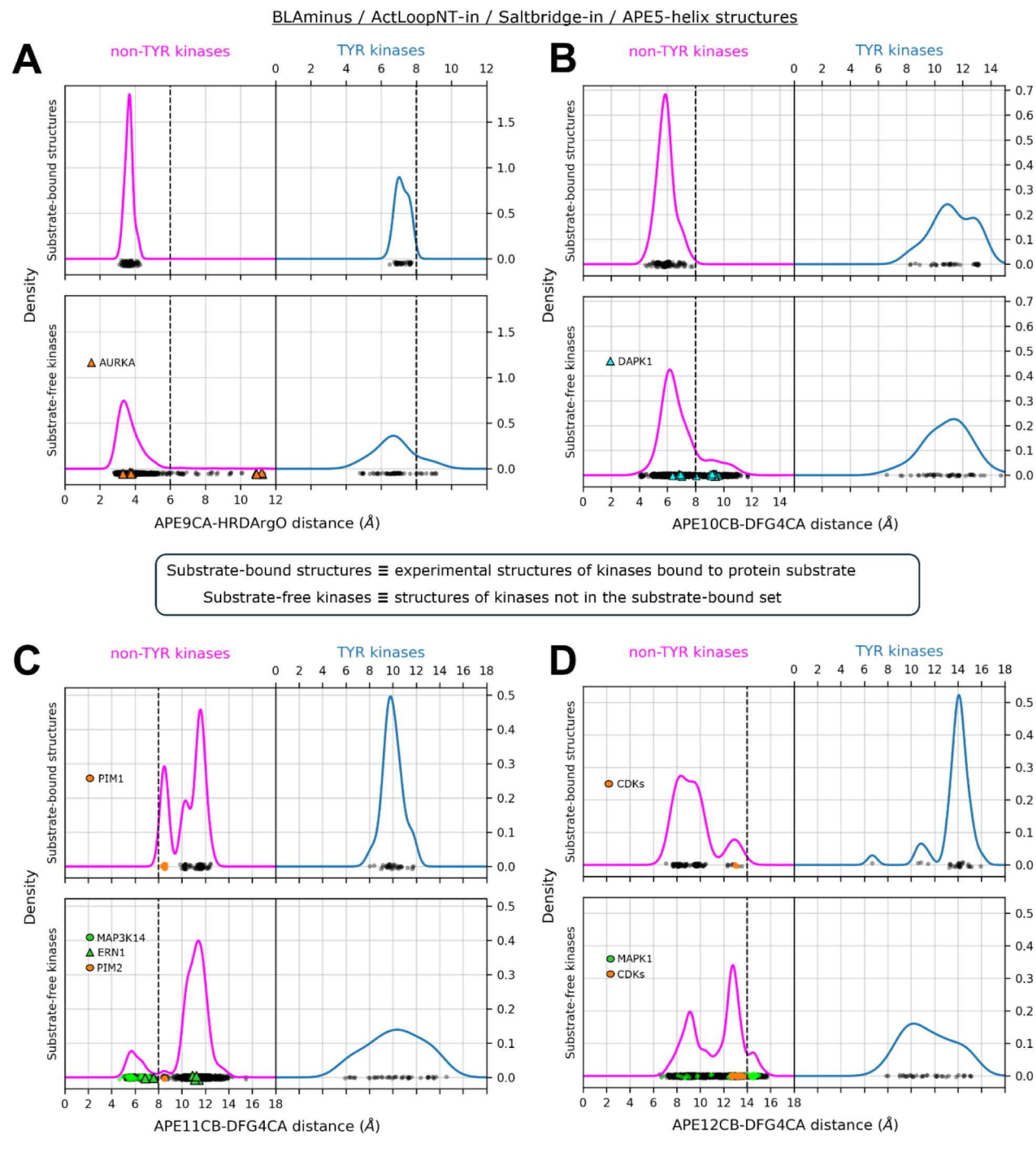
Distance criteria for Activation Loop C-terminal residues. Comparison of 248 substrate-bound structures vs substrate-free structures of the same kinases (as controls) which satisfy previously defined active-state criteria (*BLAminus*, SaltBr-in, ActLoopNT-in, APE5-helix intact) to identify the need for additional active-state criteria based on residue-residue distances that define the substrate binding surface. **A.** Distribution of the APE9-Cα/HRD-Arg-O distance in substrate-bound structures (*top* panels) compared with substrate-free kinases (*bottom* panels). Structures of CAMK_AURKA are plotted with orange triangles. Inactive structures of AURKA greatly exceed the APE9-Cα/HRD-Arg-O distance threshold which was set at 6 Å. **B.** Distribution of the APE10-Cβ/DFG4-Cα distance in substrate-bound structures (*top* panels) compared with substrate-free kinases (*bottom* panels). Structures of CAMK_DAPK1 are plotted with blue triangles. A dashed vertical line is drawn at the threshold of the APE10-Cβ/DFG4-Cα distance criteria, highlighting the presence of inactive DAPK1 structures (right of the dashed line). **C.** Distribution of the APE11-Cβ/DFG4-Cα distance in substrate-bound structures (*top* panels) compared with substrate-free kinases (*bottom* panels). Structures of PIM family kinases (CAMK_PIM1 and CAMK_PIM2) are plotted with orange circles, structures of STE_MAP3K14 are plotted with green circles, and structures of OTHER_ERN1 are plotted with green triangles. A dashed vertical line is drawn at the lower threshold of the APE11-CB/DFG4-CA distance criteria (8 Å) to highlight the separation of active CAMK_PIM1/PIM2 (right of the dashed line) from inactive OTHER_ERN1 and STE_MAP3K14 (left of the dashed line). **D.** Distribution of the APE12-Cβ/DFG4-Cα distance in substrate-bound structures (*top* panels) compared with substrate-free kinases (*bottom* panels). Structures of CDK family kinases (e.g., CMGC_CDK2) are plotted with orange circles and structures of CMGC_MAPK1 are plotted with green circles. A dashed vertical line is drawn at the upper threshold of the APE12-Cβ/DFG4-Cα distance criteria (14 Å) to highlight the separation of active CDKs (left of the dashed line) from inactive CMGC_MAPK1 (right of the dashed line).

Substrate-bound TYR kinase structures have somewhat longer APE9(Cα)-hRd(O) distances, ranging from 6 to 8 Å (**Figure 8A**, top right panel). This reflects a greater flexibility of the ActLoopCT for TYR kinases than for non-TYR kinases. Because APE9 interacts closely with the P+1 residue in TYR kinases, the upper bound on the APE9(Cα)-hRd(O) distance (8 Å) is likely important for allowing the substrate Tyr residue to reach the catalytic HRD-Asp residue. This is therefore the upper bound on this distance for TYR kinases.

##### APE10

Active TYR kinases differ significantly from non-TYR kinases in the APE9 – APE12 region to accommodate a specific binding mode for substrates [43]. TYR kinases typically have a bulky/charged residue at APE10 (most commonly Lys or Arg) with the side chain resting “on top” of the bound substrate peptide or protein, whereas for non-TYR kinases APE10 is placed “underneath” the substrate. In substrate-bound TYR kinases, the backbone of APE9 forms a short beta strand with P+1 of the substrate and APE10 and APE9 are both in the beta region of the Ramachandran map (**Figure 7B**). Hence, for active TYR kinases we require APE9 and APE10 (and APE8 described above) to have ϕ ∈ (−180°,0°), ψ ∈ (50°,180°). This is a broad region, but all active kinases identified by the full set of criteria are distributed across a narrower region (see below).

The APE8-APE10 dihedral criteria reveal the existence of inactive “ActLoopCT-out” structures of human TYR_SRC in the PDB: e.g., PDB: 1y57A and 1yi6B which are often considered “Active” in the literature but do not resemble substrate-bound structures of TYR kinases. The ActLoopCT conformation in these structures would likely interfere with substrate binding (**Figure 5E**). There is, however, a structure of chicken SRC (PDB 3dqw, chains A-C, **Figure 5E, top**) that is “ActLoopCT-in” and is quite similar to the ABL1 (PDB 2g2iB) structure listed in **Table 1**, and this structure is classified as “active” by our criteria. The chicken SRC sequence differs by only two amino acids in the kinase domain from human SRC, neither of which is in the activation loop. Hence, although there is no substrate-binding-capable structure of human SRC kinase domain in the PDB, the chicken SRC structure (PDB: 3dqW, chain A) presents a suitable active model of the human ortholog. We also note the existence of inactive ActLoopCT-out structures of TYR_JAK3-2 (JAK3 catalytic domain) in the PDB (e.g., 4rioA), which differ significantly from the substrate bound (autophosphorylation of Y836) structure of the closely related kinase TYR_JAK2-2 (6vnbA) (**Figure 5E, bottom**).

In non-TYR kinases, the side chain of APE10 is typically short and uncharged (most commonly Val or Cys). In active and substrate-bound structures of non-TYR kinases, the APE10 and DFG4 side chains pack against each other to form a component of the substrate-binding surface that interacts with the substrate P-site and P+1 residue. To capture this structural motif in our criteria for active structures in non-TYR kinases, we use a cutoff of 8 Å on the Cβ-Cα distance between APE10 and DFG4 for non-TYR kinases alone (**Figure 8B**, upper left panel). This cutoff efficiently separates two populations of DAPK1 structures (cyan triangles in Figure 8B, lower left panel); the points below 8 Å resemble active structures of other non-TYR kinases, while those above 8 Å are substantially different and would block substrate binding. Changes in the conformation of APE10 from the substrate-compatible conformation often coincide with conformational changes of other residues as described below.

##### APE11 – APE12

For the majority of non-TYR kinases, activation loop phosphorylation (typically at APE12 or APE13, and less commonly APE11 [28]) stabilizes the N-terminal and C-terminal residues in a substrate-compatible conformation. Importantly, phosphorylation at these residues constrains the conformation of downstream residues in the ActLoopCT (APE12, APE11, APE10, APE9, APE8) which interact directly with substrates (**Figure 4**). For several non-TYR kinases, the requirement for phosphorylation on the activation loop as a requirement for catalytic activity can be explained by the unphosphorylated conformational propensities of the APE11 and/or APE12 residue(s), which may interfere with substrate binding. To capture this, we measured the Cβ-Cα distance of APE11-DFG4 and APE12-DFG4 and set bounds on the active criteria for non-TYR kinases based on the distribution of distances for substrate-bound structures (**Figure 8C and 8D**, upper left panel). These bounds are (8 Å, 14 Å) and (7 Å, 14 Å), respectively.

We find that, for several inactive non-TYR kinases in the unphosphorylated state, the Cβ vectors of APE10 and APE11 have “swapped” orientations resulting in an expanded distance between APE10 and DFG4 (Cβ – Cα > 8Å) and a contracted distance between APE11 and DFG4 (Cβ–Cα < 8Å), e.g., CAMK_DAPK1 (unphos. 3dgkA, phos. 2xuuA) (**Figure 5F**), STE_MAP3K5 (unphos. 8qgyB, phos. 6e2mA), OTHER_ERN1 (unphos. 6w3bA, phos. 6w3cA) and CMGC_CDK2 (unphos. 1finA, phos. 1qmzA). Some structures violate only the lower bound of the APE11-DFG4 distance criteria, e.g., unphosphorylated CMGC_MAPK1 (4xj0A, 7w5oB) and STE_MAP3K14 (8yhwA). For comparison, CMGC_MAPK1 has phosphorylated structures in the PDB which resemble typical active structures of CMGC kinases (e.g., 2erkA). For STE_MAP3K14 however, there are no active structures of the same kinase to compare with because all existing PDB structures of MAP3K14 exhibit a contracted APE11-DFG4 distance (< 8 Å, similar to unphosphorylated ERN1 6w3bA) and have therefore been reclassified as “inactive” (**Figure 8C**, lower panel). Based on the observation that ERN1 and MAP3K14 have similar ActLoopCT sequences (^APE12^Y**IPGTE**^APE7^ in MAP3K14 vs ^APE12^G**VPGTE**^APE7^ in ERN1) we propose that the active state of MAP3K14 should resemble the structure of active ERN1 (6w3cA) (**Supplementary Figure 1**).

For CMGC_MAPK1 (ERK2), we observe an unphosphorylated activation loop conformation with a distended C-terminus (PDB: 3erkA) that places the activation loop over the APE motif and in contact with the G-helix, as indicated by an expanded distance between APE12 and DFG4 (Cβ – Cα > 14Å) (**Figure 5F**). These structures, which are likely inactive, differ significantly from the phosphorylated state of MAPK1 (PDB: 2erkA) and other substrate-bound structures in the CMGC family, e.g., CDK2 (**Figure 8D**). The APE12 distance criterion proposed here designates 3erkA as inactive and 2erkA as active.

Finally, due to the flexibility of the activation loop C-terminus depicted for substrate-bound TYR kinases in **Figure 2** we forego the additional distance criteria for APE11 – APE12 for TYR kinases.

### HRD Motif Conformation

98% of human catalytic protein kinases have a conserved Asp in the αF-helix that forms hydrogen bonds with the backbone of the HRD loop, which is in a strained conformation [44]. This structural motif stabilizes the positioning of the His and Arg residues, which themselves have a regulatory function. The side chain of HRD-His donates a hydrogen bond to the backbone carbonyl of X in the **X**DFG motif when it is in the *BLAminus* conformation [10], while HRD-Arg forms complementary charge-charge interactions with phosphorylated residues of the activation loop [13]. To include this structural motif in our criteria for structurally active kinases, based on visual inspection of the Ramachandran plots for the HRD motif over substrate-bound structures in the PDB (**Figure 7D**), we restrict active kinases to examples where the His-HRD backbone is located in the alpha region and the Arg-HRD backbone is located in the L region of the Ramachandran plot: ϕ ∈ (0°,180°), ψ ∈ (−50°,100°). There is evidence from the PDB that this structural motif plays an important role in maintaining the active state: CAMK_AURKA has 64 inactive structures which are DFGinter, and the HRD Arg backbone nitrogen is flipped “out” in all these structures. This flip breaks hydrogen bonds with the αF-Asp side chain as in the DFGinter structure of AURKA, 2j50A (**Figure 5G**) [44]. Distortion of the HRD loop appears to be responsible for breaking the hydrogen bond between the HRD-His side chain and XDF carbonyl, allowing for a partial flip of the DFG motif from DFGin to DFGinter. Two structures of TYR_ABL1 (chains A and B of 2g2i), are “HRD-out”. The substrate has a Phe at the P0 position and bound ADP. While it is possible that the HRD-out state is similar to SaltBr-out in that it precludes catalytic activity but not the binding of substrate, it is notable that the only HRD-out substrate-bound structures are bound to a substrate-mimic (Phe) which fails to satisfy catalytic interactions with the HRD motif. This suggests that the HRD-in backbone conformation may be favored structurally by the interaction of HRD-Asp with substrate hydroxyl atoms.

The αF-Asp residue is an aspartic acid in every human catalytic kinase except for: CAMK_PIM1, CAMK_PIM2, CAMK_PIM3, OTHER_HASPIN, OTHER_PKDCC, TYR_AATK (LMTK1), TYR_LMTK2, and TYR_LMTK3. PIM1-3 have an Ala or Thr substituted at this position, meanwhile the other kinases in this list have Asn. From the PDB we observe that active structures of PIM1 and PIM2 contain a pair of water molecules which occupy the cavity formed by the substitution with Ala (as in PIM1, PDB: 2bzkB) or Thr (as in PIM2, PDB: 4x7qB). These water molecules appear to form hydrogen bonds with the HRD backbone analogous to the hydrogen bonds formed by the carboxylate side chain of αF-Asp.In PDB structures of the PIM family kinases, the His and Arg backbone dihedrals are similar to all other active structures (A-L regions of the Ramachandran map). Nevertheless, we forego the HRD dihedral angle criteria for all kinases that do not have the conserved Asp at the αF helix.

### Regulatory spine

Finally, we evaluated the utility of the regulatory spine for identifying active structures. The regulatory spine consists of four amino acids:

1. the His residue of the HRD motif. This residue is: His in 393 catalytic kinases; Tyr in 38 AGC kinases, CK1_CSNK1G1,2,3; OTHER_SBK2; TKL_LRRK2; Leu in OTHER_PKDCC, Phe in TKL_LRRK1.
2. the Phe residue of the DFG motif. This residue Phe except: Leu in 38 catalytic kinases; Tyr in 11 catalytic kinases; Trp in CMGC_CSNK2A1,2,3; Met in CMGC_CDK8,19; Val in OTHER_PBK.
3. the Glu4 residue (corresponding to CAMK_AURKA Q185), which is four positions after the conserved Glu of the N-terminal domain salt bridge. In catalytic kinases, this residue is: Leu (243 kinases); Met (114); Tyr (22); His (18); Phe (10); Ile (8); Gln (6); Cys (5); Val (4); Gly (3); Asn (2); Ser (2); Ala (2); Thr (1).
4. a usually hydrophobic residue just before the β4 strand, corresponding to L196 in CAMK_AURKA. We define this as “HPN7,” which means the seventh residue from the start of the conserved HPN motif (HPNxxx***X***), which occurs in the loop between the C helix and the β4 strand. In catalytic kinases: Leu (256); Phe (58); Tyr (50); Met (25); Val (16); Ile (15); Cys (7); Ala (6); Thr (3); Gln (1); Ser (1).

These four residues define three distances: Spine1 (HRD-His/DFG-Phe), Spine2 (DFG-Phe/Glu4), and Spine3 (Glu4/HPN7). When the residues are small or polar, there may not be a contact between the side chains and such a contact may not be necessary for constructing an active kinase structure. In **Supplementary Figure 2**, the distribution of Spine1, Spine2, and Spine3 are shown for ATP-bound and unbound structures in the *BLAminus* and other states. From all three plots, it can be observed that nearly all ATP-bound, *BLAminus* structures contain an intact spine. The only exceptions are the Spine2 distances in two ATP-bound structures of PAK4 (PDB: 7s46, 7s47) in which the C-helix is twisted by about 45° starting at the residue before the salt-bridge glutamic acid (E366). This distorts the position of the M370 side chain, which forms the Spine2 distance with DFG-Phe’s side chain. It is not known whether this distortion makes these PAK4 structures inactive, since the position of the Glu4 residue does not directly mediate contacts with ATP or the substrate.

Distributions of maximum spine distances (across Spine1, Spine2, Spine3) for kinase structures with and without ATP and in the *BLAminus* and other states is shown in **Supplementary Figure 2D**. As we described in our previous paper, a majority of structures in several DFGin conformational states (*BLAminus*, *ABAminus*, *BLBplus*, etc.) contain an intact spine (defined as having all three spine distances less than 5.0 Å). 99% of *BLAminus* structures contain an intact spine. Most of those with a broken spine occur because of the position of the HPN7 residue, which does not interact with the substrate or ATP. When we apply the 5 active criteria described in the previous sections (DFGin, *BLAminus*, SaltBr-in, ActLoopNT-in, ActLoopCT-in), there are 2451 human catalytic kinase chains in the PDB that are “Active.” There are only 14 structures in this set with a Spine3 distance greater than 6 Å, all of them PAK4 with a twisted end of the C-helix. There are only 11 active structures with a spine distance between 5 and 6 Å. For the sake of simplicity, we therefore do not use the regulatory spine as a criterion for active structures.

### Active structures of catalytic kinases in the Protein Data Bank

From the considerations above, we define probable “Active” structures of kinases as those capable of binding ATP, Mg^2+^ ions, and substrate, with the criteria summarized in Table 2. From these criteria, we can label each feature required for the kinase to be “Active”: 1) DFGin; 2) *BLAminus*; 3) SaltBr-in; 4) HRD-in; 5) ActLoopNT-in (DFG6 residue); 6) ActLoopCT-in (APE6-APE12 criteria).

**Table 2.**
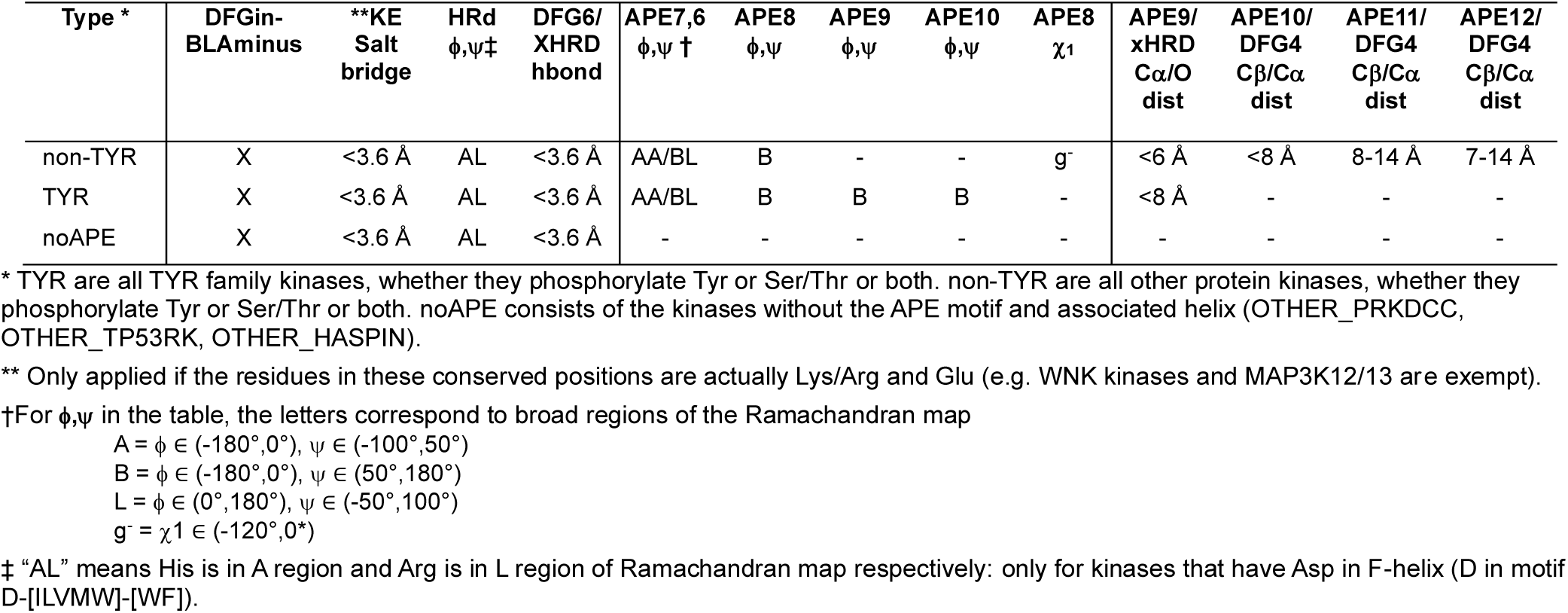
Summary of criteria for classifying protein kinase domain structures as “Active”.

We made certain exceptions to the criteria for some kinases. The salt bridge criterion is skipped for OTHER_WNK1, WNK2, WNK3, and WNK4 kinases (WNK-“With No Lysine”) and for TKL_MAP3K12 and TKL_MAP3K13. In the experimental structures of TKL_MAP3K12 (e.g., 5CEP), the residue equivalent to the salt bridge Glu is Asp161 and is turned outwards with a break in the alpha C-helix, which is shorter than that of other kinases. The presence of the Asp makes the salt bridge less likely to form so we omitted it as a criterion for these two kinases. Finally, OTHER_HASPIN, OTHER_TP53RK, and OTHER_PKDCC do not have APE motifs [10], and do not fold into the same structures as the C-terminal regions of other kinases. Thus, there is no ActLoopCT requirement for these kinases. They are listed as noAPE in Table 2.

The six conformational labels (DFGin, *BLAminus*, SaltBr-in, ActLoopNT-in, ActLoopCT-in, HRD-in) are determined by a total of 26 structural features, enabling us to apply the criteria broadly to diverse families of kinases with different activation loop sequences. This set of features still allows for substantial structural variability within ensembles of experimental or simulated structures that represent the active form of protein kinases. This comes at the potential cost of including false positives in the active set, which we refer to as “false actives”. To estimate the prevalence of false actives using our classifier, we calculated RMSD of the activation loop N-terminus and C-terminus (first 9 and last 15 residues: DFGXXXXXX and XXXXXXXXXXXXAPE) for active structures with respect to 32 kinases from **Table 1**, using one representative substrate-bound structure per-kinase. We focus on this subset of activation-loop residues as the frequency of interaction with substrates drops off significantly towards the middle region of the activation loop (**Figure 4**), and it is therefore unlikely that this outer segment of the activation loop defines kinase activity directly.

We find that only one of the kinases from **Table 1**, CMGC_CDK2, has X-ray or cryo-EM structures in the PDB which are active by our criteria and deviate from the substrate-bound structure by more than 2 Å in the N-terminal and C-terminal regions of the activation loop. These outliers comprise only 6% of the structures classified as active for this kinase: 2c4gC, 2bpmC, 2wihC, 1vywC, 2wpaC, 2bkzC, 2bkzA, 3eocA, and belong to a common active-like, unphosphorylated conformation of CMGC_CDK2 (**Supplementary Figure 3**). For the remaining 31 kinases from **Table 1** with fully resolved N-terminal and C-terminal activation loop residues, structures predicted to be active always fall within 2 Å RMSD of the substrate-bound state.

Using the criteria listed in Table 2, we labeled all human protein kinase domain structures in the PDB as “Active”, “Inactive”, or “None” to indicate missing data (e.g., missing residues). We labeled the structure as “Inactive” if any of the criteria were violated. “Active” structures must be DFGin-*BLAminus*, HRD-in, and Salt-bridge intact (or at least Chelix-in), since structures with missing atoms in these motifs cannot easily be used for further analysis (such as MD simulations). If ActLoopNT or ActLoopCT was labeled “out”, then the structure was labeled Inactive. If none of the criteria were labeled “out” but the ActLoopNT or ActLoopCT were missing atoms, then the structure was labeled “None.” The results are shown in **Table 3** for the catalytic kinases. Of the 437 catalytic kinase domains in the human proteome, only 130 (29.8%) have active structures in the PDB. Of these, only 116 have complete sets of coordinates for the backbone of the activation loop, comprising less than 30% of catalytic kinases in the human proteome.

**Table 3.** Classification of human catalytic kinase domain structures in the PDB.

|  | With or without ActLoop disorder |  |  | With no ActLoop disorder |  |  |
| --- | --- | --- | --- | --- | --- | --- |
|  | Chains | Catalytic human kinase domains | Percent (of 437) | Chains | Catalytic human kinase domains | Percent (of 437) |
| Any conformational state | 9881 | 277 | 63.4% | 5547 | 224 | 51.3% |
| DFG-in | 8490 | 261 | 59.7% | 5075 | 207 | 47.4% |
| DFG-in+BLAminus | 5401 | 214 | 49.0% | 3894 | 170 | 38.9% |
| DFG-in+BLAminus+HRD-in | 5076 | 205 | 46.9% | 3589 | 161 | 36.8% |
| DFG-in+BLAminus+SaltBr-in | 4338 | 195 | 44.6% | 3282 | 154 | 35.2% |
| DFG-in+BLAminus+ActLoopNT-in | 5224 | 206 | 47.1% | 3821 | 167 | 38.2% |
| DFG-in+BLAminus+APE6-APE7 | 4902 | 192 | 43.9% | 3606 | 155 | 35.5% |
| DFG-in+BLAminus+APE6-APE8 | 3891 | 168 | 38.4% | 2963 | 142 | 33.6% |
| DFG-in+BLAminus+APE6-APE9 | 3658 | 156 | 35.7% | 2893 | 136 | 31.1% |
| DFG-in+BLAminus+APE6-APE10 | 2555 | 118 | 27.0% | 2170 | 106 | 24.3% |
| DFG-in+BLAminus+APE6-APE11 | 2521 | 114 | 26.1% | 2161 | 105 | 24.1% |
| DFG-in+BLAminus+APE6-APE12 | 2369 | 112 | 25.6% | 2028 | 103 | 23.6% |
| Active | 2402 | 130 | 29.8% | 2180 | 116 | 26.5% |

Counts obtained for human catalytic kinases in the PDB as of January 21, 2026

Finally, for all the 10,166 labeled human protein domains in the PDB, we examined the dihedral angle distributions for the HRD residues and the APE6-APE10 residues, plotting Ramachandran maps for each residue given the activity state of the kinase (Active, Inactive, None), separately for TYR and non-TYR kinases (**Figure 9**, **Supplementary Figure 4**). Even though we used broad regions of the Ramachandran map for the HRD residues and the APE6-APE10 residues, the distributions are much narrower for the Active kinases, indicating that the various criteria are not completely independent. For example, for residue APE8, the criterion is that the backbone dihedral angles are restricted to a broad Beta region of the Ramachandran map: ϕ ∈ (−180°,0°), ψ ∈ (50°,180°). But for the active kinases, the ϕ,ψ values are much more narrowly distributed (**Figure 9**) with ψ>100° for almost all non-TYR kinases and ϕ>-100° for TYR kinases.

**Figure 9.**
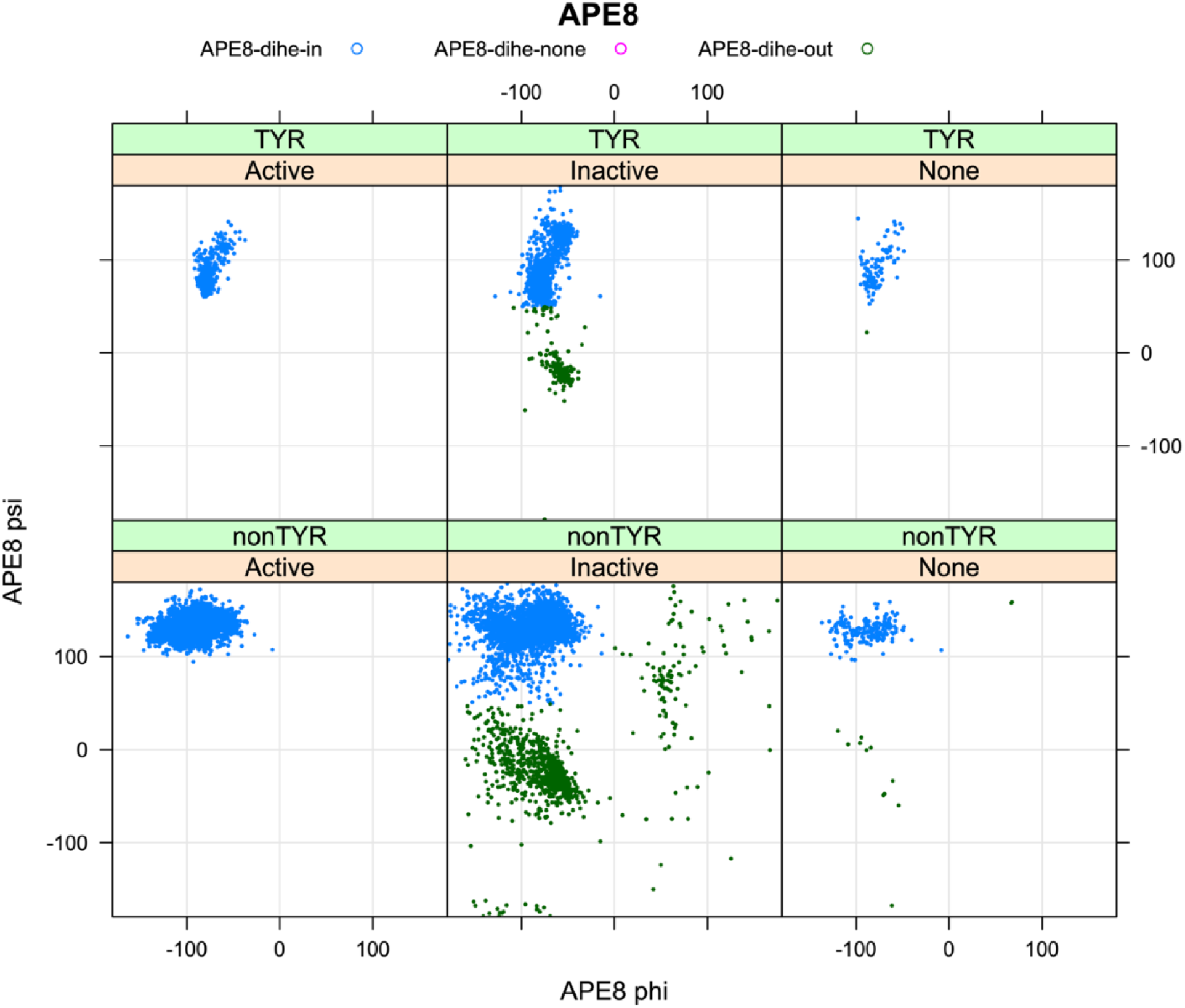
Ramachandran plots for residue APE8 of 10,166 human protein kinase domains in the PDB. The structures are divided into the TYR (top row) and non-TYR groups (bottom row) and the designation of the kinase domain as Active, Inactive, and None (missing atoms). Blue points are considered APE8-dihe-in and green points are APE8-dihe-out. Even though the criteria for APE8 is a broadly defined B region (ϕ ∈ (−180°,0°), ψ ∈ (50°,180°)), the distribution in active kinases is much more limited (compare distribution of blue dots in Active vs Inactive kinases).

### Generation of active models of catalytic protein kinase domains

To generate active models of the 437 human protein kinase domains, we created sequence sets for the multiple sequence alignments (MSAs) required by AlphaFold2 and template data sets in the active form. Sets of orthologous sequences (or near paralogs) for each kinase were created from UniProt such that each sequence in an ortholog set for a given kinase was greater than 50% identical to the target and aligns to at least 90% of the target kinase domain length with fewer than 10% gaps. Each ortholog set was filtered with CD-HIT [45] so that no two sequences in each set were more than 90% identical to each other. This was done to create diversity within the ortholog sets for each kinase. We also created “Family” sequence sets consisting of all the human kinase domains within each kinase family.

To create a set of templates, we identified all active structures of catalytic kinases in the PDB as of August 2024 (including non-human kinases) using the criteria given above and, for each kinase, selected a structure with an activation loop similar to one of the 248 substrate-bound structures (< 1.5 Å RMSD of the first five and last 15 residues of the activation loop). If more than one similar structure was available, we chose the template structure with the largest number of coordinates for the activation loop residues (to select in favor of complete activation loops). When possible, we chose a template structure which is bound to ATP or an ATP analog. This resulted in a set we named “ActivePDB”, consisting of 101 kinase domains (one structure per-kinase).

We used the Colabfold implementation of AlphaFold2 [46] to model 437 human catalytic kinase domains in the active conformation, using the ortholog and family sequence sets and each individual template from the ActivePDB set. Self-templates (where the template kinase is the same as the query kinase) were skipped to accurately assess the performance of our method. For each run of Colabfold (1 seed per run) we utilized AlphaFold2 Multimer (v3) weights. One benefit of using AlphaFold-Multimer is that all 5 models utilize provided templates (in original AlphaFold2, only models 1 and 2 use templates as input features, models 1.1.1 and 1.1.2 in Jumper et al. [18]). After each run, the models were relaxed with AMBER and the standard AlphaFold2 protocol. As a baseline, for each run of Colabfold we used a maximum MSA depth of 50 sequences. This approach yielded active models for 426 kinases in aggregate: 417 kinases using ortholog MSAs, and 415 kinases using family MSAs. The remaining 18 kinases which were resistant to adopting the active conformation at the baseline MSA depth of 50 sequences required smaller depths (25 and 10 sequences) to generate active models. Several kinases were sufficiently resistant to the active templates as to require drastically smaller MSA depths (2 or 3 sequences): TYR_AATK (LMTK1), TYR_LMTK2, TYR_LMTK3, OTHER_WEE1, OTHER_WEE2, OTHER_PKDCC, and CAMK_OBSCN-2. The kinase OBSCN-2 required additional treatment as described below.

Table 4 summarizes the results for various template and MSA sequence sources. The results of our single-template protocol (using ortholog and family MSAs) yielded active models for all but one of the 437 catalytic kinases. The only kinase for which we could not generate active models using the single-template method, AGC_MASTL, contains a ~500 residue-long insertion in the activation loop (the longest activation loop in the kinome). For AGC_MASTL, we utilized Colabfold without the use of templates and with the MSA provided by the MMseqs2 server [47] and 101 seeds (the same number of models built with templates). This produced an Active model of MASTL.

**Table 4.** Number of active catalytic kinase domains (out of 437) produced by different Template and Sequence data sources.

| Template Sources | Sequence Sources | Number of active kinases |
| --- | --- | --- |
| *EBI (PDB70)* |  | 240 |
| None | Uniref30 | 361 |
| ActivePDB | Family | 430 |
| ActivePDB | Ortholog | 435 |
| ActivePDB | Family+Ortholog | 436 |
| All | All | 437 |

For the second kinase domain of obscurin (CAMK_OBSCN-2), the C-terminal segment of the activation loop made an α helix of residues 7825-7829 in all models that blocked access to the substrate binding site. We made the mutation D7929G (residue APE9, which is conserved as Gly in 73 out of 83 catalytic CAMK kinases) which helped to unfold this helix, at an MSA depth of 2 sequences. It is possible that OBSCN-2 is a pseudokinase.

For LMTK family kinases (TYR_AATK, TYR_LMTK2, and TYR_LMTK3), all AlphaFold2 models produced from moderate to large MSA depths (50 or more sequences) formed a folded activation loop containing a strand-turn-strand motif that would be inconsistent with substrate binding. This structural motif exists in many DFGout structures of TYR kinase family members. Active models could be produced for these kinases only at significantly low MSA depths (2 or 3 sequences). We also note that LMTK2 has an asparagine residue in place of the C-helix glutamic acid residue of the N-terminal domain salt bridge. However, this kinase retains catalytic activity and has been shown to phosphorylate CFTR and other substrates involved in neuronal activity [48].

To assess the added value of our structure prediction pipeline compared with what is already available to the public, we downloaded structures of all 429 kinases from the EBI database of AlphaFold2 models (all v6 models, containing 437 catalytic domains). The EBI database contains only one model per protein, and only 240 of the 437 kinase domains (55%) contain active structures within this set according to our criteria. By comparison, when running all five models within Colabfold using default parameters, MSAs from the MMseqs2 server [47], 101 seeds, and without templates, using Uniref30 as the sequence database we obtained active models of 361 catalytic kinases (82.6%).

### Identifying the best model with interaction prediction scores of the activation loop

For each of the 437 kinases we sought to identify the active model generated by AlphaFold2 which has the most confidently predicted activation loop. We first defined an interaction confidence score for the activation loop using ipSAE (interaction prediction Score of Aligned Errors)[22]. ipSAE is a modified version of the AlphaFold ipTM score that depends on the PAEs (predicted Aligned Errors) between two sets of residues, e.g., two different proteins in a complex. Here we apply ipSAE intramolecularly to score activation loop residues *j* aligned on non-activation loop residues *i* in the kinase domain with pLDDT ≥ 60:

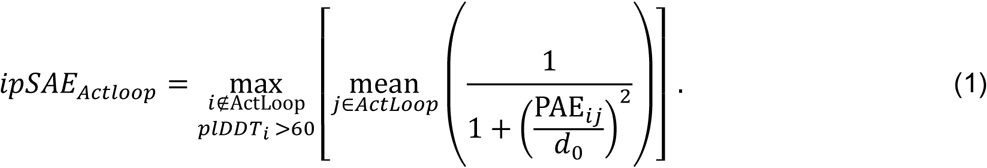

Where PAE has units of Angstroms and *d*_0_ is a scaling factor which, in typical ipSAE calculations between two interacting proteins, is a function of the length of the scored protein. Since the length of the activation loop varies from kinase-to-kinase and we wish to compare interaction scores on the same scale, we set *d*_0_ to a fixed value of 4 Å which maximizes the variance of the scores of the predicted structures (**Supplementary Figure 5**).

The experimentally determined structures of substrate-bound kinases (**Figure 2**) show that, in active kinases, the activation loop is generally situated against the kinase domain and extends from the DFG motif towards the right-edge of the kinase domain (as usually oriented and shown in **Figure 2**). It then turns around and moves leftward and concludes in the APE motif, roughly below the DFG motif. This open U shape is characteristic of substrate-bound structures and of active AlphaFold2 models produced by our pipeline. The top-scoring AlphaFold2 models produced in this study are shown in **Figure 10** for each kinase family.

**Figure 10.**
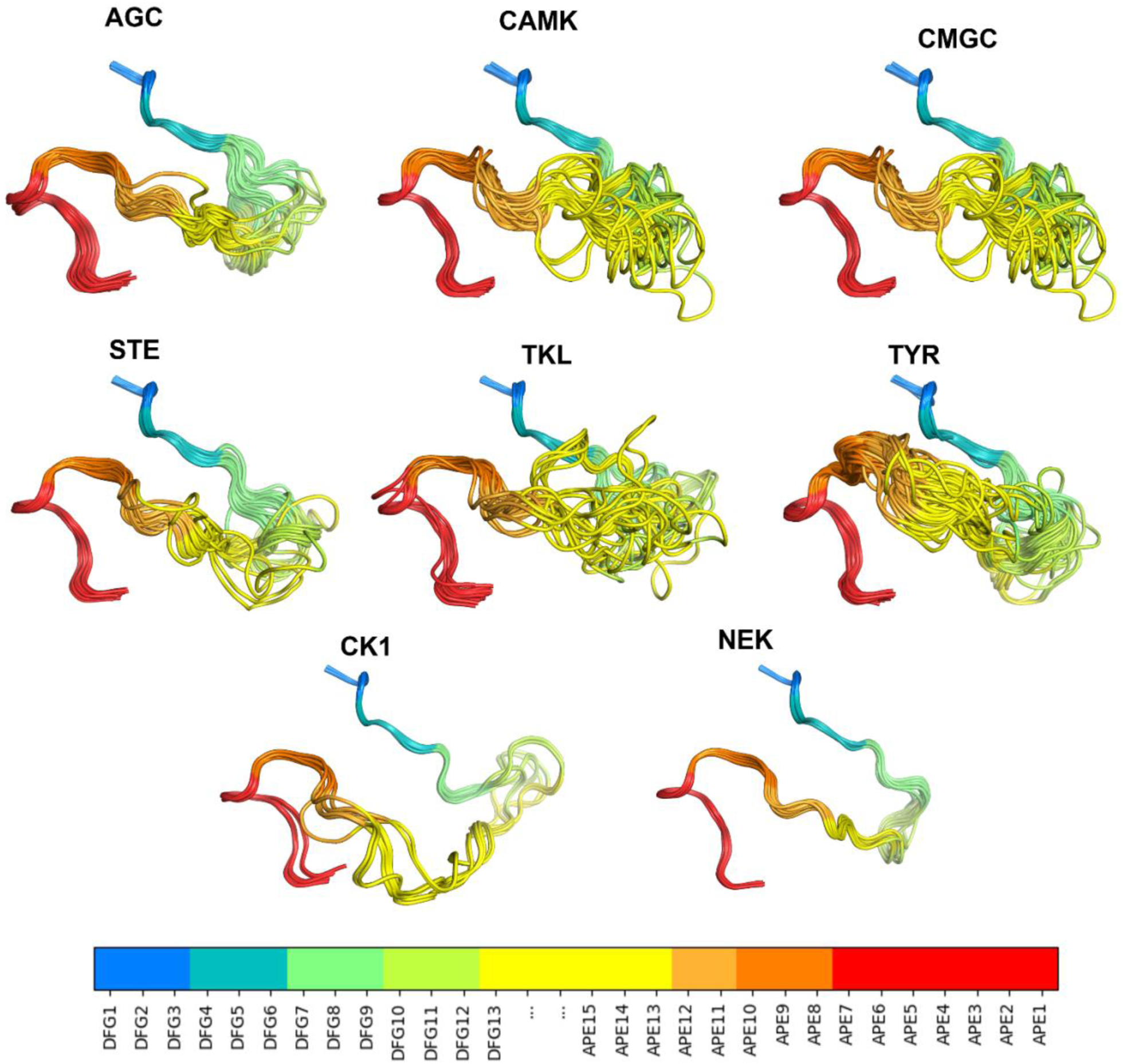
AlphaFold2 models of the active forms from each kinase family. For clarity, some structures with long disordered regions within the activation loop are not shown in each family. Diverse kinases from the group labeled “OTHER” were also excluded from the visualization. In the legend (bottom), N-terminal activation loop residues are labeled DFG*n* where *n* is the *n*^th^ residue from the beginning of the loop, starting from DFG Asp (*n*=1) and ending at DFG13 (*n*=13). C-terminal activation loop residues are labeled APE*n* in reverse order where *n* is the *n*^th^ residue from the end of the loop, starting from APE15 (*n*=15) and ending at APE1 Glu (*n*=1).

To benchmark the behavior of our pipeline in modeling active structures of catalytic kinases, we first compared the collection of AlphaFold2 models for the 29 kinases listed in **Table 1** that have complete activation loops within one of their experimental substrate-bound structures. When the same kinase is listed more than once in **Table 1**, we picked a single example since the activation loop structures were all very similar (<0.5 Å RMSD except for the two autophosphorylation structures of TKL_IRAK4). These experimental structures contain substrates and likely belong to an ensemble of substrate-binding-capable conformations of the activation loop of each kinase. Both experimental and computed structures that pass our “Active” tests still exhibit some heterogeneity of the structure of the activation loop, especially for residues far from the beginning or the end of the loop. This may be natural structural variation, and it is possible or even likely that multiple conformations are compatible with substrate phosphorylation. In any case, we explored the ability of ipSAE (Eq. 1) to pick out good models that pass our “Active” tests described above. In **Figure 11**, we show scatterplots of RMSD vs ipSAE for these 29 kinases. The results demonstrate that for most of the kinases, the model with the highest ipSAE of the activation loop also tends to correspond to the lowest or very close to the lowest RMSD to the structures listed in **Table 1**.

**Figure 11.**
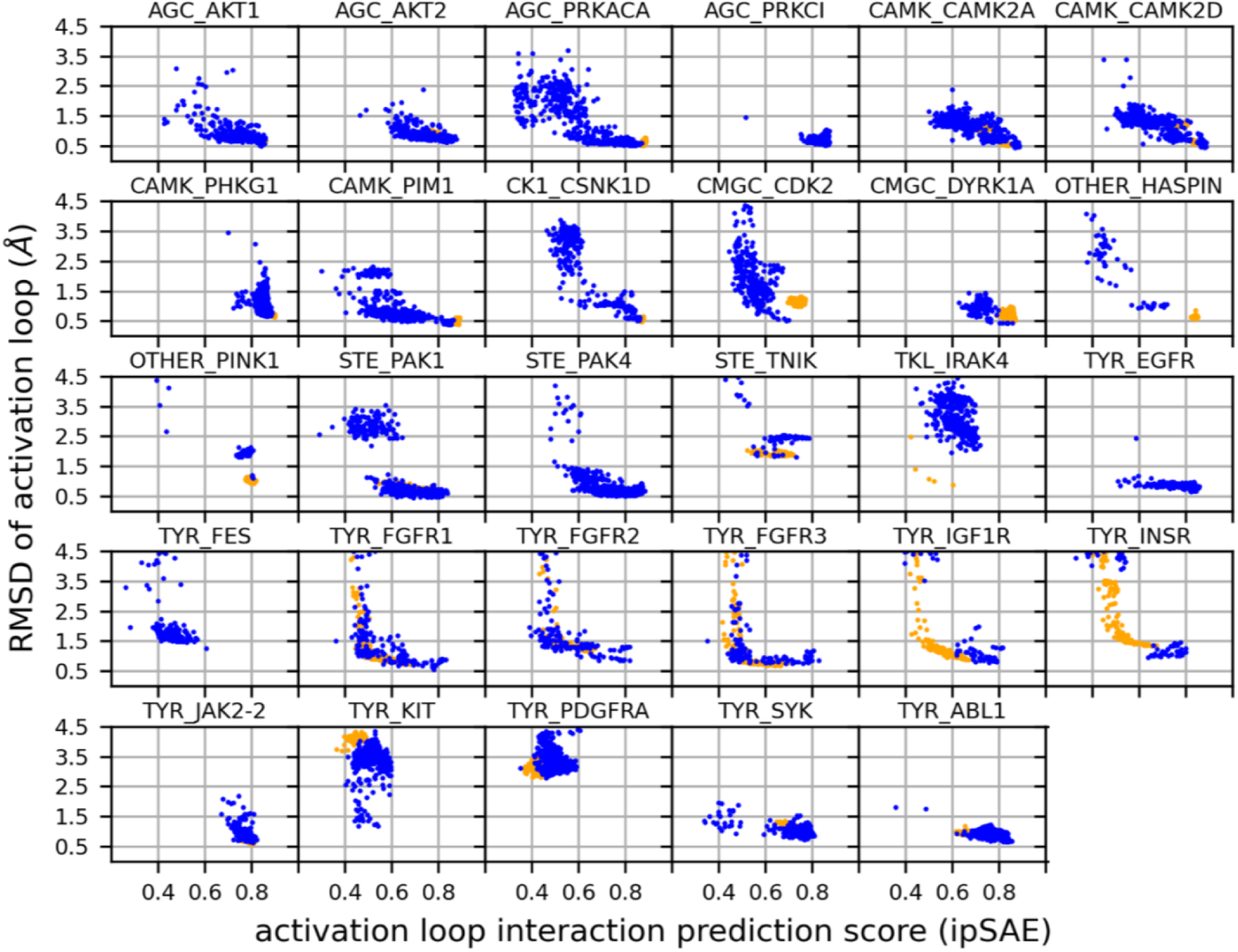
RMSD values for 29 substrate-bound structures from Table 1 versus activation loop interaction prediction score of each active AlphaFold2 model. Models from AF2 with different template modes are shown in different colors: active structures from the PDB (“ActivePDB”, blue), no templates provided to AF2 (orange). Models above 4.5 Å RMSD and intramolecular ActLoop ipSAE < 0.2 are not shown.

To extend the benchmark, we picked out at least one structure for each kinase in the August 2024 PDB with complete (or mostly complete) activation loops and extended this to orthologous kinases, resulting in 117 benchmark kinases (see Methods). The resulting benchmark set is comprised of the substrate-bound active structure (if available) or an active structure with the lowest activation loop RMSD to one of the substrate-bound structures. The distribution of RMSD to the highest scoring models (highest ipSAE score) and the distribution of ipSAE scores for these models are shown in **Figure 12**. The results show that 83 (71%) of the 117 kinases are represented by a model with less than 1 Å backbone atom RMSD (N,Cα,C,O) over the whole activation loop, after superposition of the C-terminal domains of each kinase (**Figure 12A**). A total of 108 (92%) are less than 2.0 Å.

**Figure 12.**
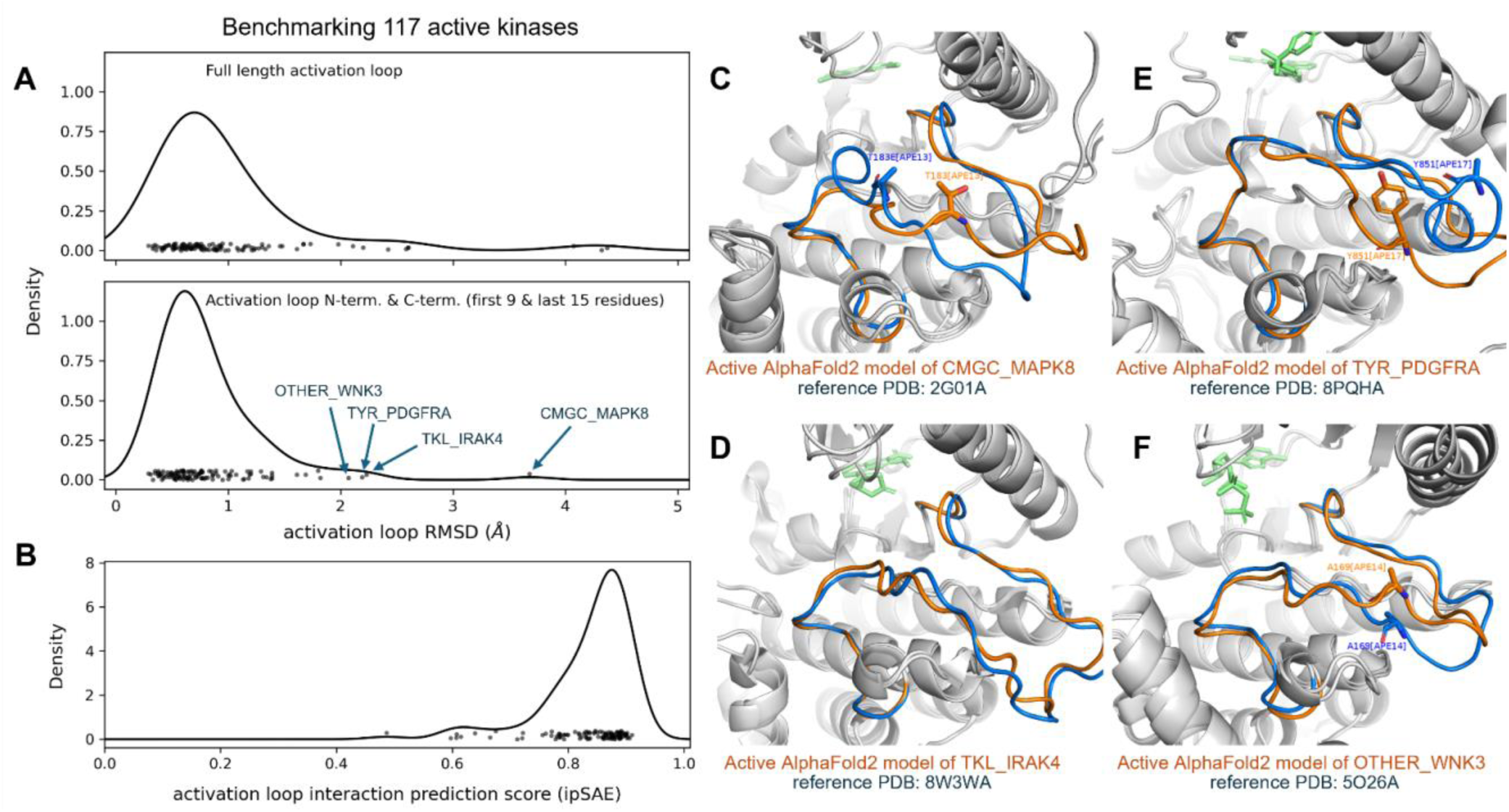
Benchmark of 117 active kinases in the PDB with full coordinates for the activation loop. **A.** Distribution of RMSD values (with kernel density estimate) for the top scoring model, (top) RMSD of the full-length activation loop, (bottom) RMSD of the first 9 N-terminal residues (DFG – DFG9) and last 15 C-terminal residues (APE15 – APE) of the activation loop. **B.** Distribution of activation loop confidence scores for these 117 AlphaFold2 models. **C-F.** Shown examples of cases where the activation loop N-term/C-term RMSD exceeds 2 Å. Activation loops of active AlphaFold2 models are shown in orange, and benchmark PDB structures are shown in blue. **C.** The CMGC_MAPK8 benchmark structure 2g01A deviates significantly from the active AlphaFold2 model, likely due to mutations made in the structure (T183E[APE13]). **D.** The AlphaFold2 modeled activation loop of TKL_IRAK4 exhibits small, evenly distributed deviations from the benchmark structure 8W3WA. **E.** The benchmark structure for TYR_PDGFRA, 8pqhA, is substrate-bound but unphosphorylated at Y841[APE17] which allows a short helix to form in the middle of the activation loop. The AlphaFold2 model, on the other hand, appears to resemble the phosphorylated state of closely related TYR kinases, e.g., FGFR2 (3gqiA) phosphorylated on Y654. In the AlphaFold2 model of PDGFRA, the equivalent residue Y841 points towards the HRD motif. **F.** OTHER_WNK3 exhibits small yet distributed deviations in the activation loop modeled by AlphaFold2, compared with the benchmark structure 5o26A.

Flexibility of the middle-region of the activation loop (between DFG9 and APE15) is expected. Conformational variability in this region is likely a natural consequence of the ensemble nature of the active state, as the frequency of interaction between activation loop and substrate drops off significantly between DFG5 and APE13 (**Figure 4**). Therefore, to conduct a more appropriate assessment of the limitations of our structure prediction pipeline we analyzed the RMSD of just the first 9 and last 15 residues of the activation loop (DFGXXXXXX and XXXXXXXXXXXXAPE), which are likely the most structurally constrained features of the active ensemble. We refer to this measure of RMSD as “NT/CT RMSD”. The results displayed in the bottom panel of **Figure 12A** show that, for 89 of the 117 benchmark kinases (76%) are within 1.0 Å of an experimental structure, and 113 of the 117 benchmark kinases (97%) our AlphaFold2 models are within 2.0 Å RMSD. The active kinase models with > 2.0 Å RMSD are: CMGC_MAPK8, TKL_IRAK4, TYR_PDGFRA, and OTHER_WNK3 (**Figure 12C-F**).

To further explore the limitations of our active criteria and scrutinize our AlphaFold2 models, we investigated each of the benchmarking outliers from **Figure 12A** as well as other examples where the AlphaFold2 model deviates from one or more active structures in the PDB by > 2 Å (ActLoop NT/CT RMSD). In total, there are 12 kinases which have at-least one structure in the PDB other than the benchmark that pass our active criteria but deviate from the corresponding predicted active AlphaFold2 model by more than 2 Å RMSD in the first 9 and last 15 residues of the activation loop. Each discrepancy between the predicted model and PDB structure may fall into one of the following cases:

**Case A:** The PDB structure is actually inactive, the AlphaFold2 model is active.

**Case B:** The AlphaFold2 model is actually inactive, the PDB structure is active.

**Case C:** Both the PDB structure and the AlphaFold2 model are active (different active states or variations within the active ensemble)

**Case D:** Both the PDB structure and AlphaFold2 model are actually inactive (different inactive states)

An overview of the 12 kinases with larger discrepancies to at least one active PDB structures is given in **Supplementary Table 5**, and the structures are shown in **Supplementary Figure 6**.

**Case A** occurs when the PDB structure is labeled active by our criteria but is likely not compatible with substrate or catalysis. Five discrepancies were assigned to **Case A**: CMGC_MAPK8, AGC_PDPK1, CMGC_CDK2, CMGC_MAPK1, and CK1_CSNK1D. For each of these discrepancies, the AlphaFold2 model resembles the active benchmark and/or substrate-bound structure, and structural deviations for one or more “active” PDB structures occur in critical substrate-binding motifs. The RMSD of our active CMGC_MAPK8 model to its benchmark PDB structure 2g01A is the most significant outlier, and upon closer examination of the structures we argue that the “active” benchmark structure of CMGC_MAPK8 (2g01A) is actually inactive. 20g1A is mutated to a phosphomimetic at the Thr[APE13] phosphorylation site (T183E) which contorts the C-terminus of the activation loop (APE13 through APE9). The activation loop of this structure has an NT/CT RMSD of 4.6 Å with respect to the active (and phosphorylated) benchmark of a closely related kinase, CMGC_MAPK1 (5v60A). In contrast, the NT/CT RMSD of 5v60A to our AlphaFold2 model of CMGC_MAPK8 is 1.9 Å, suggesting our model of CMGC_MAPK8 is likely a better representation of the active state than 2g01A. There are no wildtype structures of CMGC_MAPK8 in the PDB which pass our active criteria – all structures of this kinase that pass our active criteria are T183E mutants, and all have RMSDs greater than 2 Å from the active AlphaFold2 model. We argue that these “active-like” PDB structures of CMGC_MAPK8 are a result of noise at the boundary of our APE9 criteria: the average APE9-Cα/HRD2-O distance in these structures is 4.9 Å – an unusually large value for substrate-bound structures of CMGC kinases (which is at most 3.3 Å: see **Table 1**).

Only one discrepancy, TKL_BMPR1B, was assigned to **Case B**: in this case the active PDB structure of TKL_BMPR1B, 3mdyC, resembles the benchmark active structure of a closely related kinase (TKL_TGFBR1: 5friA) while the AlphaFold2 model of TKL_BMPR1B deviates significantly at APE10 (**Supplementary Figure 7**), which is an important substrate-binding site. This model was generated using CAMK_DAPK3 as a template. In the online dataset on the Kincore website, we replaced this model with the next highest-scoring AlphaFold2 structure (ipSAE of 0.771 to 0.769) which instead uses TKL_IRAK4 as a template, resulting in an orientation of APE10 that is more consistent with the active TGFBR1 PDB structure (**Supplementary Figure 7**).

Six discrepancies were assigned to **Case C**: OTHER_WNK3, CMGC_CLK1, CMGC_DYRK1A, TKL_IRAK4, TKL_TGFBR1, and TYR_PDGFRA, because the active AlphaFold2 model *and* PDB structures that deviate from this model by more than 2 Å both appear to be active. Structural deviations for these cases occur towards the middle of the activation loop, which are unlikely to interfere with substrate binding. We have not assigned any discrepancies to **Case D**. In summary, we identified 12 kinases with at least one structure that is labeled active but deviates from the corresponding AlphaFold2 model by more than 2 Å RMSD. Of these 12 discrepancies, 5 appear to be examples of **Case A**, 1 of **Case B**, and 6 of **Case C** (**Supplementary Table 5**).

For some other kinases, the AF2 models have poor ipSAE scores for the activation loop. This occurs for some kinases that are remotely related to other kinases in the human proteome or that have particularly long activation loops. The following 20 kinases have a top-scoring model with ipSAE < 0.5, in order of increasing score: AGC_MASTL, OTHER_CDC7, OTHER_PKDCC, AGC_LATS1, AGC_LATS2, TYR_LMTK2, TYR_NTRK3, OTHER_WEE1, OTHER_WEE2, AGC_STK38L, TYR_AATK (LMTK1), TKL_RIPK1, TYR_LMTK3, OTHER_EIF2AK4-2, AGC_STK38, TKL_LIMK2, TYR_MST1R, TYR_FRK, TKL_LIMK1, and OTHER_EIF2AK3. We note that OTHER_CDC7 bears close resemblance to the substrate-mimetic complex in **Table 1** (0.5 Å NT/CT RMSD with PDB: 6ya7A) despite its low score of 0.25.

There are only two examples of catalytic kinases which bear an unusual hydrophobic residue at the HRD-His position: TKL_LRRK1 substitutes His with Phe, and OTHER_PKDCC substitutes His with Leu. While the Phe sidechain of TKL_LRRK1 lacks hydrogen bonding capabilities, our AlphaFold2 models predict an interaction between the X[XDFG] backbone carbonyl (in the *BLAminus* state) and the “tip” of the Phe ring (Cς atom). On the other hand, OTHER_PKDCC has a Leu sidechain at the His-HRD position which is too short to interact with and stabilize the XDFG backbone in the *BLAminus* conformation according to our AlphaFold2 models. Without this stabilization of *BLAminus* provided by the HRD loop, AlphaFold tends to generate structures in the *ABAminus* conformation even when provided *BLAminus* structures as templates. In this conformation, the activation loop of OTHER_PKDCC C-terminus forms a beta sheet that folds against the αF-helix (**Supplementary Figure 8A**).

Only one *BLAminus* structure was generated by our structure prediction pipeline for this kinase, and in this case the activation loop scores very poorly. We hypothesize that the former case, where OTHER_PKDCC is in the *ABAminus* state with the activation loop folded against the αF-helix, may indeed be the active state for this kinase (**Supplementary Figure 8B**). This may be the only exception in the catalytic human kinome where the *ABAminus* conformation can be regarded as the most likely active conformation. In support of this prediction, using AlphaFold3 to generate structures of OTHER_PKDCC bound to ATP and Mg^2+^ we observe the same conformation of the activation loop and *ABAminus* state of the XDFG residues (**Supplementary Figure 8C-D**).

### Post hoc benchmarking and predicted active structure for Wee1 family kinases

Since the completion of this work, we have identified experimental structures for two new kinases that were released by the PDB in January 2026 and are classified as active using our structural criteria: OTHER_PKMYT1 (9m6pBBB) and CMGC_MAPK14 (9cj2A). These kinases did not have any experimentally determined active structures (according to our criteria) at the time of our structural bioinformatics analysis on the August 2024 PDB and modeling computations and can therefore be used to test our structural predictions *post hoc*. Our active models of these kinases closely resemble their recently solved active structures (RMSD ≈ 1.3 Å for the first 9 and last 15 residues of the activation loop).

OTHER_PKYMT1 belongs to a functionally distinct family of kinases that includes OTHER_WEE1 and OTHER_WEE2. Notably, there are no substrate-bound structures of these kinases, and they are somewhat distantly related to any structure in the ActivePDB template set. These kinases are unique in that they more closely resemble non-TYR kinases in both sequence and structure but paradoxically exhibit highly specific Tyr kinase activity (Y15 in CDK1) [49], presenting an interesting test case for our classification method outside the typical TYR vs non-TYR kinase paradigm. It is not known whether this family binds substrate like typical non-TYR families (most Ser/Thr kinases) or TYR kinases as depicted in **Figure 2**.

Due to their closer sequence homology to non-TYR kinases, for this family we apply our active criteria conditioned on substrate-bound non-TYR specific kinases. Our active model for PKMYT1 adopts the same conformation as the recent experimentally determined PKMYT1 structure, PDB: 9m6pBBB (ipSAE_Actloop_ = 0.64) [50] (**Figure 13**) which was not present in the template set for our structure prediction pipeline or AlphaFold2 training data. This is currently the only example of an experimentally determined active structure from this sub-family in the PDB. The key difference between 9m6pBBB and other PDB structures of this family (PKMYT1, WEE1, and WEE2) is the conformation of Glu-268 which is found at the APE10 position in the activation loop C-terminal segment of all three family members (E268, E477 and E394, equivalently). Glu[APE10] adopts an “APE10-in” conformation in our active models for all three kinases, as observed in 9m6pBBB (**Figure 13**). All other PDB structures of these kinases are found to adopt an “APE10-out” conformation, and we therefore classify them as inactive. It is likely that 9m6pBBB is representative of the active conformation of all three family members as predicted by our models (**Figure 13**). We reiterate however that the activation loop prediction scores (**Equation 1**) for WEE1 and WEE2 are very low (ipSAE_Actloop_ = 0.35), and the active conformation was only obtained using specific templates at very low MSA depths (three ortholog sequences each), similar to the LMTK family kinases.

**Figure 13.**
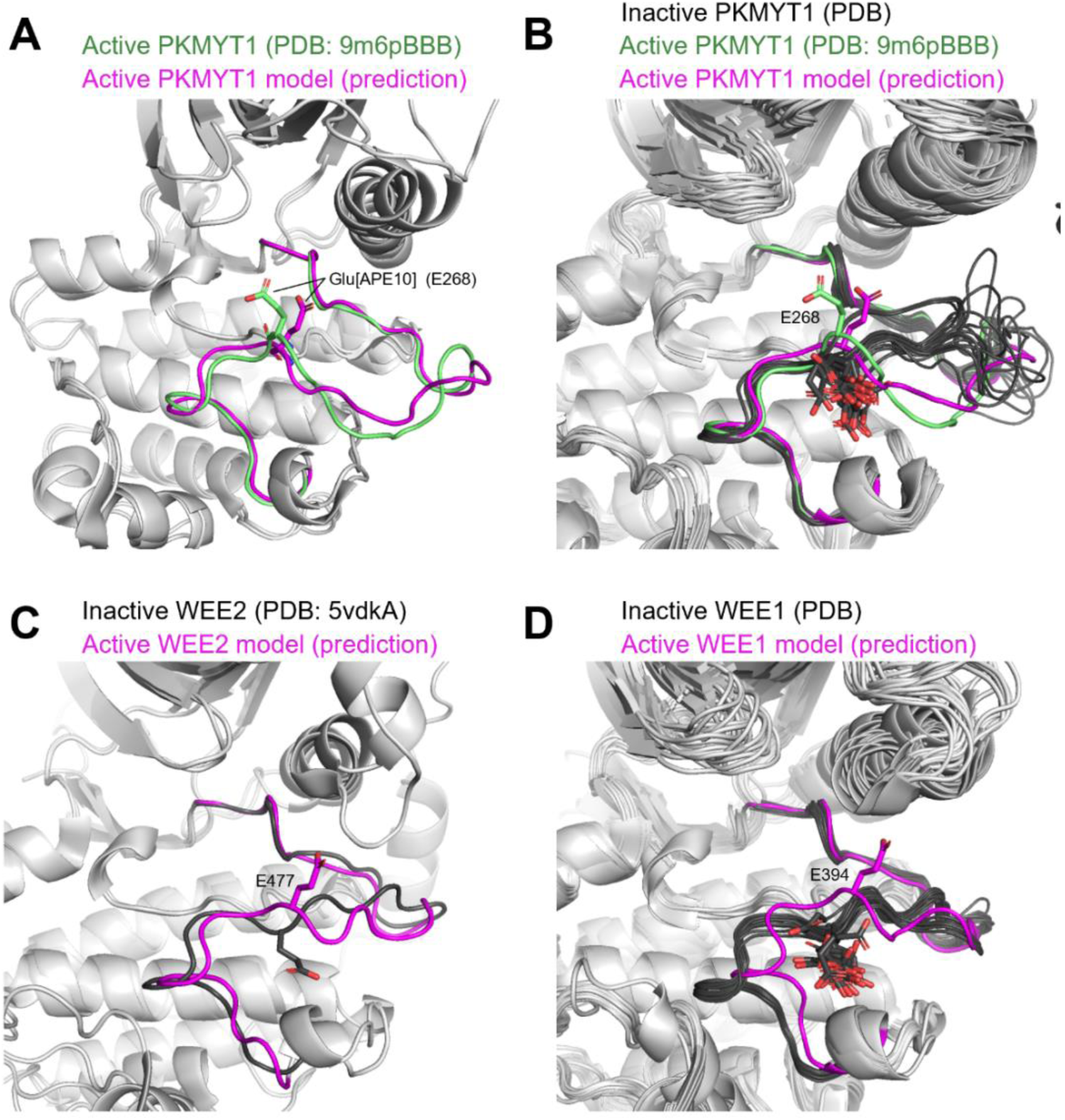
Validating the predicted active conformation of PKMYT1, WEE1 and WEE2. **A.** Superposition of a recently experimentally solved structure of OTHER_PKMYT1 (PDB: 9m6pBBB, *green*) with our predicted active model of this kinase generated with AlphaFold2 (*magenta*). The first 9 and last 15 residues of the activation loop have an RMSD of 1.3 Å. Glu[APE10] (E268 in PKMYT1) is shown with sticks; both structures are in an active “APE10-in” conformation typical of substrate-bound non-TYR kinases. **B.** All PDB structures of PKMYT1 superimposed onto our active model for this kinase (*magenta*). The recently experimentally solved active structure of PKMYT1 is shown in *green*. All other experimental PDB structures of this kinase have Glu[APE10] (E268) flipped “out” and are classified inactive/APE10-out by our criteria (*black*). The active PDB structure of PKMYT1 (*green*) validates our predicted active structure (*magenta*) and supports the classification of other PKMYT1 structures as inactive. **C.** All PDB structures of WEE2 (*black*) superimposed with our active model of this kinase which was generated with AF2 (*magenta*). All PDB structures of this kinase have Glu[APE10] (E394) flipped “out” and are classified as inactive. There is currently no experimentally determined active/APE10-in structure of WEE2 in the PDB according to our criteria. **D.** All PDB structures of WEE1 (*black*) superimposed with our active model of this kinase which was generated with AF2 (*magenta*). All PDB structures of this kinase have Glu[APE10] (E477) flipped “out” and are classified as inactive. There is currently no experimentally determined active/APE10-in structure of WEE1 in the PDB according to our criteria.

## DISCUSSION

With the advent of AlphaFold, much effort has gone into AF and similar programs to produce models of proteins and protein complexes in different states [20, 51–54]. Most of these do so by manipulating the multiple sequence alignment input to AF. For many proteins, standard AlphaFold may produce models in one state or another state but not both. Even if an ensemble of AF models can be generated with AlphaFold or its variants that fall into two or more conformational bins, it is important to be able to identify the functional characteristics of the different states for any downstream analysis. A key step in this process is establishing criteria for the desired state of the modeled proteins, especially for protein families of high biological and clinical relevance [55].

From a structural bioinformatics analysis of substrate bound typical protein kinase domains in the PDB, we have established criteria for identifying active experimental or predicted structures of protein kinases that are likely capable of binding ATP, metal ions, and substrates and catalyzing protein phosphorylation. The substrate-bound set is the largest ever compiled, consisting of 54 unique kinase-substrate pairs in 248 structures, including autophosphorylation structures that we identified in crystals [28]. Substrate binding has not typically been considered in previous efforts in identifying active structures of protein kinases.

Close inspection of the representative structures of each complex (**Table 1**) reveals substrate-binding surfaces formed by various structural elements. These structural elements are formed by residues of the catalytic loop (including the HRD motif) and residues at the N- and C-terminal ends of the activation loop (**Figure 2**). We show that the activation loop C-terminal motif (ActLoopCT) differs significantly between TYR and non-TYR kinases, and we therefore treat them somewhat differently (**Table 3**). We applied the full set of criteria to all experimental structures of kinases in the PDB and present these data on the updated Kincore website. In the PDB, as of January 21, 2026, there were active structures of 130 (29.8%) of the 437 human catalytic protein kinase domains.

Other kinase active/inactive classifiers have been developed in recent work using machine-learning [12, 56], but none of these methods have explicit criteria for binding of substrates and they were trained on “active” kinase structures incapable of binding substrates. Consequently, these classifiers often fail to classify structures as inactive that have an activation loop conformation that would clearly interfere with the substrate binding modes observed in experimental structures of closely related kinases solved to-date: e.g., CAMK_AURKA (1mq4A), CMGC_CDK2 (1okvC), CAMK_DAPK1 (1jktA), CMGC_MAPK1 (4iz5A). Our classifier is instead comprised of joint structural criteria which are based on the structural bioinformatics of substrate-bound complexes.

From the substrate bound structures, supplemented with structures of other kinases that had very similar conformations of the activation loop, we compiled a template set that we used to model all 437 human protein kinase domains, in conjunction with multiple sequence alignments (typically 50 sequences) of diverse orthologs or close paralogs to each kinase. The same criteria enabled us to identify active structures among the models produced by AlphaFold2. We demonstrated that the models with the highest intramolecular ipSAE score of the activation loop also most closely resemble substrate-bound structures of kinases in the PDB, enabling us to choose the most suitable model for public distribution via the Kincore website. In all, we generated approximately 400,000 models to identify active model structures for all 437 catalytic human kinases.

The current set of active models for all 437 catalytic typical protein kinases in the human proteome are available on the KinCoRe database (https://dunbrack.fccc.edu/kincore/activemodels). The functionality of the website has been upgraded significantly, such that tables that result from searches display all structural parameters used to determine activity (on the “advanced” pages, e.g., https://dunbrack.fccc.edu/kincore/GENE/PRKACA?advanced=1. This is true for all experimental structures currently in the PDB as well as our models. We believe our models will be useful in understanding the structural basis of kinase substrate specificity, since they place kinase substrate-binding residues of the activation loop and the active site in suitable positions for catalysis.

Of course, the true active state may be better represented by ensembles of structures. These are available for many kinases by downloading the active structures in the PDB via the Kincore website search page. Furthermore, we believe our active kinase models for all 437 human catalytic kinases serve as reasonable starting structures for molecular dynamics simulations to explore the conformational space within the active energy basin. Notably, it is not uncommon to find that the starting structures for MD simulations (intended to sample the active basin) are not actually active. For example, the inactive structure of TYR_SRC (PDB: 1y57A) is often used as the basis of molecular dynamics simulations of the *active* protein [57–59]. But the structure of chicken SRC (3dqwA,C), which is only two residues different from human SRC, much more closely resembles the structure of substrate-bound structure of the closely related ABL1 kinase domain (2g2iB). There are large differences in the C-terminal end of the activation loop that make 1y57A inactive.

Future updates to our set of models will be possible as new kinase-substrate complexes are deposited in the PDB. This will be important especially for kinases that are currently lacking structural data on the substrate-bound state, e.g., dual-specificity kinases, non-plant kinases from the TKL family, kinases from the NEK family, and several distantly related kinases designated as OTHER. We plan to add ATP-bound complexes generated with Boltz2 [60] to Kincore, using templates generated in this work. Another challenge at the forefront of kinase structural modeling is generating models of kinases in complex with various substrates. For typical protein kinases, AlphaFold-Multimer is capable of modeling substrate-bound complexes when given a peptide substrate and Uniref90 as a sequence database, but without the methods employed in this work (appropriate templates and carefully selected sequence alignments), it is not always capable of modeling the kinase activation loop in the active state. This will take additional study and implementation to develop a robust protocol that reliably makes models of kinase-substrate complexes from suitable choices of templates and multiple sequence alignments for AlphaFold-Multimer. This work is ongoing.

## METHODS

### Identifying substrate-bound structures in the PDB

Structures of kinases in complex with peptide/protein substrates (heterodimers and homodimers) were identified in the August 2024 PDB by checking each entry for intermolecular contacts between the HRD Asp delta oxygens and residues from chains in the same asymmetric unit or from neighboring asymmetric units, using the methods we described previously [28]. For crystal structures, the first step is to use crystal symmetry operators to build asymmetric units that are in contact with any protein atoms in the deposited structure. These can be in the same unit cell or neighboring unit cells. A generous heavy-atom distance cutoff of 4.5 Å was used to identify contacts of side-chain hydroxyls of potential substrates in one protein chain and the HRD Asp of any kinase chain. This search yielded a set of candidate transphosphorylation and autophosphorylation structures. As before, we compared these potential phosphorylation complexes with known peptide-substrate-bound protein kinase structures. This enables us to eliminate domain-swapped activation loop dimers, because the swapped Thr side chain in these structures approaches the HRD-Asp from below, rather than from above [28]. It is in the same position both in sequence and structure as the intramolecular hydrogen bond of APE8.

Some structures were not identified automatically by the structural search and needed to be added manually to the list of substrate-bound complexes. For example, TYR_ABL1 has two bisubstrate-complexes which have been long recognized as inactive (2g2f: DFG-out and 2g1t: “Src-like inactive”). However, the substrate-mimic structure (P0 Tyr→Phe mutant) of TYR_ABL1 bound to ADP (2g2i) has an activation loop conformation that resembles typical substrate-bound TYR structures, though the substrate-mimic Phe in each of these structures has a distance greater than 4.5 Å from the HRD Asp. 2g2i appears to be a reasonable model of the substrate-bound state of ABL1, except for the HRD motif which is HRD-out. The structure of OTHER_CDC7 bound to bisubstrate (a peptide covalently linked to an ATP analog) has historically served as a putative model of the substrate-bound state for this kinase (6ya7) and is corroborated by the structures of other Ser/Thr-specific substrate-bound kinases which display a similar “upside-down boat” conformation of the ActLoopCT. This complex was not automatically identified due to a non-standard residue code of the substrate mimic residue.

Some complexes appear many times in the PDB (e.g., CAMK_PIM1 with pimtide, OTHER_PINK1 autophosphorylation complex) which obscures the true number of unique, experimentally observed complexes. Based on the identity of enzyme kinase and substrate, we counted a total of 54 unique complexes from this dataset. Representative structures of each complex were chosen for visualization in **Figure 2** – for this we selected structures bound to ATP analogs and with complete coordinates for the activation loop, when available.

Some of the structures in **Table 1** contain kinases bound to folded proteins, and not peptides. These are either autophosphorylation complexes or bound protein substrates or inhibitor proteins. The autophosphorylation complexes involve sites in the activation loop (STE_PAK1, 4zy4; TYR_IGF1R, 3lvp; TKL_IRAK4, 4u97 and 8wtf; TYR_FGFR3, 8udu), the kinase insert loop (TYR_FGFR1, 3gqi; TYR_FGFR3, 4k33), and N or C terminal tails (CAMK_CAMK2D, 2wel; CAMK_CAMKII, 3kk8; TYR_CSF1R, 3lcd; TYR_KIT, 1pkg; TYR_EPHA2, 4pdo; TYR_JAK2, 6vnb; TYR_FGFR2, 3cly and 2pvf). The mitochondrial Ser/Thr kinase OTHER_PINK1 autophosphorylates on the β3-αC loop [61] as shown in 8uyh. For the activation loop autophosphorylation structures of TKL_IRAK4, we note the presence of both a symmetric (8wtf) and asymmetric (4u97) dimer in the PDB – this is a unique example where the activation loop of the substrate-bound enzyme adopts two different conformations, and it is likely that the asymmetric dimer 4u97, which has a more-compact activation loop structure of the enzyme monomer (resembling the fully phosphorylated structure: 8w3w), is the more biologically relevant of the two dimers. Three other complexes are with larger proteins which are either direct substrates or substrate-mimicking inhibitors or both (AGC_PRKACA:RYR2, 6mm7; AGC_PRKACA:KAP3, 3tnp; TKL_BAK1:HPAB2, 3tl8). The last of these is a plant kinase/pathogen-inhibitor complex [62]. These examples provide insights of how kinases phosphorylate amino acids in the context of folded protein domains, as opposed to intrinsically disordered regions (IDRs) [28].

### Ortholog sequence sets

We first searched UniProt for Pfams PF00069 and PF07714 to collect a set of 1.68 million sequences in UniRef100 with typical protein kinase domains. For each of 437 catalytic kinase domain sequences from our earlier alignment of all human kinase domains [3], we used PSI-BLAST [63] to get a list of the top 25,000 closest kinases to each human kinase domain. The queries used were 8 residues longer on each end of the kinase domain than our published alignment. The hit regions in the PSI-BLAST output were then filtered for sequences more than 50% identical to the query, coverage greater than 90% of the query length, and gap percentage in the alignment of less than 10%. We then applied CD-HIT [45] to create lists of orthologs (or close paralogs) with no more than 90% sequence identity to each other. These sequences were used as query databases to generate MSAs for our AlphaFold2 calculations.

### AlphaFold2/Colabfold

To predict the structures of human catalytic kinases we used the Colabfold implementation (version 1.5.0) of DeepMind’s advanced machine learning model, AlphaFold2 [18, 46]. The code was downloaded from the official Colabfold GitHub repository (https://github.com/sokrypton/ColabFold) and used to generate protein structures on a Linux-based high performance computing cluster equipped with NVIDIA L40S GPUs.

#### Data Input and Preparation

Three sets of sequence databases were used to create multiple sequence alignments: the default UniRef30 database (via the MMseqs2 server), an additional kinase family-focused sequence database (all 493 human kinases in the human proteome, separated into each family), and a kinase orthologs-focused sequence database (described above). Templates for the calculations were obtained from the August 2024 PDB set by performing an all-vs-all superposition of all kinase structures determined as active against each of the 248 substrate-bound structures, allowing us to identify active structures which are also within 1.5 Å RMSD of the substrate-bound state (based on the first 9 and last 15 residues of the activation loop), prioritizing templates which are themselves bound to substrate or an ATP analog. From this we acquired 101 template structures, limited to one template per-kinase to avoid over-representing certain kinase sequences in the template set. For each of the 437 human catalytic kinases, we ran AlphaFold2 using the 101 template structures individually (using a single chain for each template).

#### Model Configuration and Implementation

Calculations with AlphaFold2 were conducted using Colabfold[46]. The following command was used:

colabfold_batch --model-type alphafold2_multimer_v3 --num-seeds 1 --num-models 5 --templates --custom-template-path *template_folder a3m_msa_input_file output_folder/*

For model generation without templates using the default MSA from MMseqs2, the following command was used:

colabfold_batch --model-type alphafold2_multimer_v3 --num-seeds 101 --num-models 5 --templates *fasta_sequence_file output_folder/*

The multiple sequence alignment was prepared using the hh-suite package [64] and subsequently fed into the model for structure prediction. When using templates, we ran one seed using one template at a time. When using no templates, we ran Colabfold using default settings which uses MMseqs2 and the UniRef30 database [47] to generate the input sequence alignments, and 101 seeds to match the sampling provided by the template procedure.

We ran Colabfold with specific sequence data sets by providing our own custom sequence databases at runtime: Ortholog and Family for the MSA building step and individual templates to predict protein structures. When predicting the structure of a given target, we did not use “self-templates” i.e., templates of the target kinase were avoided in the template set (resulting in 100 templates deployed per-kinase target, down from 101).

### Benchmarking

The predicted structures were validated by comparing them to the benchmark PDB structures of kinases. The validation process relied on our ipSAE score [22] defined intramolecularly between the activation loop and kinase domain (**Equation 1**) and Root Mean Square Deviation (RMSD), measuring the distance between atoms in the predicted and known structures. Model structures were aligned to benchmark structures using the SVDSuperimposer() method in BioPython [65]. The alignment was performed on the C-terminal domain of each structure (40 residues C-terminal to the APE motif). RMSD was measured for the activation loop backbone atoms (N, CA, C, O) after superposition of the C-terminal domains.

Two benchmarks were constructed using the August 2024 PDB set of human kinases (plus non-human orthologs). The first benchmark contained substrate-bound structures from Table 1 with complete coordinates for the activation loop in the PDB structure (29 kinases). The second benchmark consisted of 117 structures with complete (or mostly complete) activation loops that passed our active criteria from the PDB.

While constructing the second benchmark, most kinases had multiple structures to choose from. The structure with the lowest RMSD to one of the substrate-bound structures was chosen. Structures bound to ATP or ATP analogs were prioritized when available. This resulted in a set of mostly human kinases with complete activation loops and some non-human orthologs. Only two structures from this set were missing coordinates in the activation loop: the benchmark structure for OTHER_AAK1 is missing two residues in the middle the loop (Q206 and A207) between DFG12 and APE22. The benchmark structure for TKL_LIMK1 is missing a ~16 residue-long insertion in the middle of the activation loop between DFG8 and APE18, and this part of the loop may not have a well-defined structure even when bound to substrate. In both cases, the missing residues are outside the interaction region for substrates. We note that activation loop contacts between non-TYR kinases and bound substrates drop off almost entirely at DFG7 and APE15 (Figure 4).

A third benchmark was constructed post hoc using the January 2026 PDB, which provided active structures for two human kinases not in the August 2024 set: OTHER_PKMYT1 (9m6pBBB) and CMGC_MAPK14 (9cj2A): both were found to be within 1.3 Å of the active AlphaFold2 model.

### Update to Kincore Database and Website and Kincore_Standalone3

We have updated the Kincore database and website (https://dunbrack.fccc.edu/kincore) to include the additional active criteria defined in this study (Salt bridge, HRD, ActLoopNT, ActLoopCT) for all structures in the PDB. Each structure is marked “Active” or “Inactive” or “None” (missing atoms) based on these criteria. Pseudokinases are marked “Pseudo.” We have also added the highest scoring AlphaFold2 active model for each of the human catalytic kinases to Kincore. These are labeled with the prefix “AF-” and the suffix “-K3” or “-K4” attached to the Uniprot Accession ID (e.g., the CAMK_AURKA model is AF-O14965-K3). “K4” is used for the second kinase domain of kinases that contain two kinase domains. Otherwise “K3” is used. “K1 and K2” were used for the models we produced earlier but are now marked “Active” or “Inactive” by the updated criteria presented in this paper. The website now allows the user to search for “Active”, “Inactive”, and “Pseudo” kinase structures on the main search page (https://dunbrack.fccc.edu/kincore/formSearch). We have also added a ligand-type called “ATPlike” which includes structures bound with ATP and its analogs (ACP, ANP, ADP, ACS). The updated standalone program for assessing the conformational state of protein kinases, Kincore-standalone3, is available at https://github.com/DunbrackLab/Kincore-standalone3/.

Pseudocode that describes the algorithm/pipeline used in Kincore-standalone3 is presented in the Supplementary Methods.

## Data Availability and Reproducibility

To ensure the reproducibility of our study, all data, including input sequences, ortholog sequence sets, and predicted structures, are accessible at http://dunbrack.fccc.edu/kincore/activemodels. PDB reference codes and RCSB links for all structures referenced in this paper are as follows:

[pdb_00001fin](http://rcsb.org/structure/1fin), [pdb_00001gag](http://rcsb.org/structure/1gag), [pdb_00001jkt](http://rcsb.org/structure/1jkt), [pdb_00001k3a](http://rcsb.org/structure/1k3a), [pdb_00001mq4](http://rcsb.org/structure/1mq4), [pdb_00001o6k](http://rcsb.org/structure/1o6k), [pdb_00001okv](http://rcsb.org/structure/1okv), [pdb_00001ol5](http://rcsb.org/structure/1ol5), [pdb_00001pkg](http://rcsb.org/structure/1pkg), [pdb_00001qmz](http://rcsb.org/structure/1qmz), [pdb_00001vyw](http://rcsb.org/structure/1vyw), [pdb_00001x8b](http://rcsb.org/structure/1x8b), [pdb_00001y57](http://rcsb.org/structure/1y57), [pdb_00001yi6](http://rcsb.org/structure/1yi6), [pdb_00002bkz](http://rcsb.org/structure/2bkz), [pdb_00002bpm](http://rcsb.org/structure/2bpm), [pdb_00002bzk](http://rcsb.org/structure/2bzk), [pdb_00002c4g](http://rcsb.org/structure/2c4g), [pdb_00002erk](http://rcsb.org/structure/2erk), [pdb_00002g2i](http://rcsb.org/structure/2g2i), [pdb_00002j50](http://rcsb.org/structure/2j50), [pdb_00002phk](http://rcsb.org/structure/2phk), [pdb_00002pvf](http://rcsb.org/structure/2pvf), [pdb_00002q0n](http://rcsb.org/structure/2q0n), [pdb_00002wel](http://rcsb.org/structure/2wel), [pdb_00002wih](http://rcsb.org/structure/2wih), [pdb_00002wo6](http://rcsb.org/structure/2wo6), [pdb_00002wpa](http://rcsb.org/structure/2wpa), [pdb_00002xuu](http://rcsb.org/structure/2xuu), [pdb_00003bu5](http://rcsb.org/structure/3bu5), [pdb_00003cd3](http://rcsb.org/structure/3cd3), [pdb_00003cly](http://rcsb.org/structure/3cly), [pdb_00003dgk](http://rcsb.org/structure/3dgk), [pdb_00003dqw](http://rcsb.org/structure/3dqw), [pdb_00003e5a](http://rcsb.org/structure/3e5a), [pdb_00003eoc](http://rcsb.org/structure/3eoc), [pdb_00003erk](http://rcsb.org/structure/3erk), [pdb_00003fxx](http://rcsb.org/structure/3fxx), [pdb_00003gqi](http://rcsb.org/structure/3gqi), [pdb_00003ha6](http://rcsb.org/structure/3ha6), [pdb_00003kk8](http://rcsb.org/structure/3kk8), [pdb_00003lcd](http://rcsb.org/structure/3lcd), [pdb_00003lvp](http://rcsb.org/structure/3lvp), [pdb_00003mdy](http://rcsb.org/structure/3mdy), [pdb_00003qhr](http://rcsb.org/structure/3qhr), [pdb_00003tl8](http://rcsb.org/structure/3tl8), [pdb_00003tnp](http://rcsb.org/structure/3tnp), [pdb_00003x2u](http://rcsb.org/structure/3x2u), [pdb_00004dee](http://rcsb.org/structure/4dee), [pdb_00004ekk](http://rcsb.org/structure/4ekk), [pdb_00004iz5](http://rcsb.org/structure/4iz5), [pdb_00004jdh](http://rcsb.org/structure/4jdh), [pdb_00004k33](http://rcsb.org/structure/4k33), [pdb_00004ouc](http://rcsb.org/structure/4ouc), [pdb_00004pdo](http://rcsb.org/structure/4pdo), [pdb_00004r3p](http://rcsb.org/structure/4r3p), [pdb_00004rio](http://rcsb.org/structure/4rio), [pdb_00004u97](http://rcsb.org/structure/4u97), [pdb_00004x7q](http://rcsb.org/structure/4x7q), [pdb_00004xj0](http://rcsb.org/structure/4xj0), [pdb_00004zy4](http://rcsb.org/structure/4zy4), [pdb_00005c27](http://rcsb.org/structure/5c27), [pdb_00005czh](http://rcsb.org/structure/5czh), [pdb_00005czi](http://rcsb.org/structure/5czi), [pdb_00005dt3](http://rcsb.org/structure/5dt3), [pdb_00005eg3](http://rcsb.org/structure/5eg3), [pdb_00005fg8](http://rcsb.org/structure/5fg8), [pdb_00005fri](http://rcsb.org/structure/5fri), [pdb_00005g1x](http://rcsb.org/structure/5g1x), [pdb_00005li1](http://rcsb.org/structure/5li1), [pdb_00005lih](http://rcsb.org/structure/5lih), [pdb_00005lxm](http://rcsb.org/structure/5lxm), [pdb_00005oro](http://rcsb.org/structure/5oro), [pdb_00005os0](http://rcsb.org/structure/5os0), [pdb_00005v60](http://rcsb.org/structure/5v60), [pdb_00005vdk](http://rcsb.org/structure/5vdk), [pdb_00005w84](http://rcsb.org/structure/5w84), [pdb_00006cqd](http://rcsb.org/structure/6cqd), [pdb_00006e2m](http://rcsb.org/structure/6e2m), [pdb_00006i2u](http://rcsb.org/structure/6i2u), [pdb_00006mm5](http://rcsb.org/structure/6mm5), [pdb_00006mm7](http://rcsb.org/structure/6mm7), [pdb_00006r49](http://rcsb.org/structure/6r49), [pdb_00006r4d](http://rcsb.org/structure/6r4d), [pdb_00006ru7](http://rcsb.org/structure/6ru7), [pdb_00006vnb](http://rcsb.org/structure/6vnb), [pdb_00006vpg](http://rcsb.org/structure/6vpg), [pdb_00006w3b](http://rcsb.org/structure/6w3b), [pdb_00006w3c](http://rcsb.org/structure/6w3c), [pdb_00006wlx](http://rcsb.org/structure/6wlx), [pdb_00006wly](http://rcsb.org/structure/6wly), [pdb_00006ya7](http://rcsb.org/structure/6ya7), [pdb_00007e12](http://rcsb.org/structure/7e12), [pdb_00007s46](http://rcsb.org/structure/7s46), [pdb_00007s47](http://rcsb.org/structure/7s47), [pdb_00007ujp](http://rcsb.org/structure/7ujp), [pdb_00007w5o](http://rcsb.org/structure/7w5o), [pdb_00007xzr](http://rcsb.org/structure/7xzr), [pdb_00007ztl](http://rcsb.org/structure/7ztl), [pdb_00008d7n](http://rcsb.org/structure/8d7n), [pdb_00008pqh](http://rcsb.org/structure/8pqh), [pdb_00008qgy](http://rcsb.org/structure/8qgy), [pdb_00008udu](http://rcsb.org/structure/8udu), [pdb_00008uyh](http://rcsb.org/structure/8uyh), [pdb_00008wtf](http://rcsb.org/structure/8wtf), [pdb_00008yhw](http://rcsb.org/structure/8yhw).

## ACKNOWLEDGMENTS

This work was funded by NIH Grant R35 GM122517 (to RLD) and P30 CA006927 (to Fox Chase Cancer Center). Calculations were performed on the Fox Chase High Performance Computer Cluster.

## Supplementary Material

**Supplementary Figure 1.**
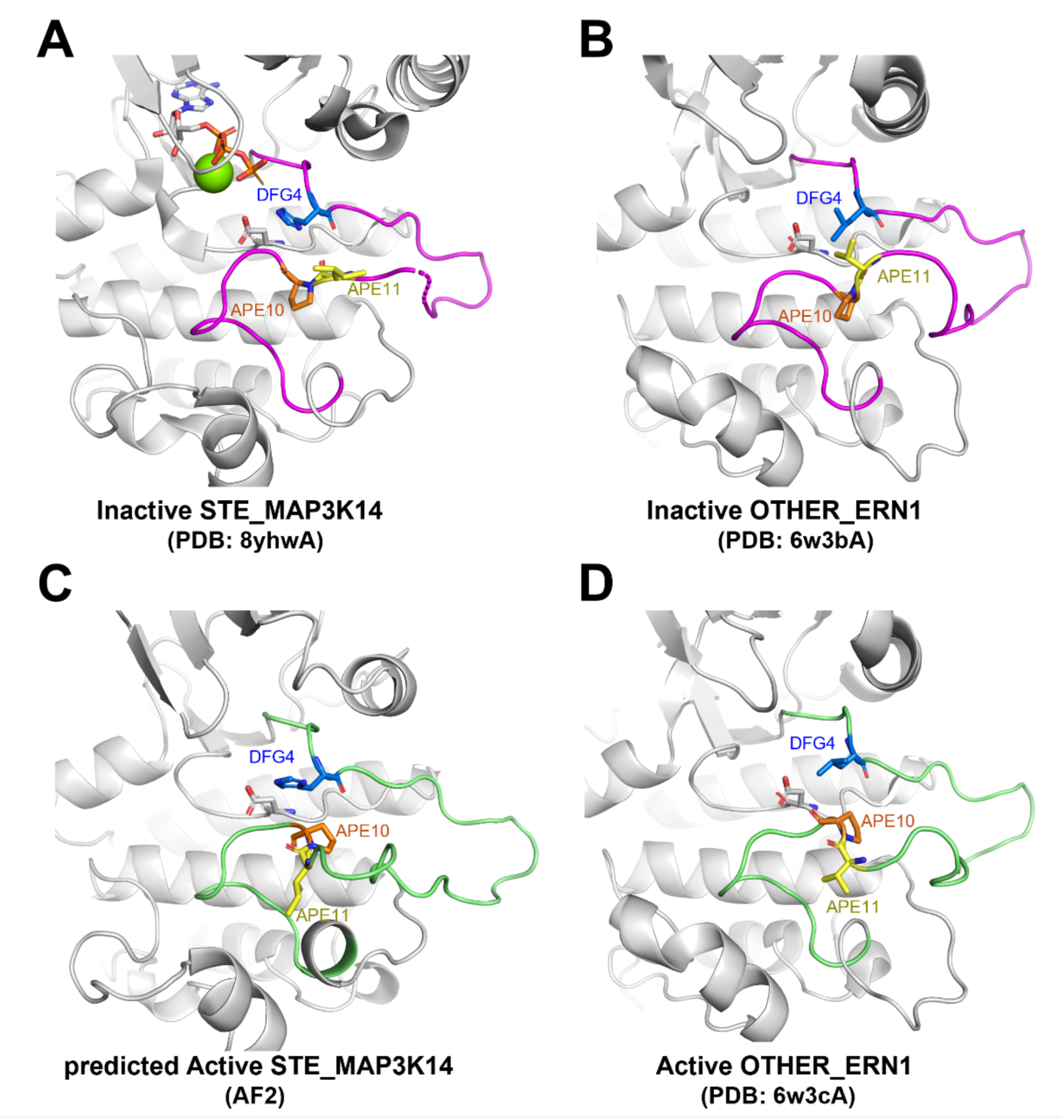
Justification for designating STE_MAP3K14 structures as inactive. Activation loops of inactive structures are colored magenta and those of active structures are colored green. The HRD Asp residue and ATP analogs (where applicable) are shown with grey sticks. **A.** Inactive STE_MAP3K14 structures have a contracted APE11-DFG4 distance that results from a flipped conformation of the Ile[APE11]-Pro[APE10] (I556 and P557) residues. We label all experimental MAP3K14 structures to-date as inactive due to this feature. B. Inactive/unphosphorylated OTHER_ERN1 structures also have a contracted APE11-DFG4 distance which results from a flipped conformation of the Val[APE11]-Pro[APE10] (V731 and P732) residues, similar to MAP3K14. **C.** AlphaFold2 model of STE_MAP3K14 generated using Colabfold with no templates (see *Methods*) and ipSAE_Actloop_ = 0.65 (see Equation 1 in main text), which is the highest-scoring model of STE_MAP3K14 generated in this study that can be classified as active. This model is also displayed in Figure 10 of the main text alongside other STE family members. **D.** An active/phosphorylated experimental structure of OTHER_ERN1 (PDB: 6w3cA) displays the typical upside-down round-bottom boat structure in its ActLoopCT that is common for non-TYR kinases despite having Pro at APE10, suggesting that MAP3K14 should also be able to adopt a similar active conformation consistent with the predicted Active MAP3K14 structure in **C**.

**Supplementary Figure 2.**
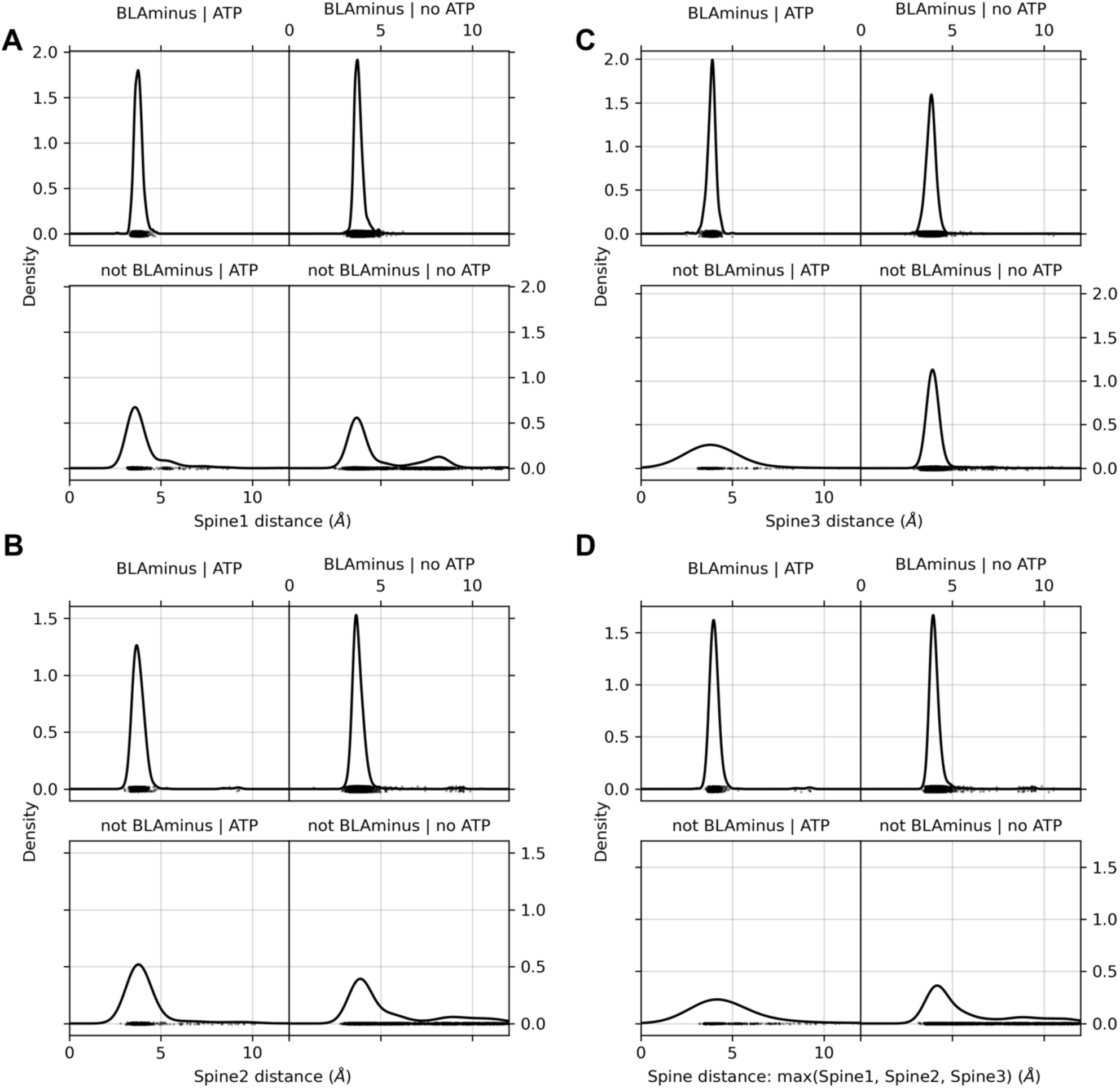
Distribution of regulatory spine distances. The Spine distances are defined as the closest distance among all side-chain atom pairs between the two residues. In each plot, there are 665 *BLAminus*/ATP points, 5043 *BLAminus*/no_ATP points, 454 not_*BLAminus*/ATP points, and 4316 not_*BLAminus*/no_ATP points.

**Supplementary Figure 3.**
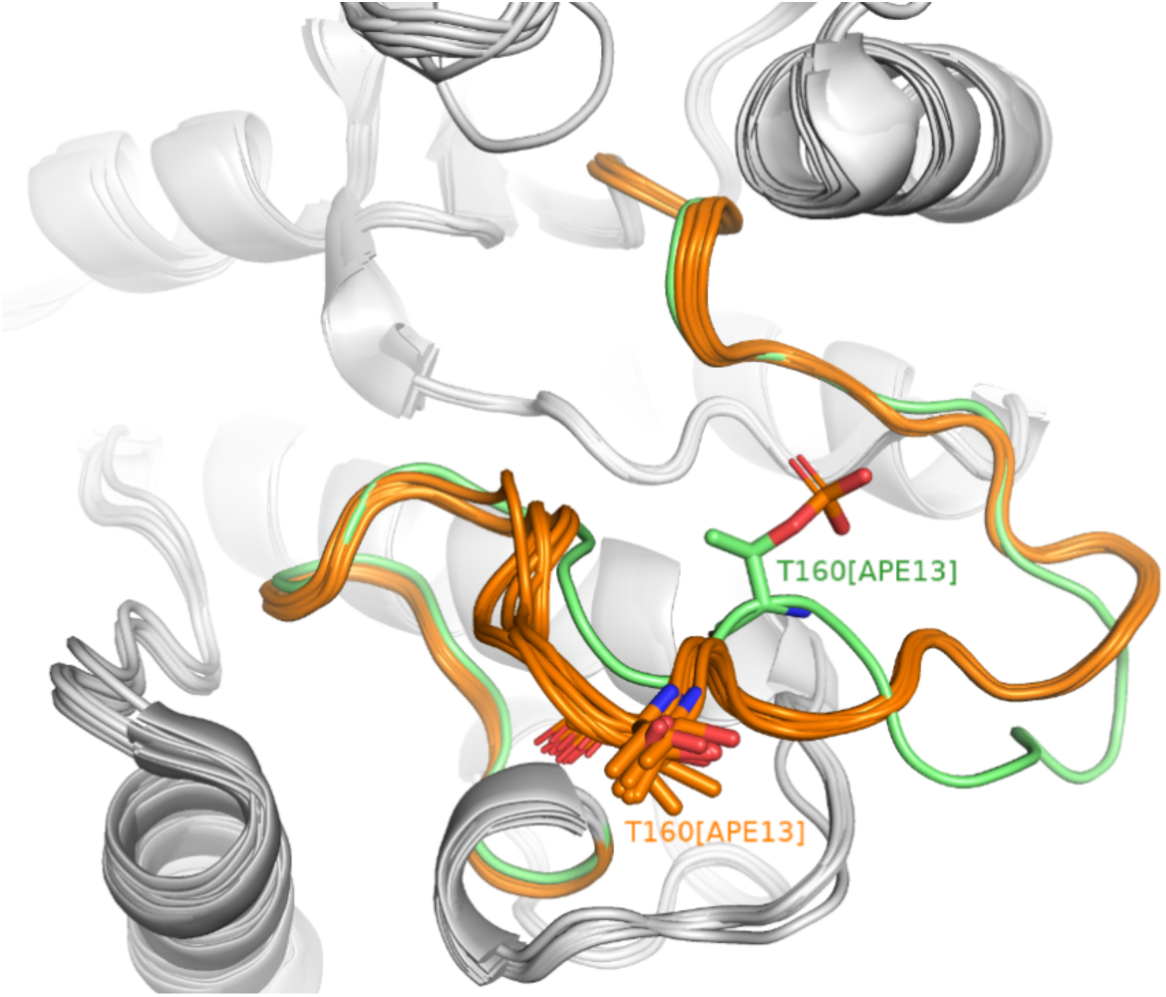
Outlier structures of CMGC_CDK2 that pass our active criteria. Among CMGC_CDK2 structures which are active by our criteria, 6% have activation loops that deviate from the substrate-bound structure by more than 2 Å RMSD. These outlier activation loops are colored orange (2c4gC, 2bpmC, 2wihC, 1vywC, 2wpaC, 2bkzC, 2bkzA, 3eocA), and the substrate-bound activation loop is colored green (1qmzA). In the outlier structures, T160[APE13] is unphosphorylated and flipped “out”.

**Supplementary Figure 4.**
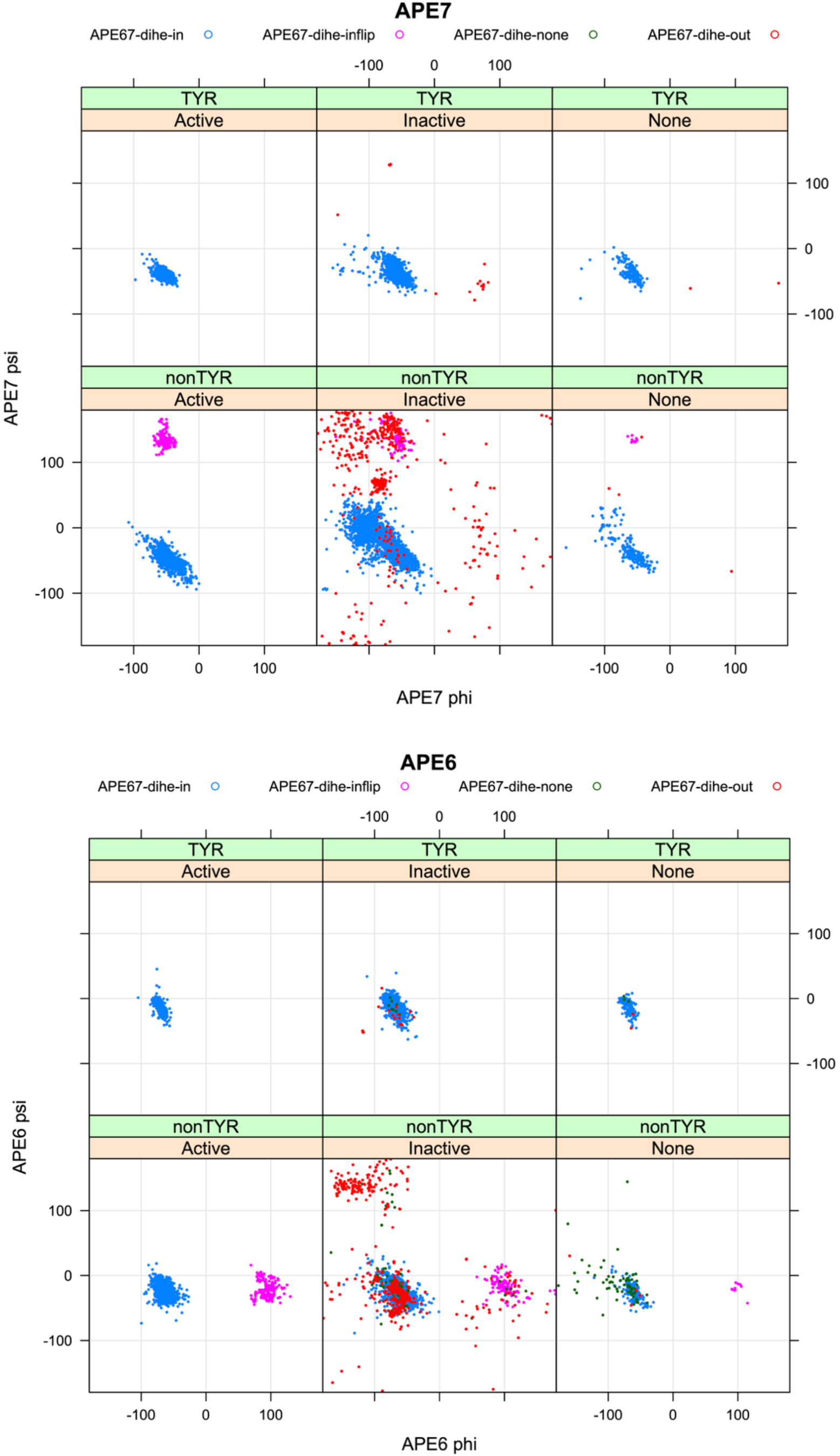
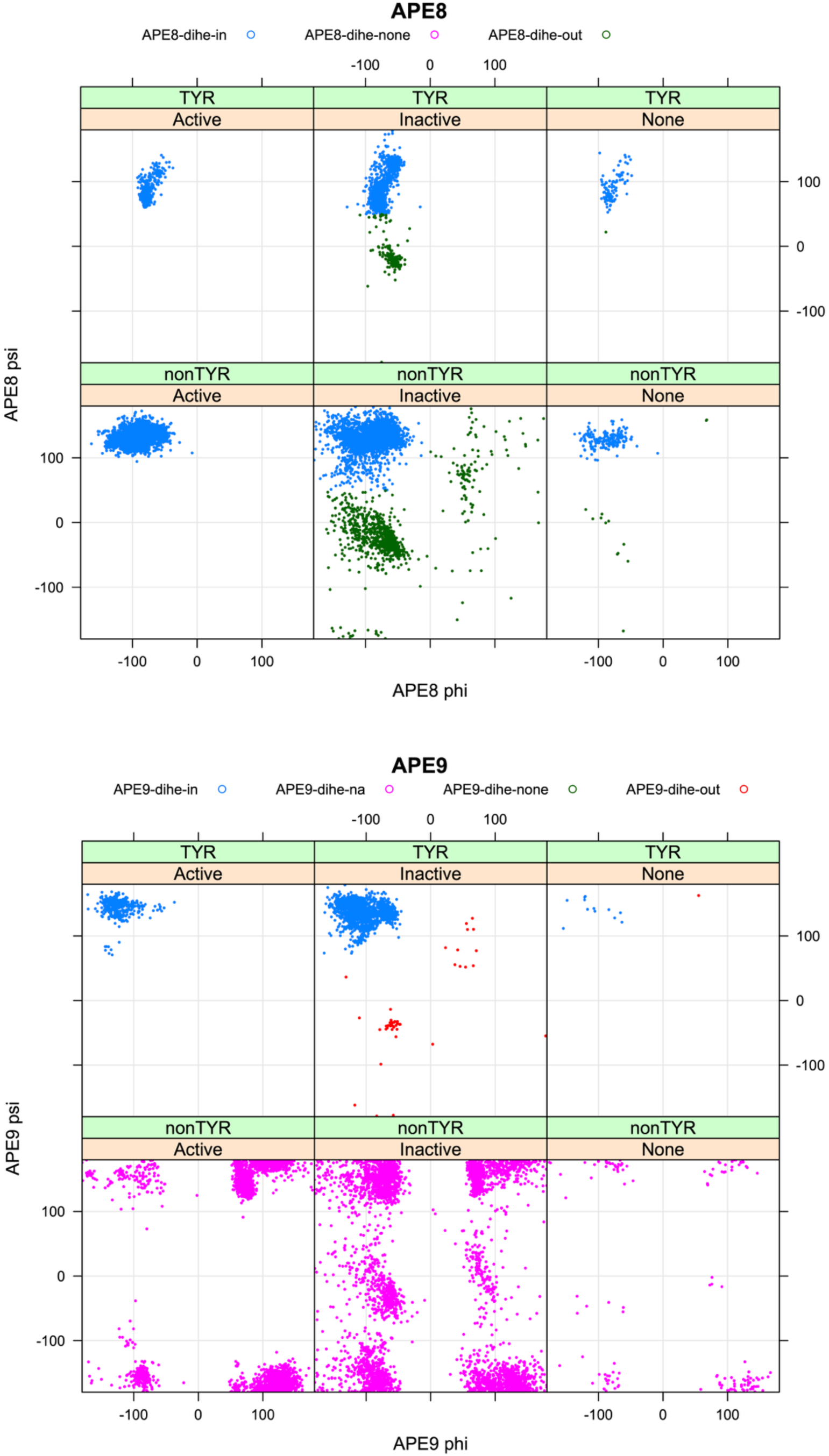
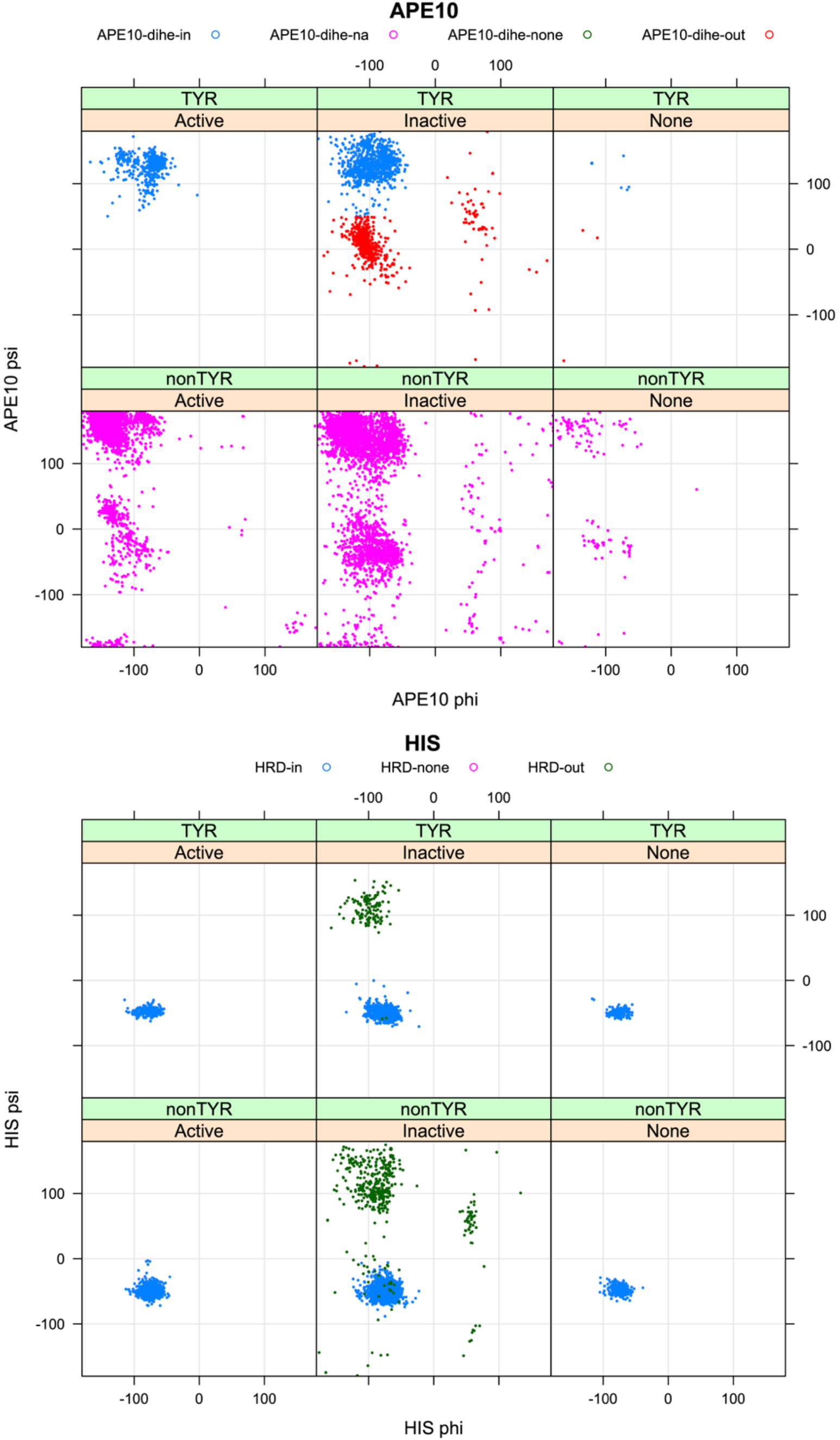
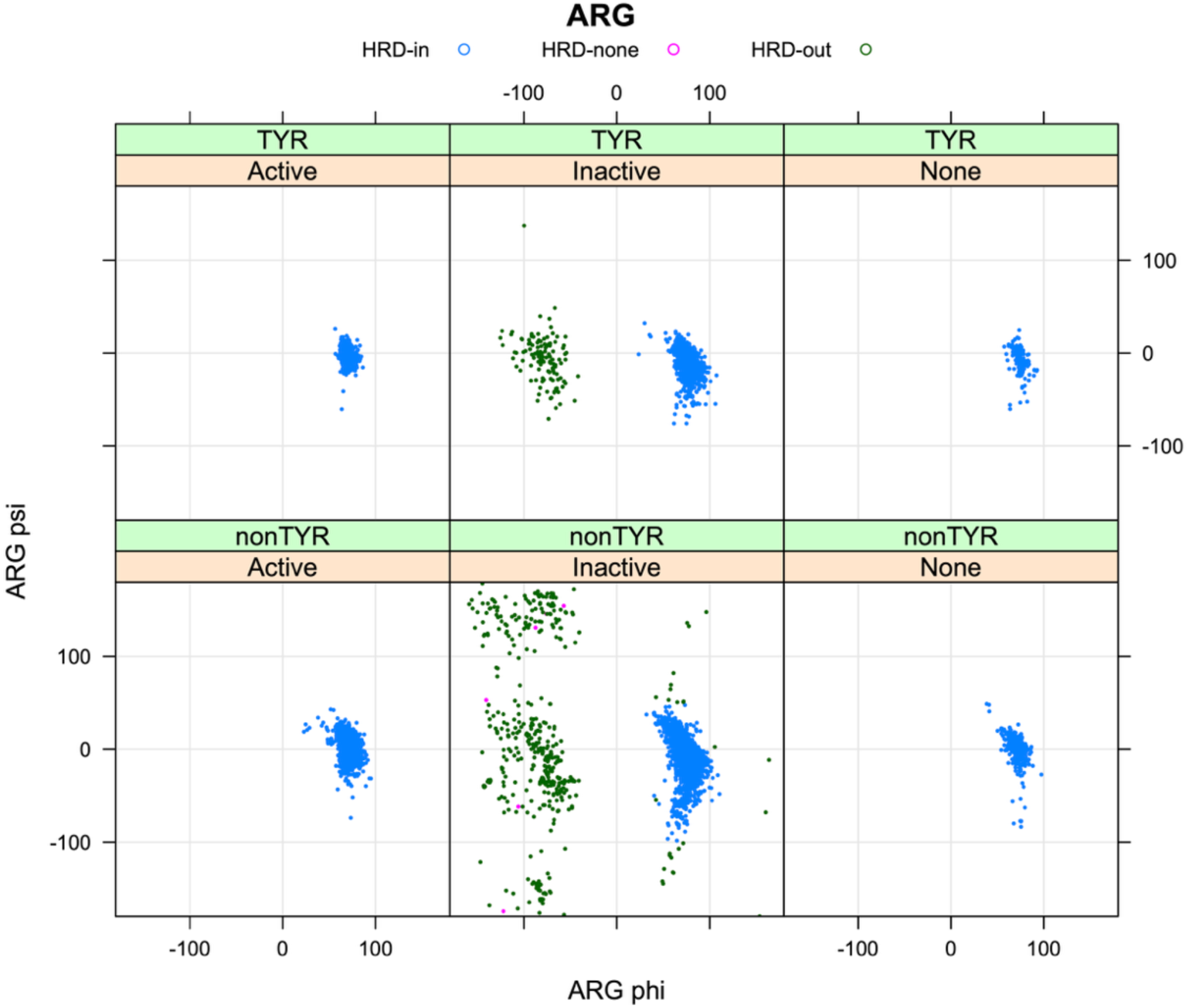
Ramachandran distributions for all human kinase domains in the PDB for residues APE6-APE10 and the His and Arg residues of the HRD motif. Distributions are considerably narrower than the broad ranges defined in the rules for labeling active structures. Each plot shows TYR (top row) and non-TYR (bottom row) kinases separately. The backbone-dihedral criteria are not used for non-TYR kinases in residues APE9 and APE10. For APE6-APE7, the state “APE67-dihe-inflip” indicates accepted structures with a peptide flip (APE7-APE6) conformations of B-L, compared to unflipped structures (“APE67-dihe-in”) which are A-A.

**Supplementary Figure 5.**
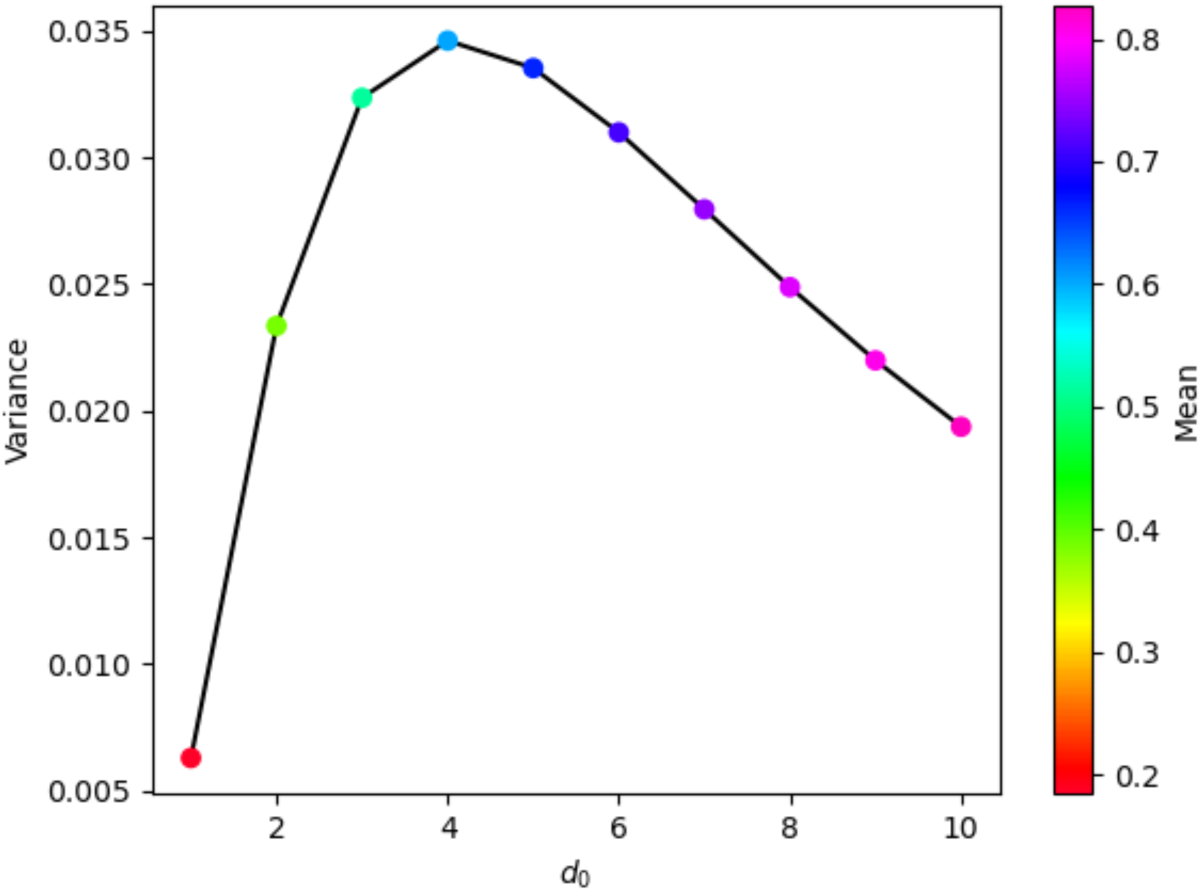
Variance and mean of ActLoop ipSAE at different values of *d*_0_. The variance and mean of ActLoop interaction prediction Scores based on Aligned Errors (ipSAE) were calculated from all 400,000 AlphaFold2 models generated in this study at different values of *d*_0_, with a maximum variance at *d*_0_ ≈ 4 Å and a mean value of approximately 0.6. The value of 4 Å was used for *d*_0_ in all activation loop ipSAE calculations reported in this study.

**Supplementary Figure 6.**
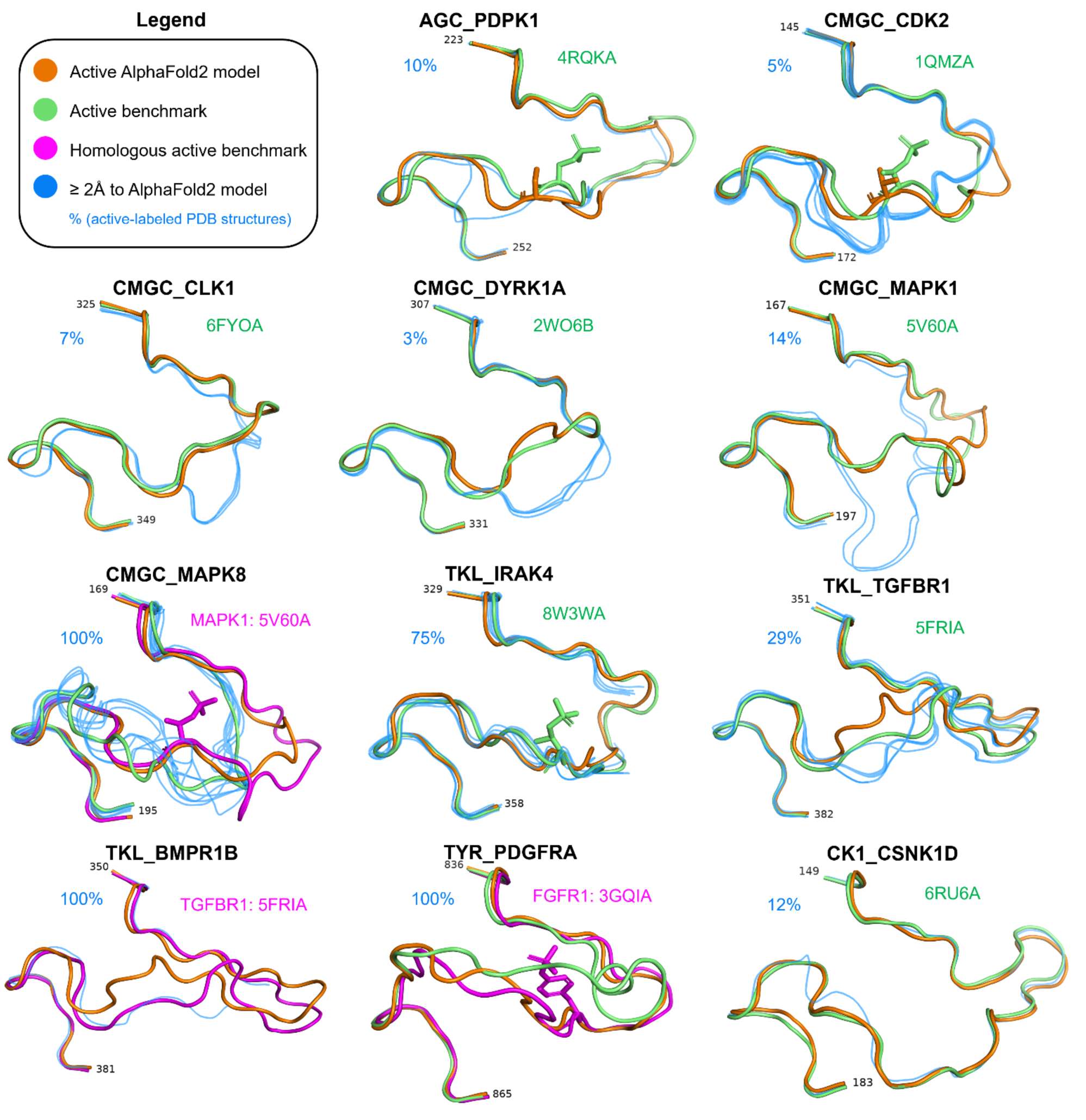
Twelve discrepancies between the active AlphaFold2 models and any of their “active” PDB structures. For each kinase displayed, at least one experimental structure labeled active by the criteria described in this paper can be found which also deviates from the active AlphaFold2 model (orange) in the first 9 or last 15 residues of the activation loop by more than 2 Å. PDB structures which are labeled active but exceed this 2 Å threshold are colored blue, benchmark active structures are colored green and homologous active benchmarks are colored magenta. Also displayed in each panel is the percentage of predicted-active structures which deviate from the active AlphaFold2 model by more than 2 Å. The benchmark structure for CMGC_MAPK8 (green) is likely incorrectly labeled as active – most likely the CMGC_MAPK1 structure 5v60A (pink) is a better active benchmark (see discussion of **Figure 12C** in the main text). For TYR_PDGFRA, the benchmark structure (green) is the only experimental PDB structure with a fully intact activation loop and differs from the active AlphaFold2 model by more than 2 Å (see **Figure 12E** in the main text) – the PDGFRA AlphaFold2 model more closely resembles the phosphorylated/active TYR_FGFR1 benchmark structure 3gqiA (pink).

**Supplementary Figure 7.**
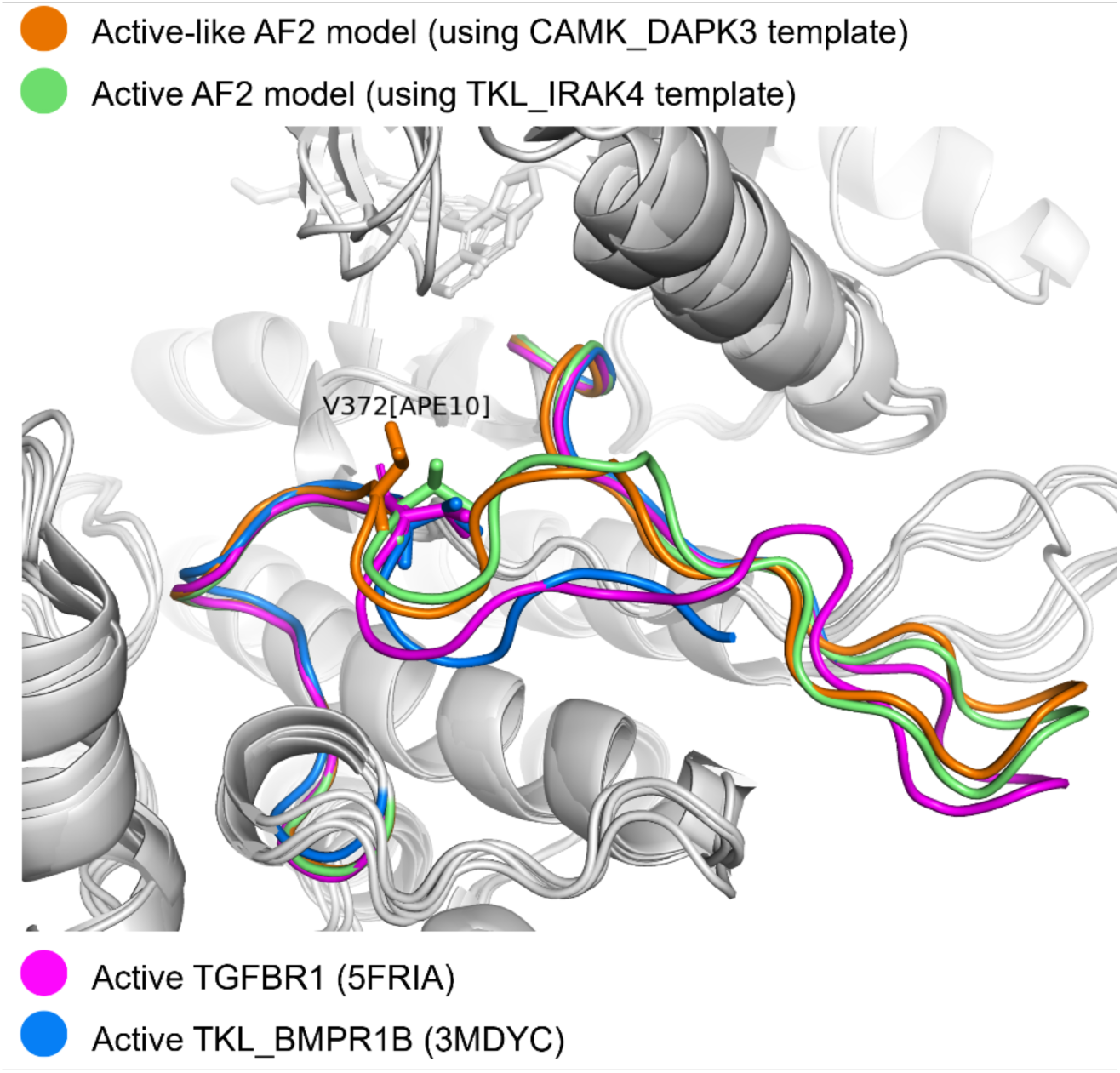
Comparison of TKL_BMPR1B models with related PDB structures. AlphaFold2 models of TKL_BMPR1B are shown in orange and green. Both models have ipSAE_Actloop_ = 0.77, but the model acquired when using TKL_IRAK4 (PDB: 8W3WA) as a template has a conformation of V372[APE10] that is more compatible with substrate binding.

**Supplementary Figure 8.**
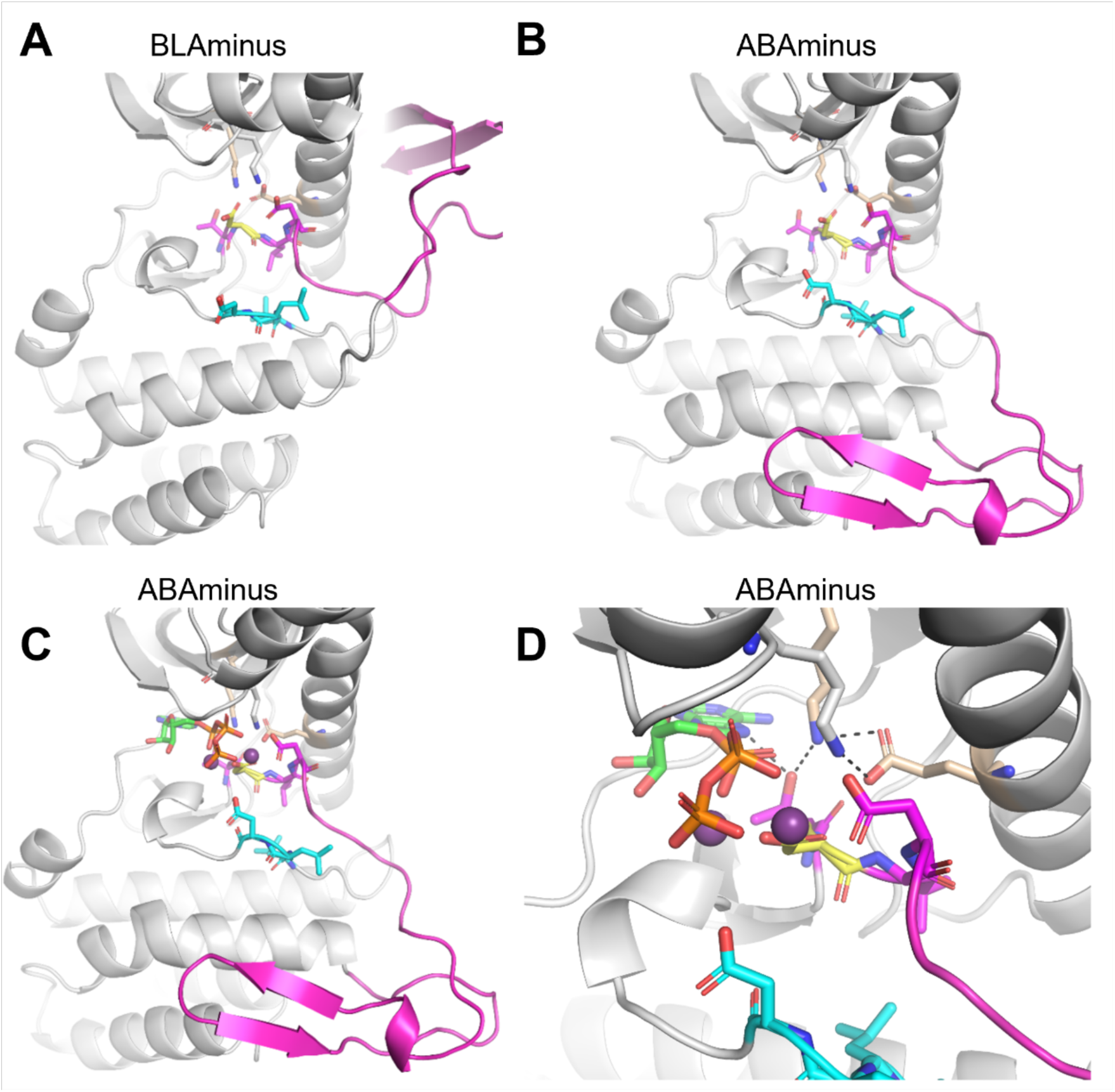
OTHER_PKDCC in a hypothesized active ABAminus conformation. **A.** The *BLAminus* conformation generated by AlphaFold2 scores poorly (ipSAE_Actloop_ = 0.27). The sequence of the DFG motif in PKDCC is DLD. **B.** The *ABAminus* conformation scores well (ipSAE_Actloop_ =0.84), which we hypothesize to be the active conformation of OTHER_PKDCC. **C.** AlphaFold3 models of OTHER_PKDCC bound to ATP·2Mg^2+^ adopt the *ABAminus* conformation. **D.**Closeup view of the Alphafold3 model of OTHER_PKDCC and its interactions with ATP in the *ABAminus* conformation. The X of the XDFG (sequence TDLD in PKDCC) and DFG3[Asp] are colored magenta, the β3-αC salt bridge residues are colored tan, the DFG1[Asp] is colored yellow and the HRD (in this case LVD) is colored cyan. An extra lysine in the β2 strand is also shown, which is normally seen in the WNK family and appears to stabilize the DFG and αC-Glu.

**Supplementary Table 1.**
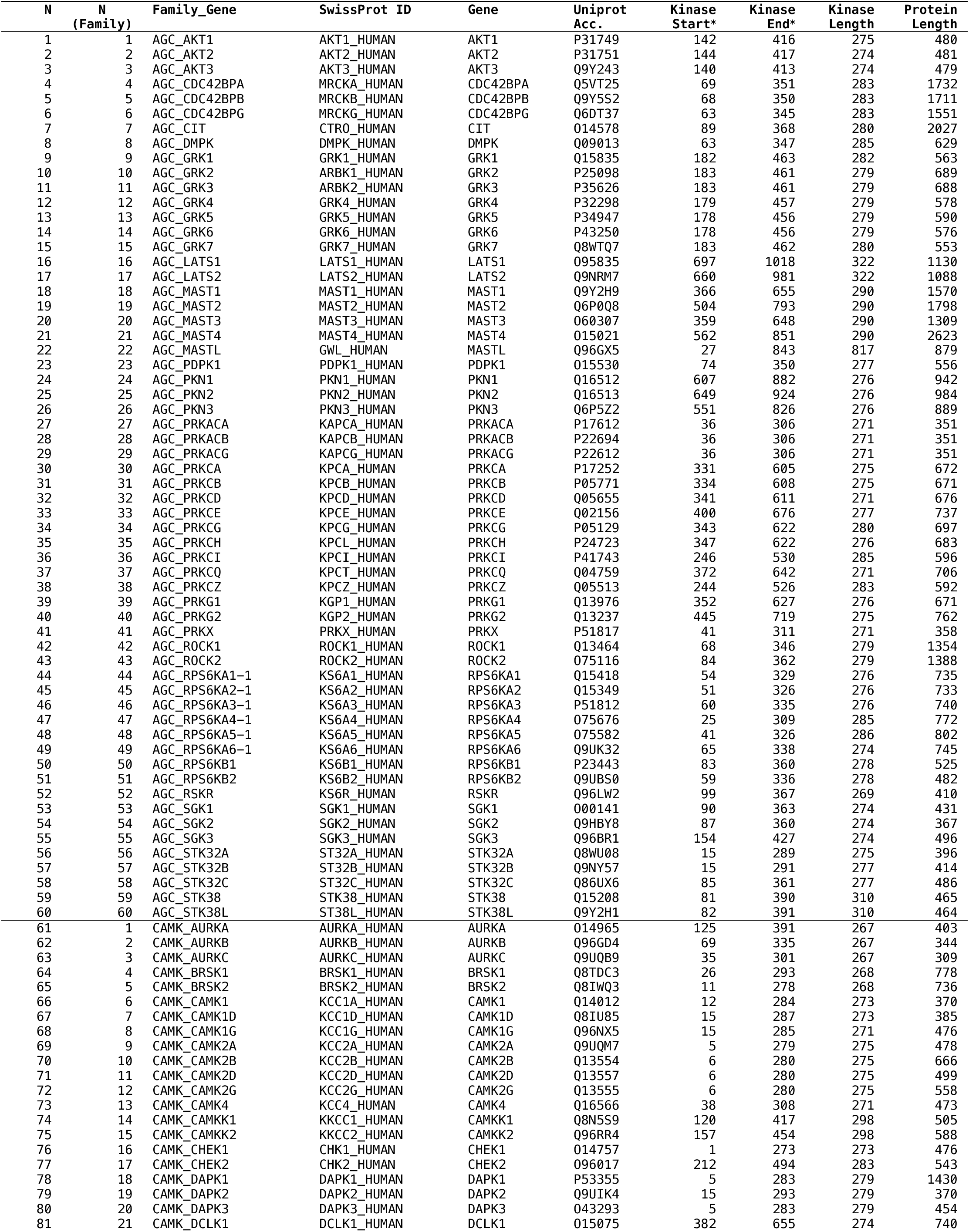
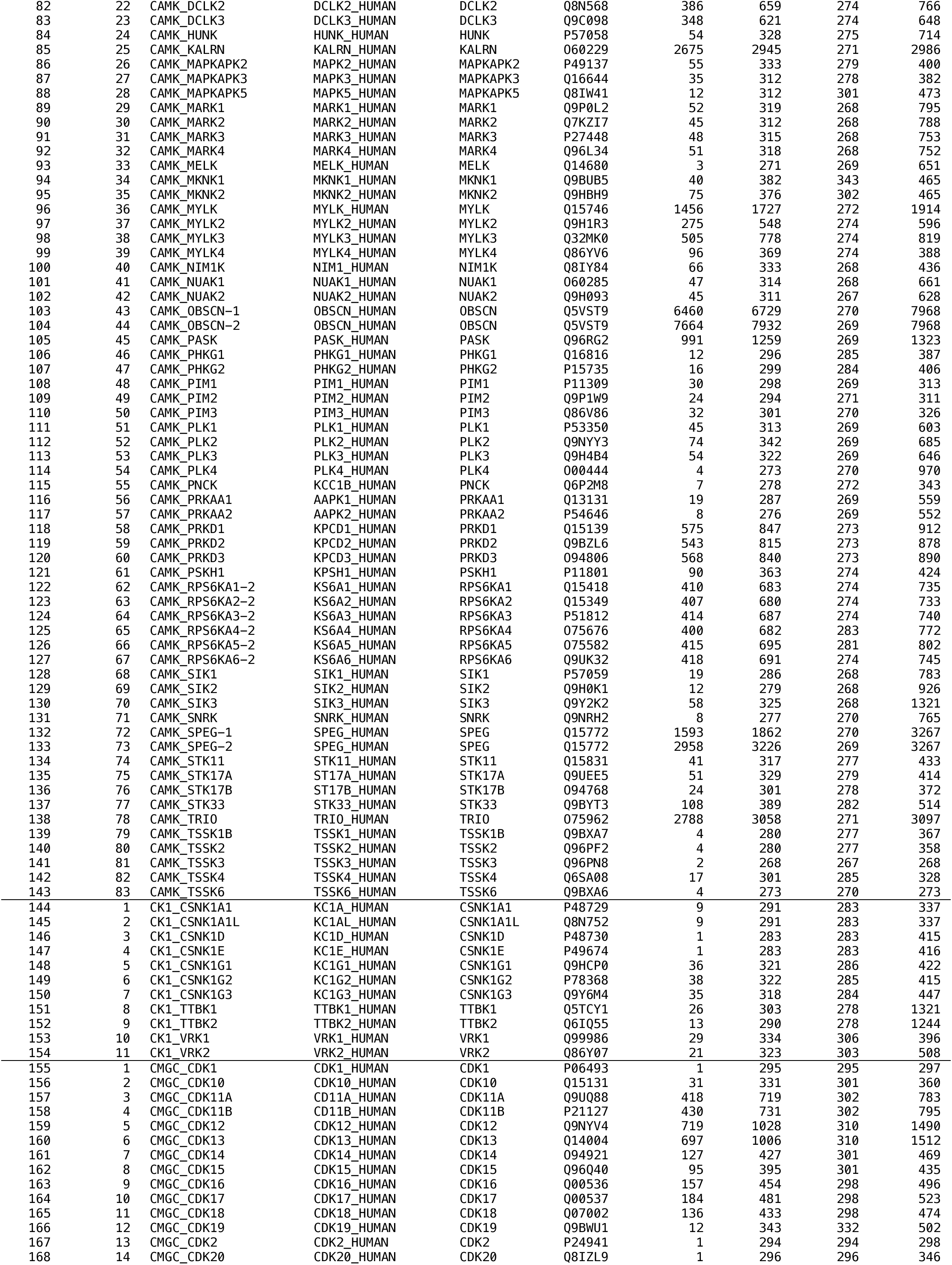
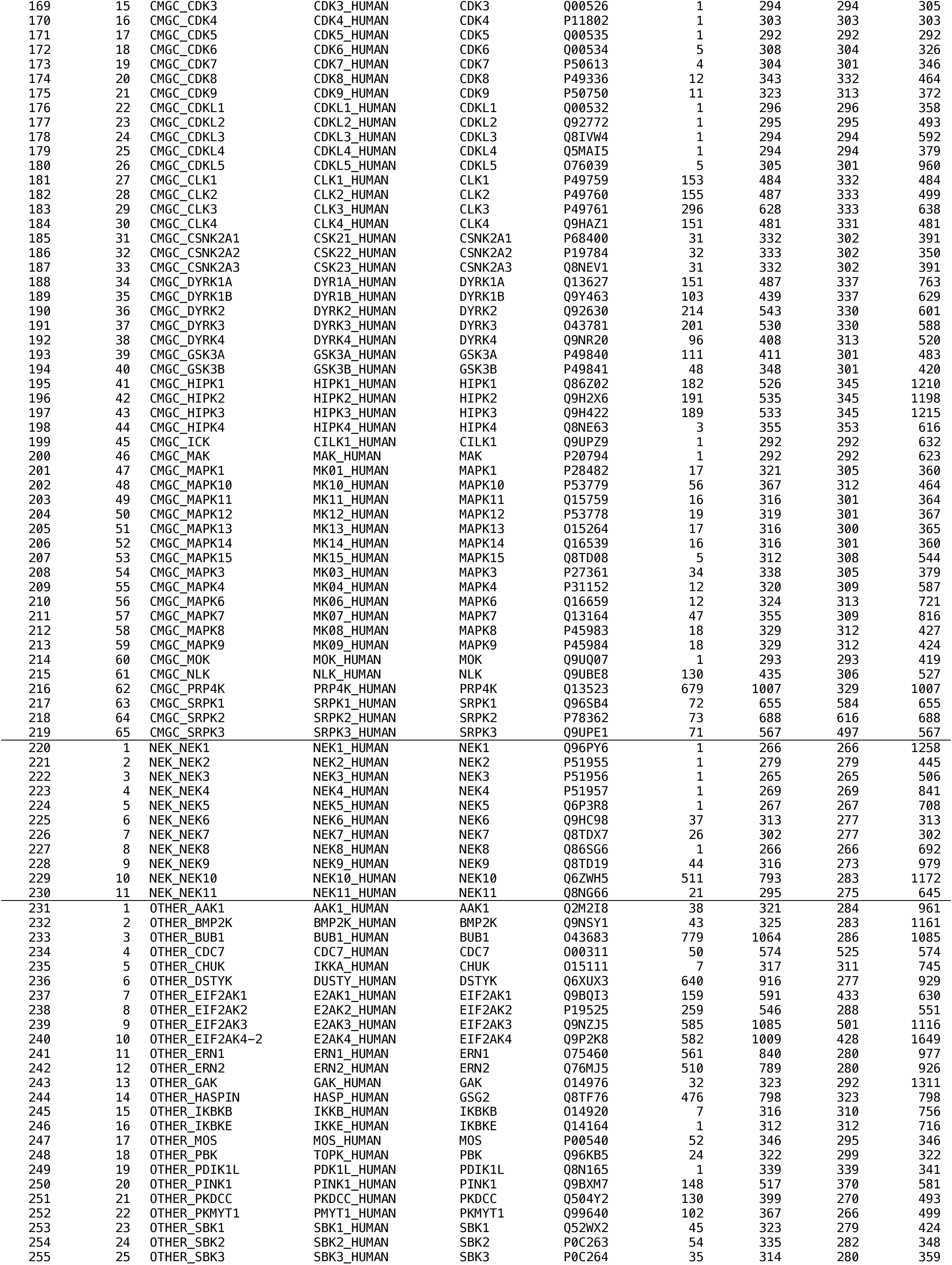
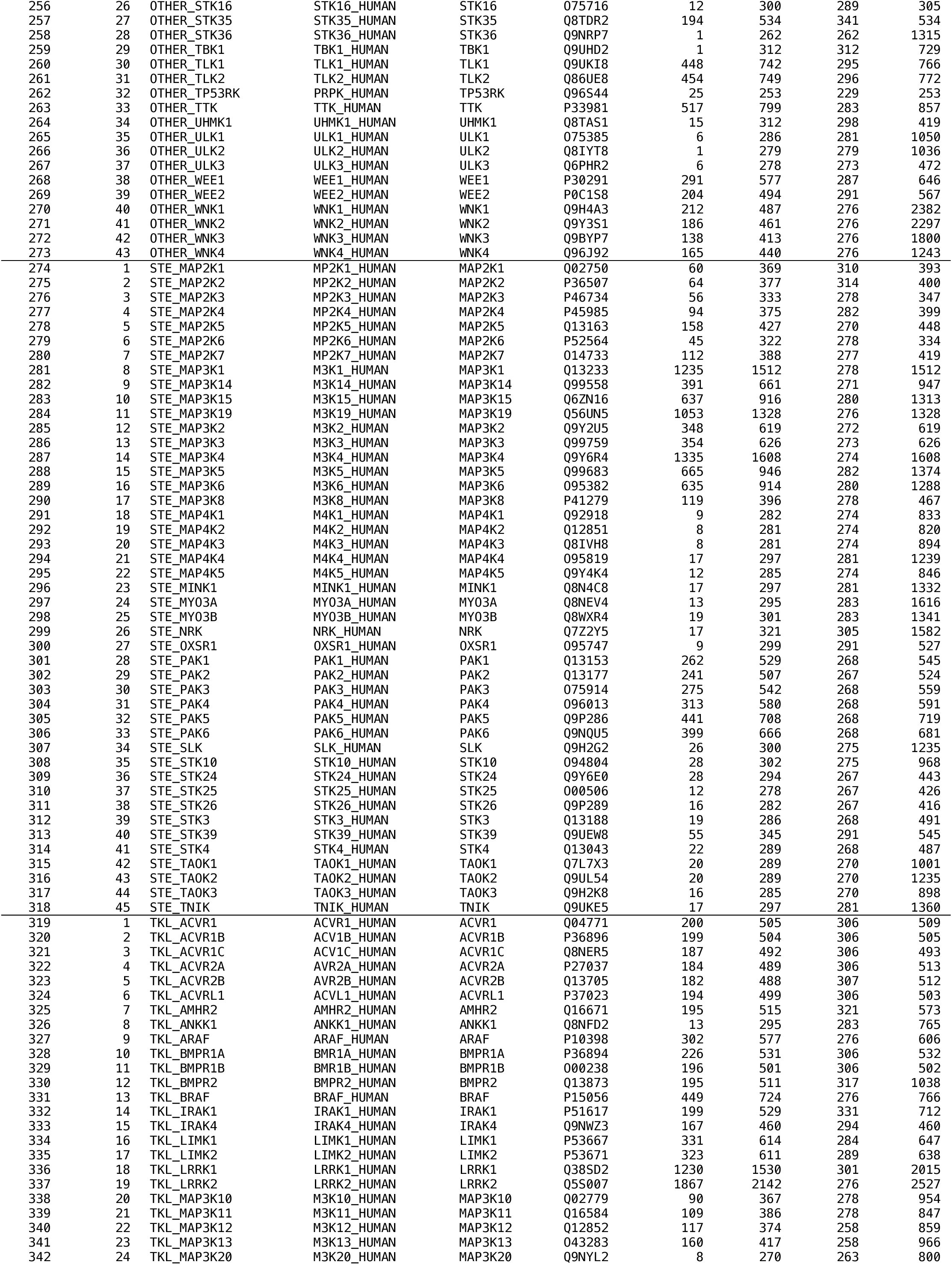
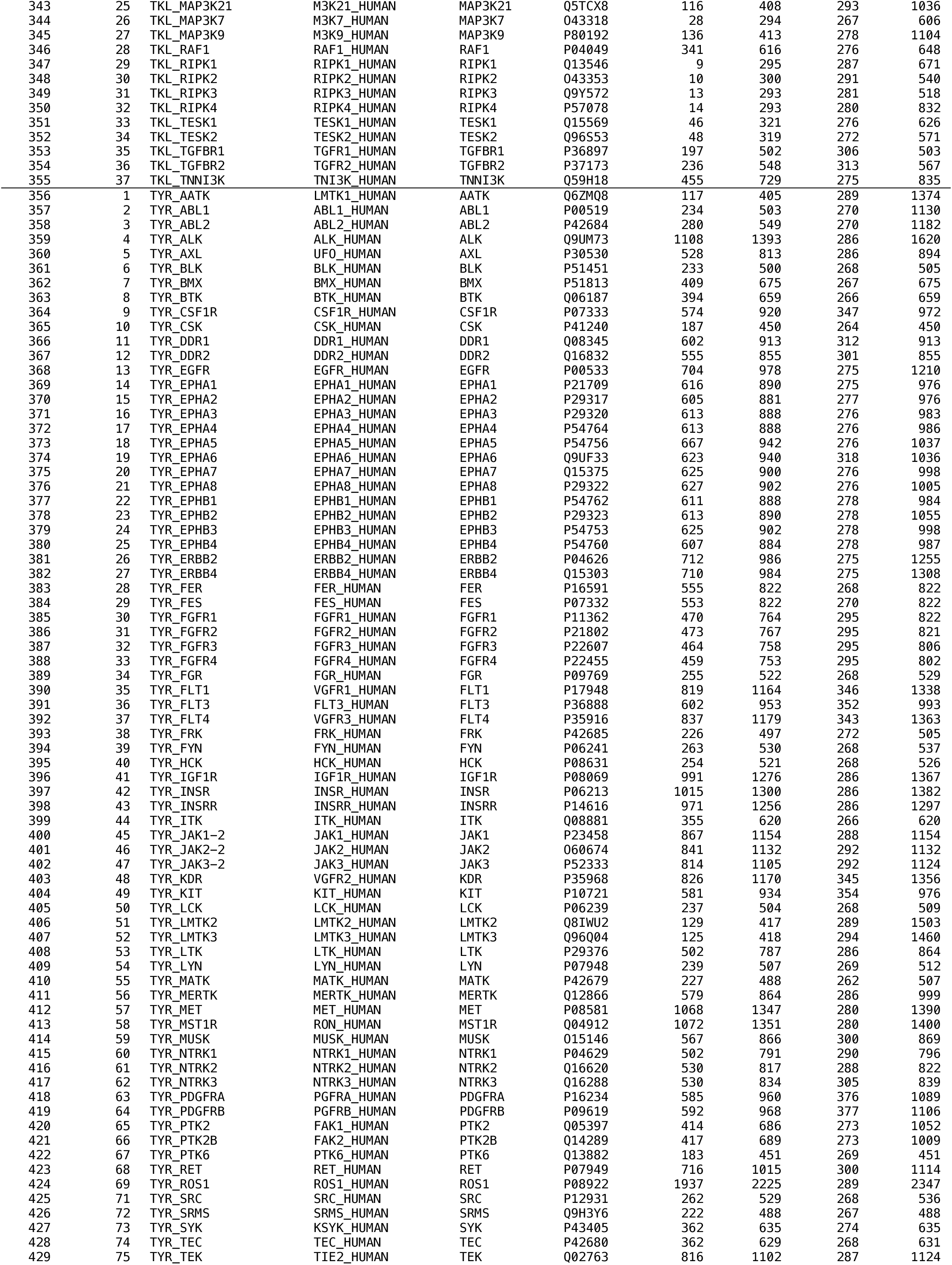
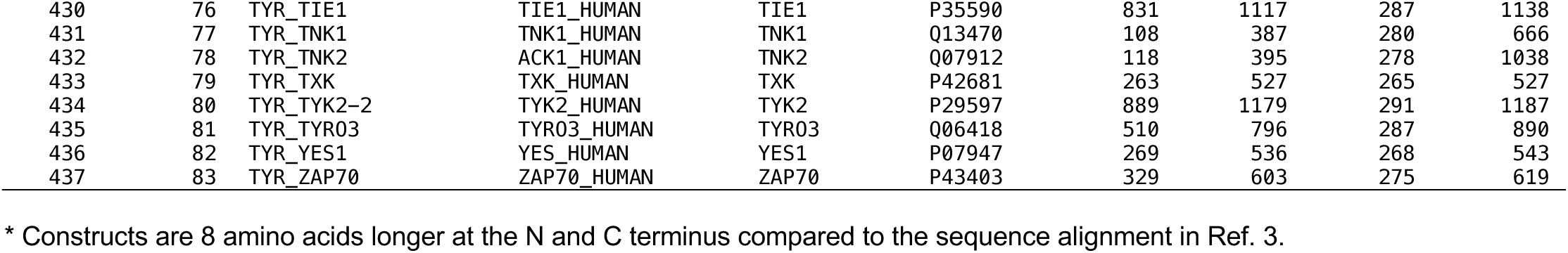
Catalytic kinase domains in the human proteome. The sequence constructs for kinase models produced in this study are given for each human catalytic protein kinase.

**Supplementary Table 2.** Pseudokinase domains in the human proteome.

| N | N<br>(Family) | Family_Gene | SwissProt ID | Gene | Uniprot<br>Acc. | Kinase<br>Start | Kinase<br>End | Kinase<br>Length | Protein<br>Length |
| --- | --- | --- | --- | --- | --- | --- | --- | --- | --- |
| 1 | 1 | CAMK_CAMKV | CAMKV_HUMAN | CAMKV | Q8NCB2 | 16 | 294 | 279 | 501 |
| 2 | 2 | CAMK_CASK | CSKP_HUMAN | CASK | 014936 | 4 | 284 | 281 | 926 |
| 3 | 3 | CAMK_PSKH2 | KPSH2_HUMAN | PSKH2 | Q96QS6 | 55 | 328 | 274 | 385 |
| 4 | 4 | CAMK_STK40 | STK40_HUMAN | STK40 | Q8N2I9 | 27 | 338 | 312 | 435 |
| 5 | 5 | CAMK_TRIB1 | TRIB1_HUMAN | TRIB1 | Q96RU8 | 86 | 346 | 261 | 372 |
| 6 | 6 | CAMK_TRIB2 | TRIB2_HUMAN | TRIB2 | Q925I9 | 56 | 316 | 261 | 343 |
| 7 | 7 | CAMK_TRIB3 | TRIB3_HUMAN | TRIB3 | Q96RU7 | 63 | 323 | 261 | 358 |
| 8 | 8 | CAMK_TTN | TITIN_HUMAN | TTN | Q8WZ42 | 32170 | 32440 | 271 | 34350 |
| 9 | 1 | CK1_VRK3 | VRK3_HUMAN | VRK3 | Q8IV63 | 158 | 463 | 306 | 474 |
| 10 | 1 | OTHER_BUB1B | BUB1B_HUMAN | BUB1B | 060566 | 758 | 1029 | 272 | 1050 |
| 11 | 2 | OTHER EIF2AK4-1 | E2AK4_HUMAN | EIF2AK4 | Q9P2K8 | 272 | 547 | 276 | 1649 |
| 12 | 3 | OTHER_MLKL | MLKL_HUMAN | MLKL | Q8NB16 | 193 | 471 | 279 | 471 |
| 13 | 4 | OTHER_NRBP1 | NRBP_HUMAN | NRBP1 | Q9UHY1 | 56 | 335 | 280 | 535 |
| 14 | 5 | OTHER_NRBP2 | NRBP2_HUMAN | NRBP2 | Q9NSY0 | 29 | 314 | 286 | 501 |
| 15 | 6 | OTHER_PAN3 | PAN3_HUMAN | PAN3 | Q58A45 | 472 | 759 | 288 | 887 |
| 16 | 7 | OTHER_PEAK1 | PEAK1_HUMAN | PEAK1 | Q9H792 | 1319 | 1673 | 355 | 1746 |
| 17 | 8 | OTHER_PEAK3 | PEAK3_HUMAN | PEAK3 | Q6ZS72 | 163 | 405 | 243 | 473 |
| 18 | 9 | OTHER_PIK3R4 | PI3R4_HUMAN | PIK3R4 | Q99570 | 18 | 321 | 304 | 1358 |
| 19 | 10 | OTHER_POMK | SG196_HUMAN | POMK | Q9H5K3 | 73 | 341 | 269 | 350 |
| 20 | 11 | OTHER_PRAG1 | PRAG1_HUMAN | PRAG1 | Q86YV5 | 984 | 1335 | 352 | 1406 |
| 21 | 12 | OTHER_PXK | PXK_HUMAN | PXK | Q7Z7A4 | 138 | 404 | 267 | 578 |
| 22 | 13 | OTHER_RNASEL | RN5A_HUMAN | RNASEL | Q05823 | 353 | 594 | 242 | 741 |
| 23 | 14 | OTHER_RPS6KC1 | KS6C1_HUMAN | RPS6KC1 | Q96S38 | 332 | 1066 | 735 | 1066 |
| 24 | 15 | OTHER_RPS6KL1 | RPKL1_HUMAN | RPS6KL1 | Q9Y6S9 | 145 | 547 | 403 | 549 |
| 25 | 16 | OTHER_SCYL1 | SCYL1_HUMAN | SCYL1 | Q96KG9 | 6 | 271 | 266 | 808 |
| 26 | 17 | OTHER_SCYL2 | SCYL2_HUMAN | SCYL2 | Q6P3W7 | 24 | 335 | 312 | 929 |
| 27 | 18 | OTHER_SCYL3 | PACE1_HUMAN | SCYL3 | Q8IZE3 | 3 | 253 | 251 | 742 |
| 28 | 19 | OTHER_STK31 | STK31_HUMAN | STK31 | Q9BXU1 | 702 | 980 | 279 | 1019 |
| 29 | 20 | OTHER_STKLD1 | STKL1_HUMAN | STKLD1 | Q8NE28 | 20 | 305 | 286 | 680 |
| 30 | 21 | OTHER_TBCK | TBCK_HUMAN | TBCK | Q8TEA7 | 1 | 281 | 281 | 893 |
| 31 | 22 | OTHER_TEX14 | TEX14_HUMAN | TEX14 | Q8IWB6 | 219 | 520 | 302 | 1497 |
| 32 | 23 | OTHER_ULK4 | ULK4_HUMAN | ULK4 | Q96C45 | 1 | 288 | 288 | 1275 |
| 33 | 1 | RGC_GUCY2C | GUC2C_HUMAN | GUCY2C | P25092 | 468 | 767 | 300 | 1073 |
| 34 | 2 | RGC_GUCY2D | GUC2D_HUMAN | GUCY2D | Q02846 | 502 | 815 | 314 | 1103 |
| 35 | 3 | RGC_GUCY2F | GUC2F_HUMAN | GUCY2F | P51841 | 505 | 853 | 349 | 1108 |
| 36 | 4 | RGC_NPR1 | ANPRA_HUMAN | NPR1 | P16066 | 507 | 829 | 323 | 1061 |
| 37 | 5 | RGC_NPR2 | ANPRB_HUMAN | NPR2 | P20594 | 491 | 814 | 324 | 1047 |
| 38 | 1 | STE_STRADA | STRAA_HUMAN | STRADA | Q7RTN6 | 61 | 387 | 327 | 431 |
| 39 | 2 | STE_STRADB | STRAB_HUMAN | STRADB | Q9C0K7 | 50 | 377 | 328 | 418 |
| 40 | 1 | TKL_ILK | ILK_HUMAN | ILK | Q13418 | 185 | 452 | 268 | 452 |
| 41 | 2 | TKL_IRAK2 | IRAK2_HUMAN | IRAK2 | Q43187 | 197 | 509 | 313 | 625 |
| 42 | 3 | TKL_IRAK3 | IRAK3_HUMAN | IRAK3 | Q9Y616 | 152 | 453 | 302 | 596 |
| 43 | 4 | TKL_KSR1 | KSR1_HUMAN | KSR1 | Q8IVT5 | 605 | 887 | 283 | 923 |
| 44 | 5 | TKL_KSR2 | KSR2_HUMAN | KSR2 | Q6VAB6 | 658 | 938 | 281 | 950 |
| 45 | 1 | TYR_EPHA10 | EPHAA_HUMAN | EPHA10 | Q5JZY3 | 637 | 910 | 274 | 1008 |
| 46 | 2 | TYR_EPHB6 | EPHB6_HUMAN | EPHB6 | Q15197 | 662 | 925 | 264 | 1021 |
| 47 | 3 | TYR_ERBB3 | ERBB3_HUMAN | ERBB3 | P21860 | 701 | 975 | 275 | 1342 |
| 48 | 4 | TYR_JAK1-1 | JAK1_HUMAN | JAK1 | P23458 | 575 | 855 | 281 | 1154 |
| 49 | 5 | TYR_JAK2-1 | JAK2_HUMAN | JAK2 | Q60674 | 537 | 815 | 279 | 1132 |
| 50 | 6 | TYR_JAK3-1 | JAK3_HUMAN | JAK3 | P52333 | 513 | 787 | 275 | 1124 |
| 51 | 7 | TYR_PTK7 | PTK7_HUMAN | PTK7 | Q13308 | 788 | 1070 | 283 | 1070 |
| 52 | 8 | TYR_ROR1 | ROR1_HUMAN | ROR1 | Q01973 | 465 | 756 | 292 | 937 |
| 53 | 9 | TYR_ROR2 | ROR2_HUMAN | ROR2 | Q01974 | 465 | 756 | 292 | 943 |
| 54 | 10 | TYR_RYK | RYK_HUMAN | RYK | P34925 | 322 | 606 | 285 | 607 |
| 55 | 11 | TYR_STYK1 | STYK1_HUMAN | STYK1 | Q6J9G0 | 105 | 390 | 286 | 422 |
| 56 | 12 | TYR_TYK2-1 | TYK2_HUMAN | TYK2 | P29597 | 581 | 876 | 296 | 1187 |

Notes: Pseudokinase proteins may have catalytic activity that does not involve protein phosphorylation. The kinase domain in POMK is a protein O-mannose kinase. RNASEL is a 2'-5' endonuclease, in which the pseudokinase domain facilitates homodimerization. The pseudokinase domain of PAN3 participates in mRNA deadenylation by PAN2. The RGC proteins contain active guanylyl cyclase domains; the kinase domains are inactive. Human PLK5 contains a truncated kinase domain at its N-terminus and is not included in the table. The mouse ortholog contains a full kinase domain.

**Supplementary Table 3.**
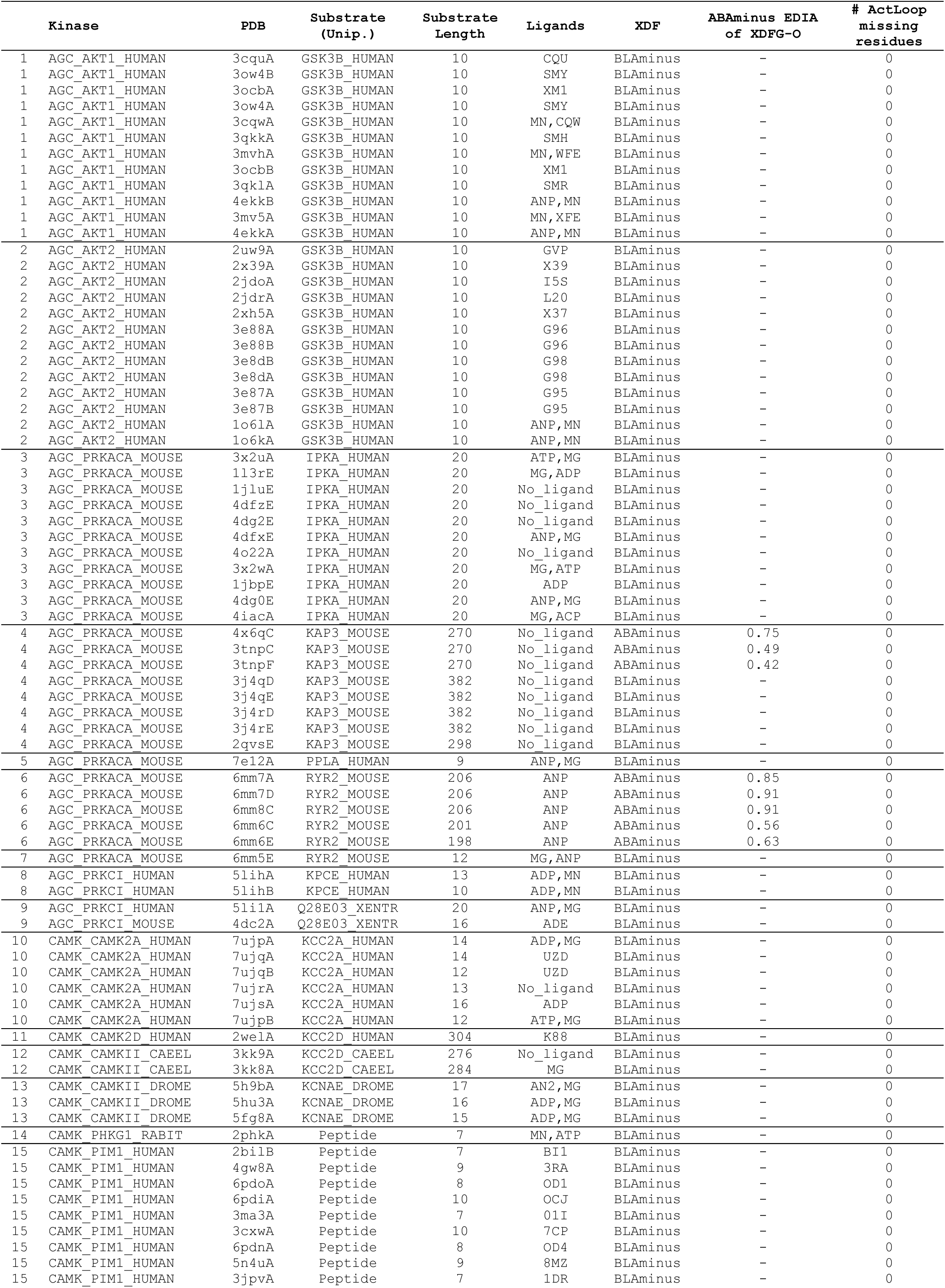
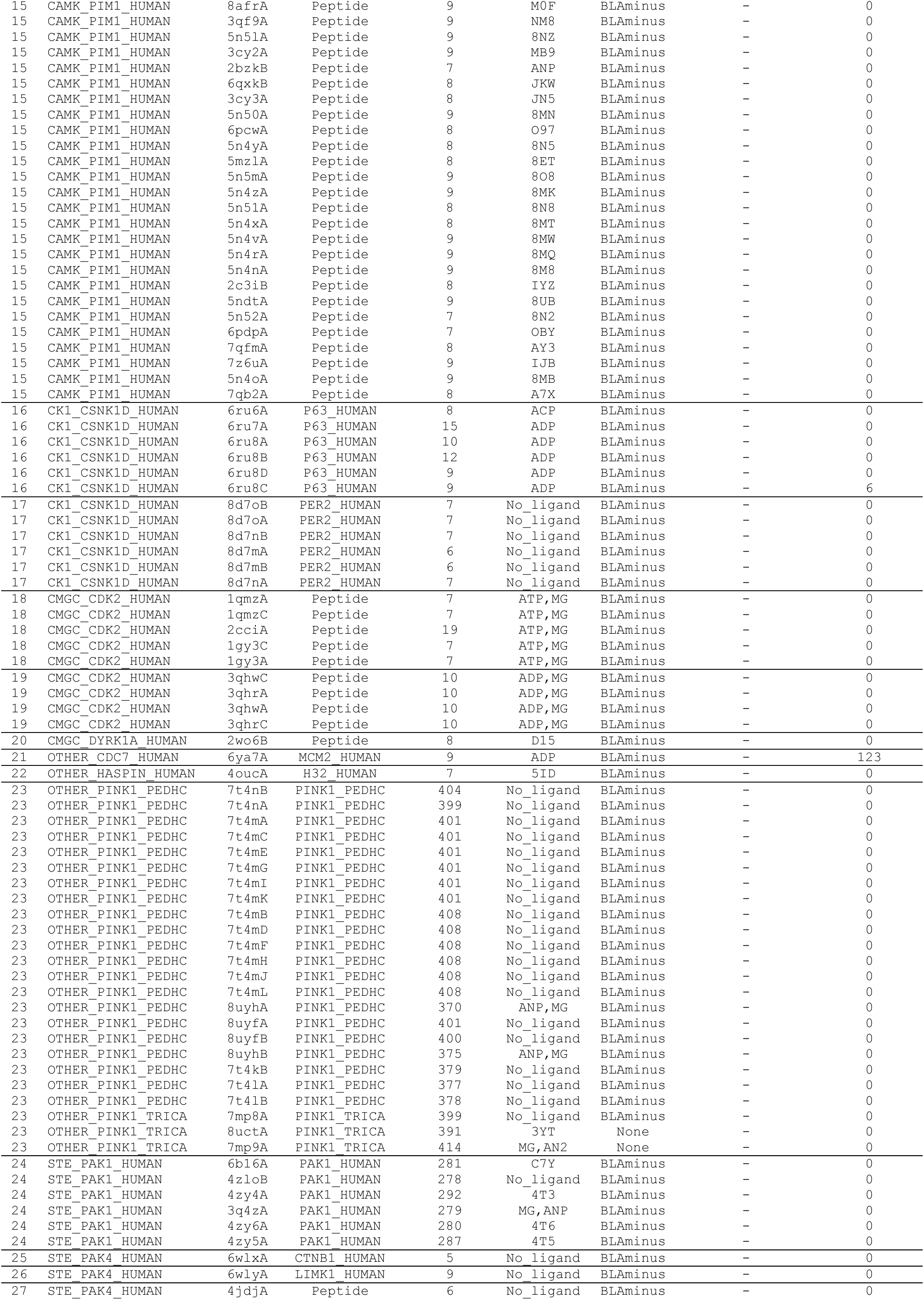
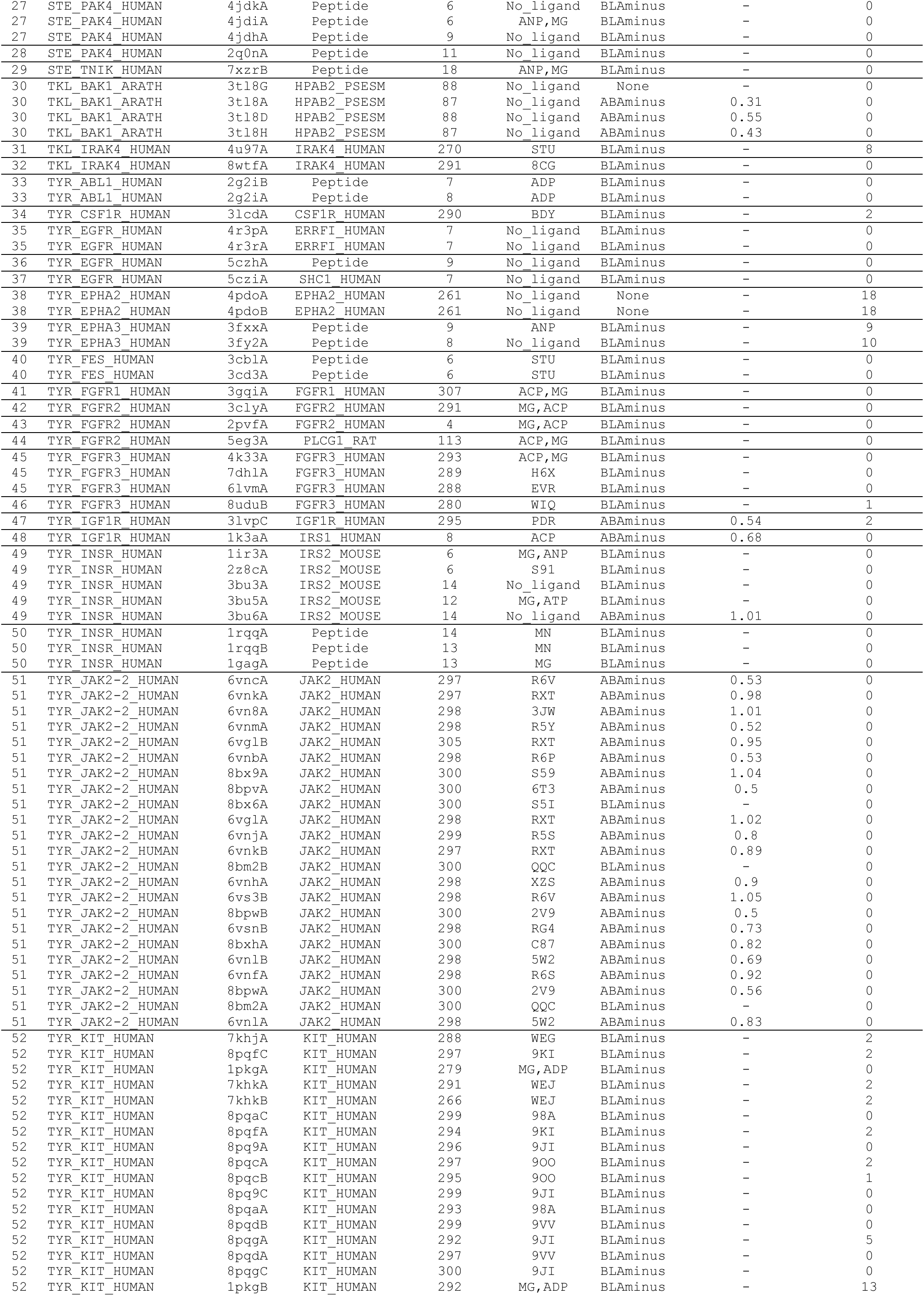
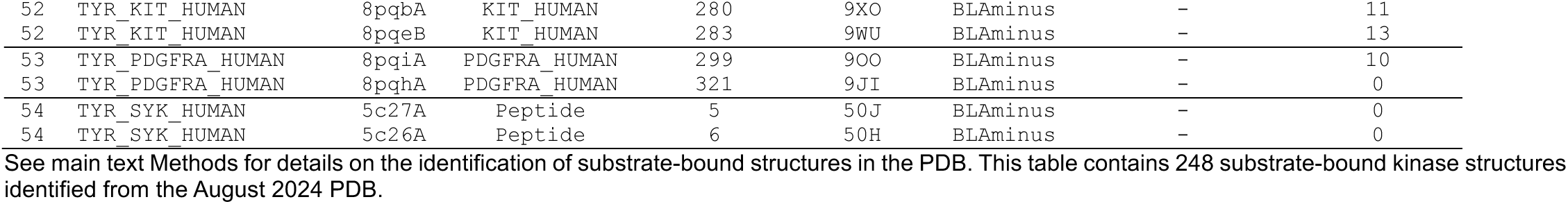
Data for 248 substrate-bound kinase structures.

**Supplementary Table 4.**
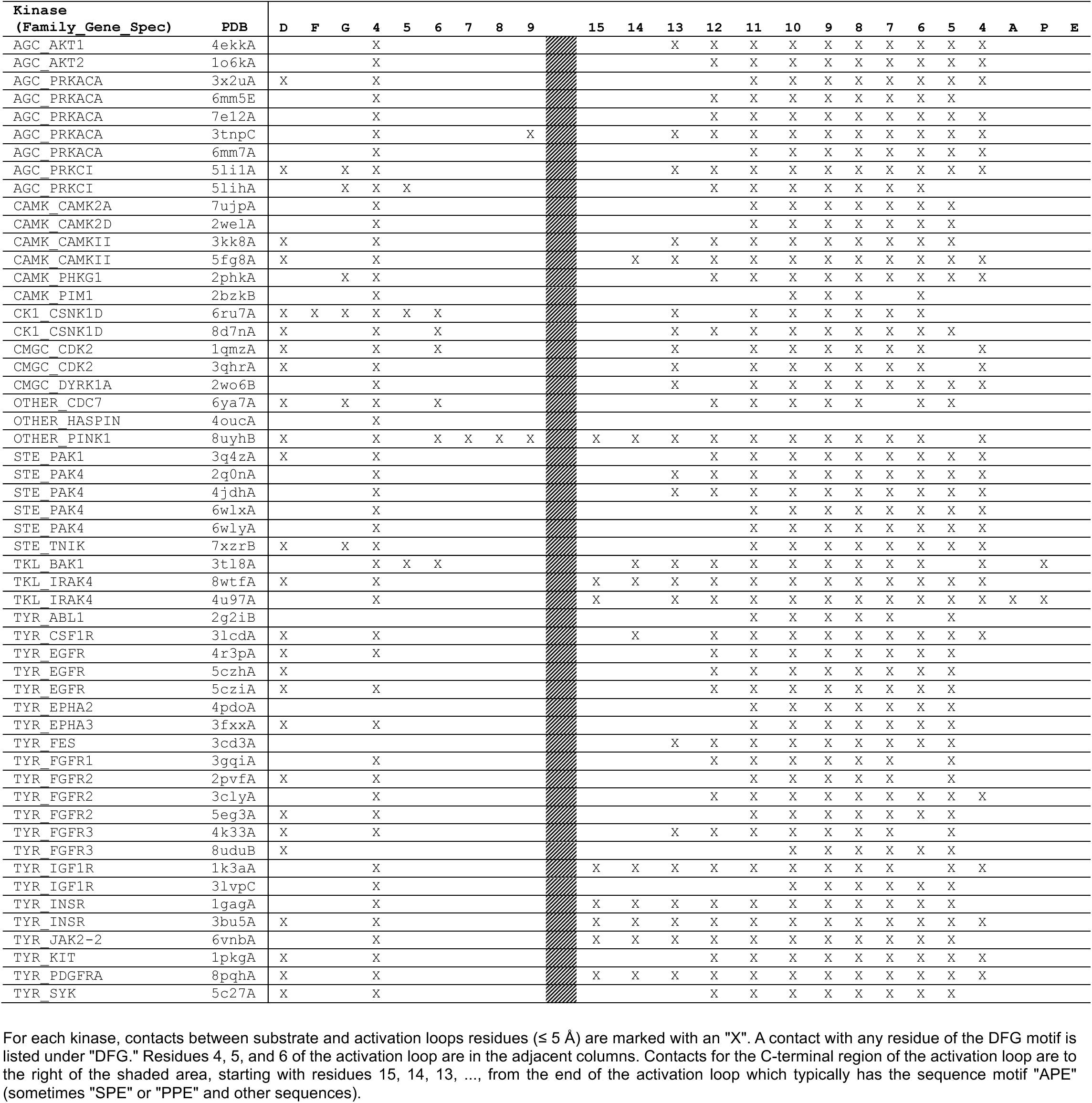
Contacts between activation loop residues and substrate.

**Supplementary Table 5.**
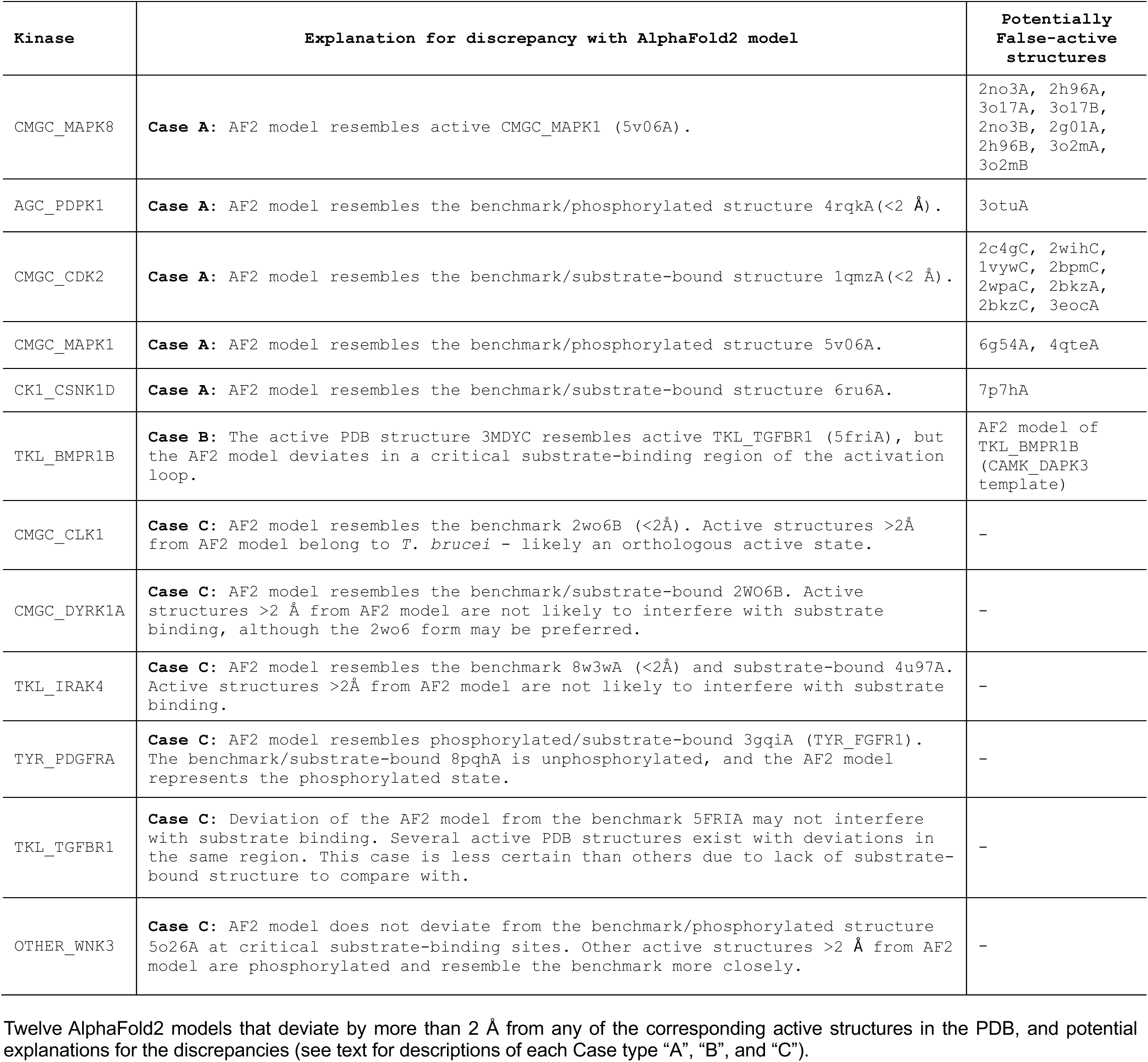
Discrepancies between active-passing PDB structures and AF2 models.

### Supplementary Methods. Pseudocode for Kincore-Standalone3 and for code that produces the Kincore database

1. `Read in list of PDB/mmCIF files to be analyzed`
2. `For each structure:`

a. Extract protein sequence(s) from PDB/mmCIF file for all chains. Sequences are extracted from SEQRES records or _entity_poly.pdbx_seq_one_letter_code records if present. Else they are obtained from the coordinates.
b. `For each chain:`

i. Run hmmsearch of chain sequence against HMMs for each kinase family. HMMs are included for specific kinases that do not align well to the general HMMs (AGC, CAMK, CK1, CMGC, NEK, OTHER, RGC, STE, TKL, TYR), consisting of BUB, EIF2AK41, HASP, MAP3K123, MOS, PAN3, PEAK, PKDCC, PXK, RNASEL, TBCK, TP53RK, ULK, and WNK.
ii. `Determine if the score of the HMM alignment is high enough to be a kinase (>30) or too short <100 residues). Chains which are not kinases are not processed further.`
iii. If the structure is a kinase, identify residues used to determine the conformational state of the structure from the highest scoring HMM. This is performed with the SearchIO function in BioPython. For each HMM, the important residues have been identified by their match-state number in the HMMs. The alignment is used to identify those residues within the kinase sequence/structure. The identified residues consist of: Saltbridge-Lys, Saltbridge-Glu, Glu+4, XHRD residues, XDFG residues, DFG4 and DFG6 residues, APE and APE6-APE12 residues, HPN and HPN7 residues (for the regulatory spine).
iv. `Determine if the kinase chain is a pseudokinase: if the gene name associated with the UniProt identifier (if available) is in a list of 56 human pseudokinase domains (Table S2), then it is labeled “Pseudo.” Otherwise, if the HRD-Asp or DFG-Asp are other residue types, the kinase is labeled “Pseudo.”`
v. Calculate distances required for the conformational classification, including the Saltbridge NZ/OE distance (for SaltBr-in and SaltBr-out), the LysCB-GluCB distance (for Chelix in or out), the PheCZ-LysCA and PheCZ-Glu4CA (for DFGin, DFGinter, DFGout), the XHRD-DFG6 backbone hydrogen bond distances (save the minimum value, for ActLoopNT-in and out), the APE9CA-ArgO distance (for ActLoopCT-in and out), and the APE10-APE12CB/DFG4-CA distances.
vi. `Calculate dihedral angles for the X-D-F residues for the dihedral angle state determination.`
vii. Determine the spatial state: if PheCZ-Glu4CA≤11 Å and PheCZ-LysCA≥11 Å, then the state is DFGin. If PheCZ-Glu4CA≥11 and PheCZ-LysCA<14, then the state is DFGout. If PheCZ-Glu4≤11 and PheCZ-LysCA<11, then the state is DFGinter. If any atoms are missing or the distances do not fall in the above ranges, the spatial state is labeled ‘None’.
viii. Determine the Chelix and salt bridge states: if the LysCB-GluCB distance is ≤ 10.0 Å, then the Chelix state is Chelix-in. Else it is Chelix-out. If the smaller LysNZ-GluOE distance (between OE1 and OE2) is ≤ 3.6 Å, then the structure is SaltBr-in. Else it is SaltBr-out. If atoms are missing, then the states are labeled ‘None’. If the Lys position is not K (as in WNK kinases) or the Glu residue is not E, set the labels to Chelix-na and Saltbr-na (not applicable).
ix. Determine the dihedral angle states by calculating the dihedral angle distances of the backbone ϕ and ψ angles of the X, D, and F residues between the kinase structure and the centroids of each dihedral angle state, given the spatial state. Each distance has a value of D=2(1-cos(a_input-a_centroid)) where a_input is the input structure dihedral angle and a_centroid is the centroid center dihedral angle. If the structure is in the right spatial state (DFGin, DFGinter, DFGout), the average dihedral angle difference over the six backbone angles is < 0.45 and the chi1 dihedral angle of the Phe side chain is in the right rotamer state (minus, plus, trans), then the structure is assigned to the relevant state (DFGin-BLAminus, DFGin-BLAplus, DFGin-BLBminus, DFGin-BLBplus, DFGin-BLBtrans, DFGin-ABAminus, DFGout-BBAminus, DFGinter-BABtrans). If any data are missing, the structure is classified as ‘None.’
x. Determine the ActLoopNT state. If the XHRD-DFG6 backbone-backbone hydrogen bond distance is ≤3.6 Å, then the structure is ActLoop-in. Else it is ActLoopNT-out. If any atoms are missing, this state is labeled ‘None.’
xi. `Determine the ActLoopCT state.`

1. `Calculate the backbone dihedrals of APE6 and APE7 and test for an intact APE5-helix. If APE6 has values ϕ` ∈ `(−180°, 0°) and ψ` ∈ `(−100°, 50°), and if APE7 has values ϕ` ∈ `(−180°, 0°) and ψ` ∈ `(−100°, 50°), the APE5-helix is considered intact. The APE5-helix is also considered intact if a flip of the APE7-APE6 peptide plane is encountered, indicated by APE6 values ϕ` ∈ `(0°, 180°) and ψ` ∈ `(−50°, 100°) and APE7 values ϕ` ∈ `(−180°, 0°) and ψ` ∈ `(50°, 180°).`
2. `Calculate the backbone and χ1 sidechain dihedral angles of the APE8 residue. Active structures are considered with backbone values ϕ` ∈ `(−180°, 0°), ψ` ∈ (50°, 180°). For non-TYR kinases to be considered active, the conserved Thr[APE8] and Ser[APE8] residues must adopt the *g-*sidechain conformation where χ1 ∈ (−120°, 0°) (or 240° to 360° in the shifted coordinate frame computed by adding +360° to the value of χ1 while respecting periodic boundaries at 0° and 360°). If any of these conditions are not satisfied the structure is labeled ActLoopCT-out.
3. `Measure the APE9CA-ArgO distance. If ≤ 6.0 Å for non-TYR kinases and ≤ 8.0 for TYR kinases, then the structure is ActLoopCT-in. Else it is ActLoopCT-out.`
4. `For TYR kinases, also check that the APE9 and APE10 backbone dihedrals have values ϕ` ∈ `(−180°, 0°) and ψ` ∈ `(50°, 180°), else the structure is ActLoopCT-out.`
5. `If the structure is of a non-TYR kinase, measure the APE10-Cβ/DFG4-Cα distance and check if < 8 Å, else it is ActLoopCT-out.`
6. `If the structure is of a non-TYR kinase, measure the APE11-Cβ/DFG4-Cα distance and check if it is in the range (8 Å, 14 Å), else it is ActLoopCT-out.`
7. `If the structure is of a non-TYR kinase, measure the APE12-Cβ/DFG4-Cα distance and check if it is in the range (7 Å, 14 Å), else it is ActLoopCT-out.`
8. `If the kinase is closest to the PKDCC, TP53RK, or HASP HMMs, then set the label to ActLoopCT-na (not applicable).`
xii. `Determine the HRD state by calculating backbone dihedrals of the first and second residues of the HRD motif (HRD1 and HRD2). For HRD1 check if ϕ` ∈ `(−180°, 0°) and ψ` ∈ `(−100°, 50°). For HRD2 check if ϕ` ∈ `(0°, 180°) and ψ` ∈ `(−50°, 100°). Else it is HRD-out. If the F-helix Asp is not present in the sequence (DWW motif), then set HRD_label to “HRD-na” (not applicable).`
xiii. `Determine the activity label of the kinase domain, if the kinase has not been labeled “Pseudo”:`

1. `Label the domain as Active.`
2. `If the spatial_label is not DFGin, then set to Inactive.`
3. `If the dihedral_label is not BLAminus, then set to Inactive, unless the assigned HMM is PKDCC, then set to Inactive if the dihedral_label is not ABAminus.`
4. `If the HRD_label is HRD-out or HRD-na, set to Inactive.`
5. `If neither the Chelix_label or Saltbr_label is “in” or “na”, then set to Inactive.`
6. `If the current label is Active and the Saltbr_label is “none”, set to None.`
7. `If the current label is Active or None, and the Saltbr_label is “out”, set to Inactive.`
8. `If the current label is Active and the ActLoopNT_label is “none”, set to None.`
9. `If the current label is Active or None, and the ActLoopNT_label is “out”, set to Inactive.`
10. `If the current label is Active and the ActLoopCT_label is “none”, set to None.`
11. `If the current label is Active or None, and the ActLoopCT_label is “out”, set to Inactive.`
xiv. `Output the results.`

## Notes

https://dunbrack.fccc.edu/kincore

### Competing Interest Statement

The authors have declared no competing interest.

### Summary of Updates

Spelling error in title of paper (Kinases Domains --< Kinase Domains)

